# Whole genomes of the amazonian *Cacajao* reveal complex connectivity and fast differentiation driven by high environmental dynamism

**DOI:** 10.1101/2023.07.22.550156

**Authors:** Núria Hermosilla-Albala, Felipe Ennes Silva, Sebastián Cuadros-Espinoza, Claudia Fontsere, Alejandro Valenzuela-Seba, Harvinder Pawar, Marta Gut, Joanna L. Kelley, Sandra Ruibal-Puertas, Pol Alentorn-Moron, Armida Faella, Esther Lizano, Izeni Farias, Tomas Hrbek, Joao Valsecchi, Ivo G. Gut, Jeffrey Rogers, Kyle Kai-How Farh, Lukas F. K. Kuderna, Tomas Marques-Bonet, Jean P. Boubli

**Author notes:** These authors contributed equally (Núria Hermosilla-Albala, Felipe Ennes Silva). These authors contributed equally (Tomas Marques-Bonet, Jean P. Boubli).

## Abstract

Despite showing the greatest primate diversity on the planet, genomic studies on Amazonian primates show very little representation in the literature. With 48 geolocalized high coverage whole genomes from wild uakari monkeys, we present the first population-level study on platyrrhines using whole genome data. In a very restricted range of the Amazon rainforest, eight uakari species (*Cacajao* genus) have been described and categorized into bald and black uakaris, based on phenotypic and ecological differences. Despite a slight habitat overlap, we show that posterior to their split 0.92 Mya, bald and black uakaris have remained independent, without gene flow. Nowadays, these two groups present distinct genetic diversity and group-specific variation linked to pathogens. We propose differing hydrology patterns and effectiveness of geographic barriers have modulated the intra-group connectivity and structure of uakari populations. Beyond increasing their representation, with this work we explored the effects of the Amazon rainforest’s dynamism on platyrrhine species.

## Introduction

Recently, there has been an increased effort to describe wild mammalian populations ^1^ from an evolutionary and conservation perspective ^2–6^. This has been particularly noteworthy in the highly endangered Primate order ^7,8^, for which the Amazon has the greatest diversity on the planet. Surprisingly, nearly 20% of Amazonian primate species have been described recently, since 1990 ^9^, evidencing the scarcity of knowledge on this group. For the first time, whole genomes have been generated for many long-neglected and not yet investigated primate species ^8,10–13^. In particular, platyrrhine genomes (those of primates found in Central and Southern America) remain largely unexplored compared to other primate groups, likely due to their phylogenetic distance to the human species and the difficulty of obtaining biomaterials ^14,15^.

In this study, we focus on *Cacajao* or uakari monkeys of the Pitheciidae ^16^, which remains one of the least known primate genera in the Americas. Endemic to the Amazon, uakaris are part of the rich yet poorly known local biodiversity found in it. Their geographic distribution is the most restricted of the three Pitheciinae genera (*Cacajao*, *Chiropotes*, and *Pithecia*), occurring between the Negro–Branco and Ucayali–Solimões–Juruá Rivers ^17^. Eight species have been described in the *Cacajao* genus, which in turn are clustered into two groups, the bald and the black uakaris ^16^. They show contrasting phenotypic (pelage coloration ^16,20^ and skin pigmentation ^18^), ecological (habitat preference ^19^, geographic distribution ^19,20^) and genomic traits ^19,21^. The bald uakaris comprise five species (*C. calvus*, *C. amuna*, *C. novaesi*, *C. rubicundus*, *C. ucayalii*) ^20^ and black uakaris, three (*C. ayresi*, *C. hosomi*, *C. melanocephalus*) ^22^. Black uakaris are, as their name suggests, black haired and in some cases partially brown ^16^. They are mostly found in the forests of Solimões-Japurá and Negro-Branco river systems, and inhabit extensive areas of *terra firme* forest during some months, including high altitude areas ^17^. On the other hand, bald uakaris are mainly flooded-forest specialists found around white water rivers (*várzea*) and present a red bald face with pelage that ranges from very light-greyish white (white bald uakaris) to red-chestnut (red bald uakaris) ^18,20^. The populations of the five species of bald uakaris are found in the Ucayali, Solimões, and Juruá river systems, nearly overlapping but in general south of the black uakaris’ distribution range ^23^ (Figure 1A).

**Figure 1.**
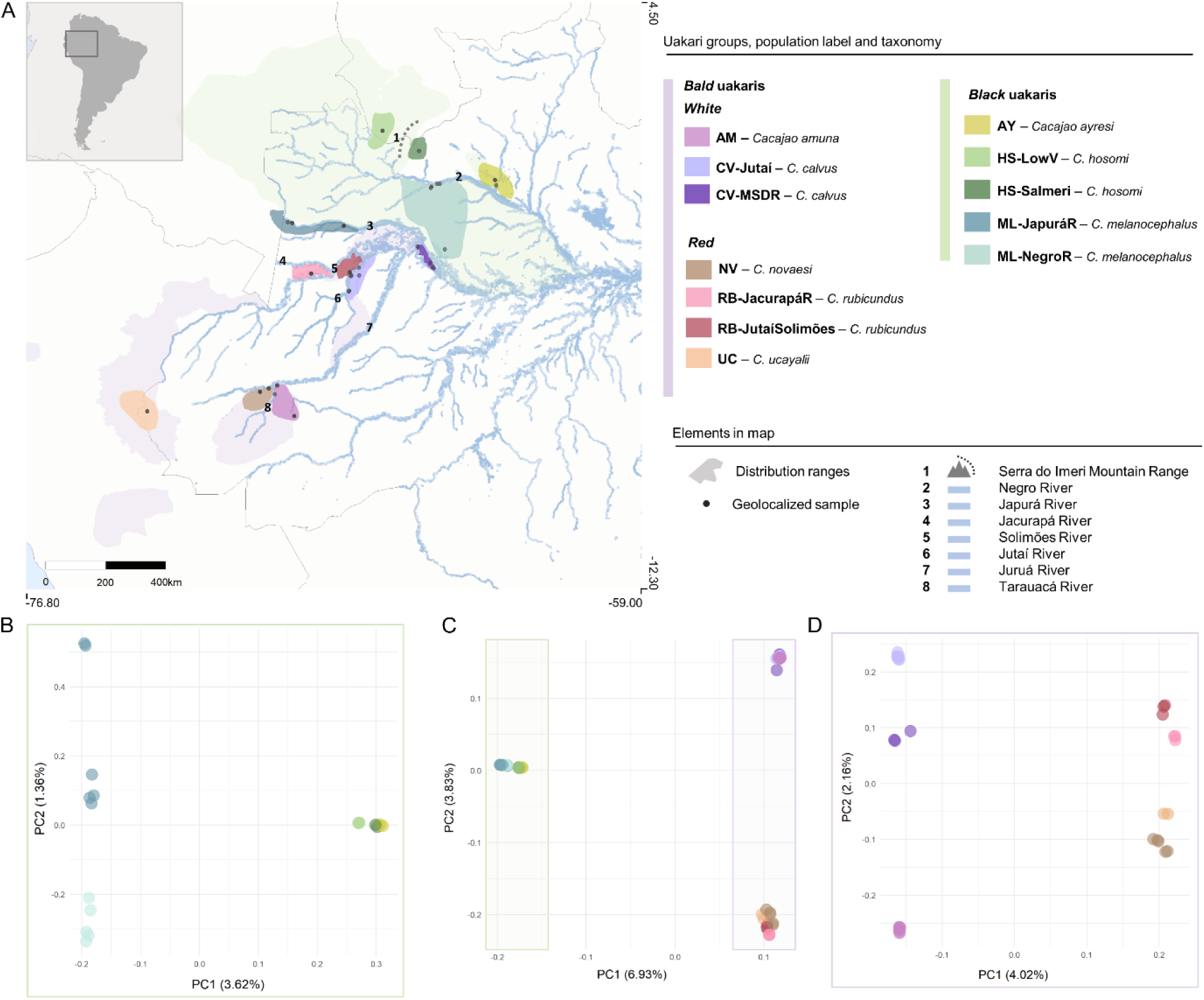
Sampled population distributions and population structure. Overall geographic distribution and structure of *Cacajao* wild populations. **A)** Samples localities and population geographic distributions. **B)** Principal component analysis on the black uakari dataset (n = 2.24M SNPs). **C)** Principal component analysis on the global dataset (n = 10.46M SNPs). **D)** Principal component analysis on the bald uakari dataset (n = 6.10M SNPs).

Complex demographic histories (recent divergence times, incomplete lineage sorting, complex gene flow patterns), combined with limited resolution of previous studies ^19,20,24–26^ have resulted in an incomplete understanding of the evolutionary history of many platyrrhine lineages. This includes the uakaris ^12,19,20,27^, which show shallow divergence times and rapid radiation ^19,20,24^. Multiple studies have addressed the evolutionary history of *Cacajao* employing mitochondrial and double digest restriction-site (ddRAD) associated sequences, whose resolution was insufficient to fully resolve their evolutionary history ^19,20,24^. In accordance with the described phenotypes, genetic data in these studies has suggested that bald and black uakaris form two differentiated phylogenetic clades ^19^ which in turn can be split into red and white bald uakaris, and north and south bank of the Negro river populations for black uakari species ^19,20,24^. Nonetheless, neither the detailed population structure nor gene flow patterns among the populations of wild uakaris have been described, population-level phylogenetic resolution has not been reached and the reported divergence time estimates inside the genus have not been consistent among studies ^19,20,24^. Here we present the first population-level study on whole genome data of a platyrrhine genus. Our dataset consists of 48 geolocalized uakari whole genomes at ~30X coverage, with which we describe the detailed dynamics and population structure of wild *Cacajao* populations with increased statistical power ^15,28^. This is also key to refine previous phylogenetic and demographic estimates ^6,15,28,29^: we obtained the first whole genome phylogeny for uakaris with population-level resolution, narrower or new confidence intervals for divergence times and effective population size (*Ne*) estimates in the genus. Our work provides insights for future efforts to protect *Cacajao* wild populations, whose habitat integrity is becoming increasingly vulnerable in the face of global climate change and anthropogenic threats ^30^.

## Results

Here, we generated 48 whole genomes at a mean coverage of 31x ranging from 14x to 43x (Supplementary Table S1) in which we discovered a total of 191,4 million single nucleotide polymorphisms (SNPs). Twelve populations from all described *Cacajao* species were analyzed here, which were defined based on taxonomy and sampling localities (Supplementary Table S3).

### Population structure and phylogenetics follow pelage coloration and agree with intra-group geographic distribution patterns

To determine the structure of current wild *Cacajao* populations (Figure 1A), we ran a principal component analysis (PCA) (Figure 1B-1D), ADMIXTURE (Figure S4-S6) and phylogenetic analyses (Figure 2). PCA and ADMIXTURE analyses consistently show that population structure in the genus follows patterns described by pelage coloration: the largest genetic dissimilarity being found between bald and black uakari populations (variance explained by PC1: 6.93%; Figure 1C, S4A), followed by differences within each group which were addressed independently (bald: red bald uakaris - white bald uakaris with a variance explained of 4.02% by PC1; black: ML (*C. melanocephalus*) *-* AY (*C. ayresi*) + HS (*C. hosomi*) with PC1 accounting for 3.62% of the variance) (Figure 1B, 1D, S5A, S6A) overall in the absence of admixed individuals (Figure S4A, S5A, S6A). Strong intra-species population structure was identified in white CV (*C. calvus*) and red RB (*C. rubicundus*) bald uakari populations (Figure 1D), as well as for ML in black uakaris (Figure 1B). In the latter group, AY and HS consistently cluster together (Figure 1B, 1C). To further investigate the identified genomic clusters, *F_ST_* was estimated as a proxy for genetic differentiation between populations. As expected, the highest differences in the genus were between bald and a black populations: the red bald RB-JutaíSolimões population and both black HS-SaImeri and AY (*F_ST_*=0.546) (Figure S11). Then, AY and ML-JapuráR were the most differentiated intragroup pair with an *F_ST_*of 0.247, close to that between CV-Jutaí and RB-JutaíSolimões (*F_ST_*=0.233) (Figure S11). Overall, we observed that the geographic distribution of the samples aligned with the intra-group genomic clusters, and even at the species level, where subpopulations could be identified both geographically and at the genomic level in some cases (Figure 1).

**Figure 2.**
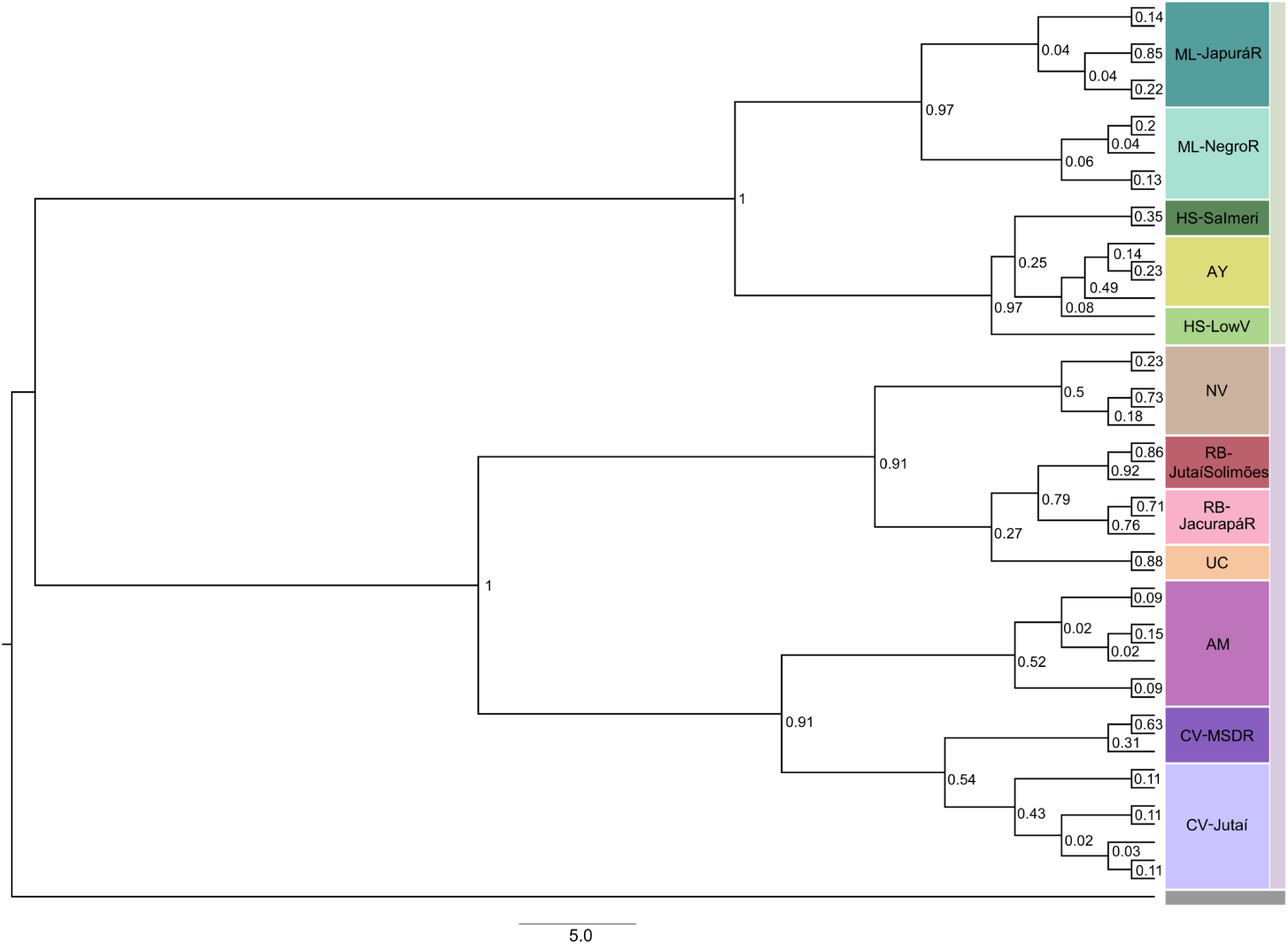
Whole genome phylogeny phylogram. Whole genome Bayesian phylogeny from 2144 maximum likelihood trees based on 1Mb sliding windows. Posterior probability indicated in each split. Phylogram where branch lengths are proportional to the observed phylogenetic distance in terms of substitutions. Colors depict labeling system, outer bar indicates bald (plum) or black (green) uakari group and bottom gray bar indicates outgroup node (*Pithecia pithecia*).

Moreover, the phylogenetic signal in the *Cacajao* lineage was explored by generating the first set of whole autosomal genome Bayesian phylogenies. Two independent phylogenies were generated to assess the influence of variable window sizes on the final posterior probability of the respective topologies. We built a first phylogeny based on 2144 1Mb maximum likelihood (ML) trees covering ~80% of the genome (Figure 2), and a second phylogeny based on 9313 250kb ML trees covering ~85% of the genome (Figure S8A, S8B). The node support in the phylogeny based on 1Mb windows appeared overall higher (Figure 2, S9A) when compared to the second phylogeny based on 250kb windows (Figure S8). Furthermore, the phylogenetic signal retrieved was observed to be homogeneously distributed through the genome rather than driven by specific regions (Figure S9B, S9C). The topologies described here were in accordance with previous phylogenies built from restricted autosomal regions (ddRAD sequences) ^20^, although a more precise reconstruction was achieved employing the whole autosome reaching population-level resolution.

Focusing on the highly supported phylogenetic tree (1Mb windows) and in accordance with PCA and ADMIXTURE analyses, black and bald uakaris split following pelage colouration patterns as well as based on geographic distribution. This was with the exception of a very low-supported deviation (Posterior_probability = 0.08) for one sample from HS-LowV (PD_0914), which clusters with AY. This was not surprising, as these populations showed very little genomic differentiation (*F_ST_* _(HS-LowV,AY)_ = 0.04, Figure 1B, 1C). On the other hand, in accordance with previous studies ^20^, the phylogeny built from the respective whole mitochondrial genomes (Figure S10) showed lower resolution: it recovered species-level classification for black uakaris, and bald uakaris were correctly classified into red and white except for one sample from CV-MSDR (PD_0434), which clustered into red uakaris.

Lastly, a major difference in the topology of red bald uakaris arose when the two phylogenies based on the whole autosome were compared. The 250kb windows - based tree showed RB to diverge earlier from the ancestor of UC (*C. ucayalii*) and NV (*C. novaesi*) (Posterior_probability = 0.67, Figure S8). This was in disagreement with the 1Mb windows - based tree (Posterior_probability = 0.91, Figure 2) and previous studies based on ddRAD ^20^, both of which describe the split between RB and UC happened after the split of their common ancestor from NV. This divergence has been estimated to be quite recent (0.26 Mya) ^20^, which may explain why the complete resolution of the phylogenetic relationship inside red bald uakaris appears harder to reach.

### Recent divergence times in *Cacajao* have resulted in different genetic diversity ranges

In order to uncover the past demographic history of wild uakaris, constant effective population sizes (*Ne*) and divergence times were estimated employing a maximum likelihood approach with fastsimcoal2 and three different model topologies (Figure 3A, Supplementary Methods 1.3.1, Supplementary Table S6). Among these, split times were estimated providing a refined and narrower confidence interval in three of the cases when compared to previous ddRAD estimates (Bald-black: 0.92 Mya [0.89-0.98], bald_red-bald_white: 0.66 Mya [0.64-0.68], CV-AM: 0.16 Mya [0.15-0.18]) ^20^, a similar estimate for the divergence time inside black uakaris to previous studies (0.66 Mya [0.27-0.82]) ^20^ and lastly, a new estimate for the divergence between the two ML populations (0.03 Mya [0.02-0.15]).

**Figure 3.**
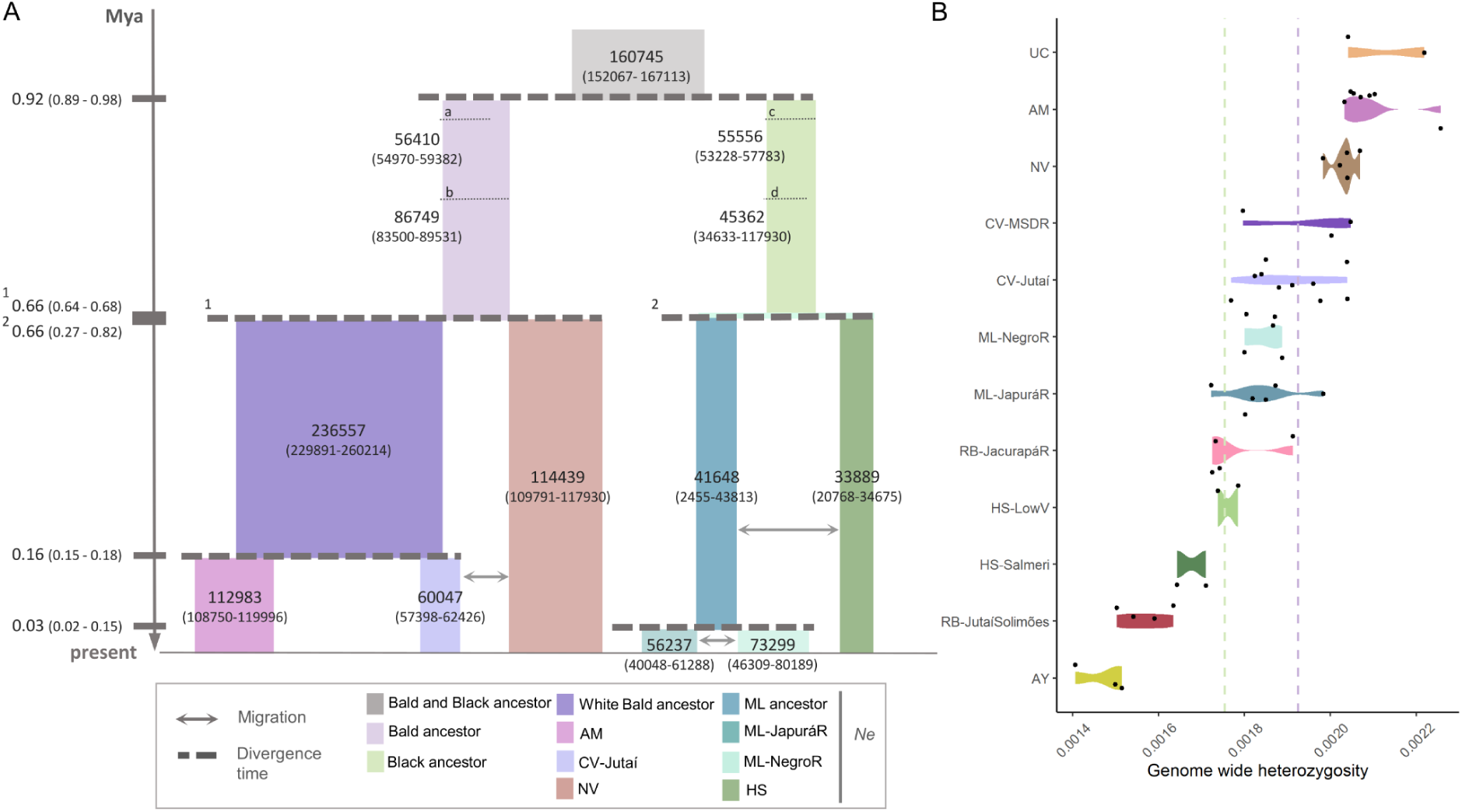
Demographic history of *Cacajao* and genetic diversity in extant populations. **A)** Maximum likelihood model topologies and estimates for divergence times and effective population sizes. Three models summarized in Figure 3A: i) bald and black uakaris general model, ii) bald uakaris model and iii) black uakaris model. Bald and black ancestor *Ne* estimates were calculated twice: in the general model (a,c) and in the bald (b) and black (d) models respectively. **B)** Genome-wide heterozygosity per population, dashed lines indicate mean values per group (bald: plum, black: green).

Focusing on the *Ne* estimates for extant populations, a trend can be observed where bald uakaris tend to have an overall larger *Ne* and mean genome wide heterozygosity estimates when compared to black populations – with the exception of CV-Jutaí, which shows lower *Ne* than the ML-NegroR population, and both RB populations, whose mean heterozygosity falls in the black range. The black AY population presented the lowest genetic diversity among *Cacajao* populations (Figure 3B). Nonetheless, inbreeding was not widespread in the genus and remarkably non-existent in the sampled black uakari populations (Supplementary Table S1). Overall, *Cacajao* showed a lower genetic diversity than other genera in the Pitheciidae in accordance with Kuderna *et al.* (2023) ^8^ (Figure 3B). This ranged between 0.0014 and 0.0025 bp^-1^ (Figure 3B) with variable density distribution in each population (Figure S3).

### Geographic barriers of distinct effectiveness drive independent population dynamics in bald and black uakaris

The dynamics of wild uakari populations were investigated here as a potential underlying force driving the observed genomic clustering in our dataset. No significant signals of gene flow were detected between bald and black uakaris (Figure 4). Population dynamics in black uakaris were shaped by stable allopatric (geographic) barriers (Negro River and Serra do Imeri mountain range, Figure 1A), which determined a discrete genomic structure in the group (Figure 1B, S6B) while not preventing gene flow in some regions (Figure 4). On the other hand, while the genomic structure in bald uakari populations was identifiable and populations were delimited by rivers too, we hypothesized variations in the hydrology in the floodplains of the Amazon basin and related river rearrangements have had an impact on the dynamics of bald uakaris ^21^, given their proposed variable effect as effective barriers to gene flow ^31–34^.

**Figure 4.**
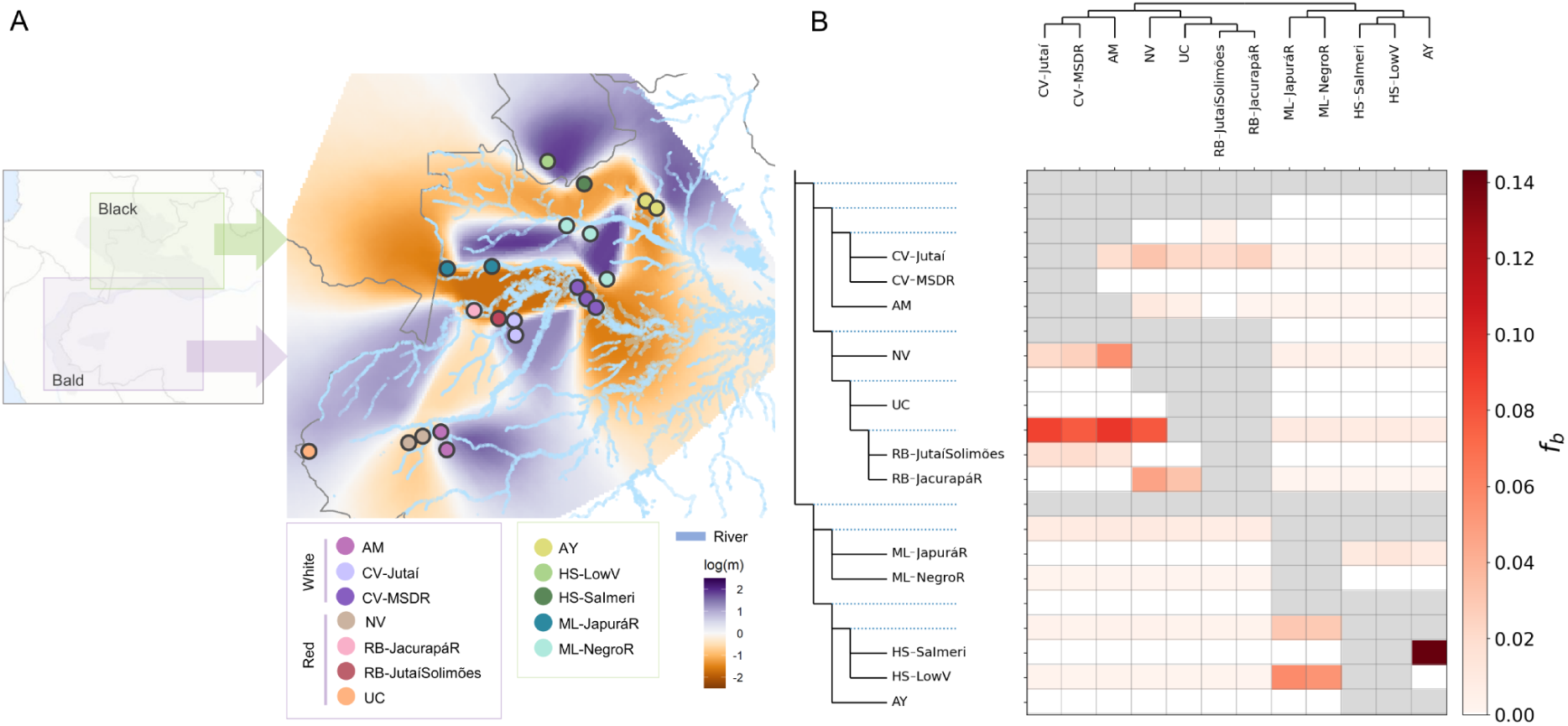
Migration patterns and gene flow in the *Cacajao* genus. **A)** Estimated effective migration surfaces on black and bald uakaris together, color palette depicting effective migration rate values (*log*(*m*)). **B)** F-branch results on full dataset; significant values considered (*f*-branch ≥ 0.05).

#### Bald uakaris

Consistently elevated *f*-branch values (Figure 4B) indicated widespread gene flow between the ancestors of white bald uakaris and the red bald RB. Focusing on CV-Jutaí, this approximated Isolation by Distance (IBD) with RB populations. This was geographically plausible particularly in the case of the RB-Jutaí-Solimões, where the dynamics of the Juruá, Solimões and Jutaí rivers would have affected the connectivity between the two groups ^33,34,32^ found at opposite banks of it (Figure 1A), allowing the approximation to a population following an IBD model (Figure 4).

On the other hand, after the split of white bald uakaris, deviations from IBD appeared: (i) Positive effective migration rates were observed between the two CV populations (*F_ST_ =* 0.067) as shown in Figure 4A, although apart from recurrent interbreeding, strong shared ancestry could intensify such a signal. (ii) CV-MSDR and AM (*C. amuna*) (*F_ST_ =* 0.065) did show connectivity (Figure 4A), and although ADMIXTURE analyses (Figure S5B) showed the first population would share a small ancestry component with AM, we suggest that CV-MSDR represents a unique ancestral component on its own, as this signal was not captured by any other method. (iii) Connectivity was limited between CV-Jutaí and AM (*F_ST_* _(AM,CV-Jutaí)_ *=* 0.092) (Figure 4B).

Gene flow signals were detected between AM and NV (Figure 4B), species currently found on opposite banks of the Tarauacá River (Figure 1A), where no migration was previously detected (Figure 4A).

#### Black uakaris

Negro River - at its widest - together with Serra do Imeri appeared to fragment black uakari populations (Figure 1A, 4A). In agreement with field observations which propose that ML distribution expands beyond Negro River’s headwaters towards Colombia, results suggest that at the north-west of their distribution range, ML had historically interbred with HS from lowland Venezuela (HS-LowV) (Figure 4B) (*F_ST_* _(HS-LowV,ML)_ = 0.019, *F_ST_* _(HS-SaImeri,ML)_ = 0.021). On the other hand, we observed high connectivity between ML populations potentially related to river channels formed in the past between Japurá and Negro River ^35^ (Figure 1A, 4A), although this signal is most likely influenced by their very shallow divergence time (0.03 Mya [0.02-0.15]). Finally, at the northern bank of Negro River, connectivity was found between AY and HS (Figure 4A) (*F_ST_* _(HS-SaImeri,AY)_ = 0.041, *F_ST_* _(HS-LowV,AY)_ = 0.046) although it had been historically more intense between the first and the population of HS-SaImeri (Figure 4B). These results explain the persistent clustering between these species in structure analyses (Figure 1B, S6B, S11). The latter observations were also anticipated by field observations given the absence of clear barriers between HS-SaImeri and AY versus the allopatric barriers between them and the rest of the black populations.

In spite of the high connectivity among HS populations (Figure 4A), these showed very distinct gene flow patterns (Figure 4B). The split between AY and HS was estimated to be as recent as 0.2Mya ^20^, therefore all the events these are involved in - population split in HS, interspecies gene flow - must have been even more recent. Furthermore, the effective migration rates among HS populations were likely overestimated because of their strong shared ancestry. All this would be consistent with the independent gene flow signals shown by these two populations with AY and ML respectively.

### Functional genetic differentiation between groups is led by Malaria and Integrin-related pathways

Given the consistent genomic separation observed between bald and black uakaris (Figure 1, 2, S11), their distinct phenotype and independent population dynamics (Figure 4), we aimed to explore the trace of their genome-wide differentiation at the functional level. For that, we identified those coding regions appearing most divergent in all intergroup comparisons: differentially fixed non-synonymous (NS) variants among bald and black uakaris were filtered by alignability of the given amino acid and conservation levels of these across the primate lineage (See Methods - *Genetic differentiation using Fst*) obtaining 1126 NS variants. The identified non-synonymous mutations clustered into 956 genes (Supplementary Table S2), specifically, 336 were detected only in bald uakaris, 406 only in black and 98 had different mutations in both groups.

Focusing on genes with NS variation in bald uakaris, the dopachrome tautomerase gene (DCT) involved in melanin biosynthesis downstream to *MC1R*, showed a mutation in the position tarseq_64:7868470 (T>C) that causes a NS change (Q>E). This gene has been linked to hair and skin color variation in animals ^36–38^

Further, through a functional overrepresentation analysis, categories of OMIM, KEGG and PANTHER pathway databases were found to be significantly enriched in the above described sets of genes, as well as GO-Biological process categories for bald and black uakaris and HPO in the case of genes affected in both groups (Table 1). The strongest enrichment in the common list of genes (different mutation in each group) was of the *Lipoprotein localization* category from KEGG pathway, nonetheless, there was an overall enrichment for immune-related categories in the three databases (Table 1, Figure S14). On the other hand, *Malaria* - related categories were detected both from OMIM and KEGG pathway databases for bald uakaris (Table 1, Figure S15), although in the latter database the most enriched category was *Myocardial infarction*. Lastly, for genes with fixed NS variation in black uakaris, *Integrin* - related categories were the most significantly enriched in GO-Biological process and KEGG pathway databases, further, *Hypercholesterolemia familial* was the strongest enriched category from the OMIM database (Table 1, Figure S16).

**Table 1.** Significant functional differences between bald and black uakaris. Significant Over-Representation Analysis (ORA) results (FDR ≤ 0.05) on genes with non-synonymous mutations in conserved sites across the primate lineage which were differentially fixed between groups.

|  |  |
| --- | --- |
| **336 genes show fixed mutations only in bald uakaris** |  |
| Malaria<br>ECM-receptor interaction<br>Hematopoietic cell lineage<br>Cytokine-cytokine receptor interaction<br>*KEGG pathway* | Myocardial infarction<br>Malaria, susceptibility to **Malaria**, **resistance to**<br>Lung cancer<br>alveolar cell carcinoma<br>*OMIM* |
| **406 genes show fixed mutations only in black uakaris** |  |
| Integrin activation<br>Regulation of cell cycle phase transition<br>*GO – Biological process* | Hypercholesterolemia familial<br>Schizophrenia<br>Tracheoesophageal fistula<br>Colorectal cancer<br>*OMIM* |
| Integrin signalling pathway<br>*PANTHER pathway* |  |
| **98 genes show different fixed mutations for the two groups** |  |
| Lipoprotein localization<br>Regulation of innate immune response<br>Positive regulation of defense response<br>Immune response-regulating signaling pathway<br>*GO - Biological Process* | Posterior pharyngeal cleft<br>Recurrent bronchitis<br>IgG deficiency<br>Emphysema<br>Leukopenia<br>Lymphadenopathy<br>Abnormality of the lymph nodes<br>Hematological neoplasm<br>Elevated hepatic transaminase<br>Abnormal leukocyte count<br>*Human Phenotype Ontology* |
| Tracheoesophageal fistula<br>Breast cancer<br>Prostate cancer<br>*OMIM* |  |
| ORA significant results ($FDR < 0.05$) | |

## Discussion

We have generated the first population-level study based on whole genome data of a platyrrhine genus: the *Cacajao*, endemic to the Amazon and divided into bald and black uakaris ^16^. By the analysis of 48 geolocalized whole genomes of wild uakari populations, we provide a detailed description of population structure and dynamics as well as to refine the pre-existing phylogenetic and demographic estimates ^12,19,20,27^. Using population-level data, we built a phylogeny based on 1Mb-windows across the whole genome of uakaris. Moreover, narrower confidence intervals have been obtained for divergence times and ancestral or current Ne estimates while migration rates were estimated for the first time. Through this, we have begun to detangle the complex relationship between a highly dynamic environment such as the Amazon rainforest and primate whole genomes. We believe our dataset provides an exceptional representation of all wild uakari species considering the difficulty in sampling wild mammalian populations. Nonetheless, we are aware our study might be constrained by the lack of evenness in population sampling and sometimes limited sample sizes.

### Bald and black uakaris

Bald and black uakaris present strong differences both phenotypically and ecologically ^16,18–20^ which hint at differences at the genomic level that have been described in this study. Since their split, the two groups have taken divergent phylogenetic trajectories and come to represent i) two distinguished genetic clusters of populations (Figure 1B-1D, 2) with ii) different genetic diversity ranges (Figure 3B) and effective population size (*Ne*) estimates (Figure 3A) that display iv) independent population dynamics (Figure 4).

#### Population structure and phylogenetics

By considering the whole autosomal genome, the samples in the dataset clustered into north and south bank of the Negro River for black uakaris and into red and white for bald, overall showing strong population structure sometimes even past the species level. In the phylogenetic inference, we were able to go beyond previous ddRAD-based analysis ^20^ by classifying all the samples into their designated populations (Figure 2) in accordance with population structure analyses (Figure 1B-1D, S4-S6). The only exception to this fine clustering was a sample of HS_LowV, which appeared closer to AY with very low probability (0.08). The overall high level of resolution was reached with high support homogeneously distributed along the genome, which means the signal was not driven by specific regions in the genome when 1Mb windows were employed (Figure 2, S9B, S9C). Nonetheless, slight discrepancies between the latter and the results obtained from employing 250kb windows were observed specifically regarding the red bald uakaris clade (Figure S8). This could be reflecting incomplete lineage sorting due to shallow divergence times in this clade: such signal would be better captured when smaller portions of the genome are employed to build independent window-trees, leading to a more topologically divergent set of trees to build the consensus from.

On the other hand, the phylogeny built from the whole mitochondrial genomes (Figure S10) only recovered the main splits ((white bald, red bald), (ML, (HS, AY))), being unable to get to the species level as reflected in all preceding publications ^12,19,20,27^. The signal retrieved from mitochondrial genomes is limiting since it reflects a partial evolutionary history, that of the female lineage ^39^. It has been proposed that given the high levels of male-male affiliation observed, dispersal in *Cacajao* could be female-biased ^40^. If this was the case, we could hypothesize that prevalent female dispersal inside the resolved clades could be the cause of the observed topology: the phylogenetic trajectory described by a limited number of female lineages does not necessarily have to coincide with the trajectory of all inspected lineages in the genus. Besides, in comparison to the autosomal range, analyses based on mitochondrial genomes rely on a much lower number of markers proportionally, which can hinder the resolution of the most terminal nodes of a tree because of decreased statistical power ^15,28^.

#### Demographic history and current genetic diversity

The first split identified by all phylogenies, which depicts the divergence of black and bald uakaris, was estimated to be 0.91 Mya [0.89-0.98], falling in the Pleistocene as previously suggested ^19,20^ (Figure 3A, S13). This estimate falls inside the confidence interval of ddRAD estimates (1.13 Mya [0.67-1.72] ^20^), but far outside that of estimates based on cytochrome B data (5 Mya ^19^ and 2.38 Mya [1.3-3.58] ^20^). Further, the first intragroup divergences were estimated to happen simultaneously in both groups at around 0.66 Mya (0.66 Mya [0.64-0.68] in bald, 0.66 [0.27-0.82] in black) (Figure 3A, S13). While we remain confident 0.66 Mya is an accurate estimate for the first divergence in bald uakaris, given the wide confidence interval in the estimate for black uakaris we do not rule out the exact match might be artefactual. Nonetheless, in accordance with previous studies based on ddRAD sequences (0.46 Mya [0.29-0.7] in bald, 0.48 Mya [0.27-0.78] in black) ^20^, we believe the divergence time between ML and (HS, AY) does fall inside the estimated confidence interval, still being a close time frame to the first split in bald uakaris and thus a relevant point worth exploring further. In this case, estimates based on mitochondrial sequences are again far from those obtained with whole genomes ^20^. Overall, these comparisons indicate that despite being a partial representation of the whole genome, the level of resolution obtained through the employment of ddRAD sequences is much closer to that obtained via whole genomes than through mitochondrial gene sequences, yet not complete.

On the other hand, we tested three demographic models considering constant *Ne* and estimated values for ancestral (*Cacajao*, bald uakaris, white bald uakaris, black uakaris, ML) and extant populations of AM, CV-Jutaí, NV, HS, and ML (Figure 3A). Given that our sample size was not even among populations and this was limited in some cases, we estimated constant *Ne*s to reduce the complexity of the models, hence acknowledging the constraints of our estimates. Although confidence intervals in the black model were overall wider than those in bald, current estimates reflected that bald *Ne*’s were in general bigger than those of black in those populations included (Figure 3A, S13). In accordance, we saw that genetic diversity in bald populations was higher than black’s, being highest in UC and lowest in AY (Figure 3B), likely reflecting the known largest distribution range in bald uakaris in the case of UC ^23^ and the smallest in black AY ^22^. In this context, both RB populations were the only exception to bald’s higher genome-wide heterozygosity when compared to black’s: RB populations show a very disjunct and small distribution range ^23^, which together with their diversity falling below bald’s range, could point towards a past bottleneck ^6^, although this would need to be explicitly tested. Regarding the estimates of *Ne*, again bald’s higher sizes were with the exception of ML-NegroR (Ne=73499 (43309-80189)), which showed a slightly higher point estimate than CV-Jutaí (Ne=60047 (57298-62436)) – yet displaying a much wider confidence interval. In context with genome-wide heterozygosity, we find that their closeness in *Ne* is not unreasonable given that ML-NegroR was the black population with the highest mean genetic diversity in the group and CV-Jutaí that with the lowest in bald (Figure 3B), with overlapping ranges. We are aware these are constant estimates of effective population size and hence may fail to reflect the true demographic history of the inspected lineages throughout time; nonetheless, the presented estimates are overall coherent with found heterozygosity ranges and thus useful in their interpretation.

#### Independent population dynamics

Despite the disparities in habitat preference and mostly disjunct distribution of bald and black uakaris, their range limits become very close in the area between the right bank of the Negro River and the Solimões River at opposite banks of the Japurá River (Figure 1A). In this region, a new and isolated population of the white bald-headed uakari *C. calvus* was found on an oxbow island of flooded forest that moved from the right bank of the Japurá River to the left; therefore in the same bank of the black uakari *C. melanocephalus* ^23^. Yet, no significant gene flow signal was detected between groups (Figure 4B). We are unable to describe the origin of the observed reproductive isolation, but given the proximity of these two distribution ranges at this point and the fact that other populations in this study showed connectivity despite the presence of the Japurá River (Figure 4A), we hypothesize that the isolation between bald and black uakaris is currently maintained by overall independent population dynamics driven by differential habitat preference, dissimilar phenotypes and relevantly by reproductive incompatibilities. Chromosomal rearrangements have been reported to alter recombination rates in the best cases leading to infertile descendants between differing karyotype-species ^41^. Two distinguishing chromosomal rearrangements have been identified in the karyotypes of *C. rubicundus* and *C. melanocephalus* ^42^, which are probably related to the observed reproductive isolation of bald and black uakaris. On the other hand, divergent habitat specializations have been proposed to affect patterns of gene flow ^34^ - despite populations of bald and black uakaris can be found both in flooded and upland forests, these do not share overall habitat preferences: flooded areas around white or black water rivers and general migratory patterns between habitat types ^19,43^. Furthermore, differing phenotypic features - in this case, notably pelage coloration ^16^ and bald’s characteristic red face ^18^ - have been described as drivers to reproductive isolation in primates ^37,44^ and could be playing a major role in mate choice of the same *Cacajao* group. Likewise, the highly vasculated red face of bald uakaris, not present in black, has been suggested to represent a group-specific communication signal of health status to potential sexual mates ^18^.

On the other hand, past admixture is detected in wild populations in this study (Figure 4), although no admixed individuals are found (Figure S4-S6) and relevantly, population structure is maintained.

The genomic structure of black uakari populations is driven by Negro River ^22^ and Serra do Imeri (Figure 1A, 4A). River width has been suggested to be determinant of population connectivity hindering gene flow in the wider sections of its course ^33,45,46^. Along its course, the Negro River poses a barrier of variable intensity to black uakari populations: at its lower course, the river gets wider ^45^ and this could explain the absence of gene flow signals between AY and ML populations (Figure 4B). Nonetheless, connectivity has been enabled between ML and HS-SaImeri populations in the north-west distribution of black uakaris, beyond the highwaters of the Negro River (Figure 4A) ^45^. Moreover, the populations of HS show differential gene flow patterns by interbreeding with different populations (Figure 4B), which could have been influenced by the presence of the mountain range acting as a barrier: HS-LowV would have interbred with ML and on the right side of the mountain range at a higher altitude, and HS-SaImeri with AY (Figure 4B), the latter relevantly being the strongest detected gene flow signal. The two ML populations, found in the upper Japurá River (ML-JapuráR) and in the right bank of Negro River up to the Solimoes-Japurá confluence (ML-NegroR) respectively, displayed disjunct genomic clustering (Figure 1B) despite the fact that these appeared connected and shared gene flow patterns (Figure 4). The time split between these two populations was estimated at 0.03 Mya [0.02-0.15] (Figure 3A). Field observations have suggested these populations interbreed irrespective of Japurá River, moreover, the existence of a paleochannel connecting Negro and Japurá River has been proposed based on digital elevation model analyses ^35^. However, considering all our results we are not able to discern between high connectivity between these populations or strong ancestry sharing caused by their very recent split time.

In contrast to black populations, whose structure is mainly driven by two prominent barriers, bald uakaris are mostly found in flooded forests of western Amazonia ^23^, a seasonally ^47^ and historically dynamic environment ^35,20,34^. This dynamism could hypothetically explain why *a priori*, current disjunct geographic distribution alone does not hint at the genomic pattern drawn by white and red bald uakari populations. These groups were estimated to diverge 0.66 million years ago [0.64-0.68] (Figure 3A). This is a more recent estimate than those previously introduced based on cytochrome B (0.91 Mya [0.5-1.42] ^20^), but falling in the estimated interval of those based on ddRAD sequences (0.46 Mya [0.29-0.7] ^20^), now with higher confidence. In the first place, seasonal variations in hydrology, which are particularly common in the floodplains of the Amazon basin linked to rain patterns ^48^, have been proposed to affect gene flow by intensifying and diminishing the effect of rivers as effective barriers ^21,31–34^. River dynamics have been proposed to particularly affect population connectivity in systems found in the western part of the Amazon ^12,21,34^, where lowlands are of a much younger age than those in the east, thus shaping a less stable geological topography ^31,33^. In this framework, intermittency of rivers as effective barriers could hypothetically contribute to explain the observed interspecies connectivity in bald uakaris in spite of apparent geographic barriers – in the case of RB populations and CV-Jutaí separated by the Jutaí River, and AM and NV far in the south delimited by the Tarauacá River (Figure 1A, 4) as previously suggested by Silva *et al.* (2023) ^21^. Secondly, it is widely understood that current geographic arrangement represents a very precise moment in time, hence evolutionarily relevant geographic barriers for a given lineage might not be present anymore ^22,32^. In this sense, geography does not need to agree with the observed genomic landscape, which on the contrary summarizes the evolutionary history of a given lineage. Related to this, major geographic rearrangements such as the formation of riverine channels ^48^, specifically between Juruá and Jutaí Rivers ^21,23,35^, have been reported in the Amazon basin throughout the recent geological past (100kya). This river rearrangement could have led to the isolation of white bald uakari populations from red bald populations in opposite banks (Figure 1A) by altering their patterns of geographic distribution ^21^.

#### Genetic differentiation

In order to explore what was the functional impact of the observed independent genomic clustering of bald and black populations, we further inspected which were consistently, the most divergent coding genomic regions between all intergroup pairs of populations (Table 2). Stringent filtering was applied to these to account for the limitations of the annotation being used - which despite being crucial to our study as others alike are to explore non-model organism genomics, was known to be constrained. By focusing specifically on non-synonymous (NS) variants found in codons that showed diversity along the primate phylogeny, we first detected 98 genes with differentially fixed SNPs in both groups (Supplementary Table S2). These were mostly enriched for functions involved in the regulation and performance of the immune system (Table 1, Figure S14), which was not striking in accordance with the extensive number of studies done on genes under selection in primates ^49^. Additionally, 336 (Supplementary Table S2) and 406 genes (Supplementary Table S2) were detected to harbor differentially fixed NS SNPs only in bald or black uakaris respectively. In the set of genes of bald uakaris, we show a NS mutation in the Dopachrome tautomerase gene (*DCT*). Involved in melanin biosynthesis, this gene has been found to be under positive selection in East Asian human populations related to light skin pigmentation ^37^, and to dilute coat color phenotypes when knockout in mice due to reduced melanin content in hair ^36^. Here, from a maximum parsimony perspective, one could assume the ancestral phenotype for *Cacajao* is dark for coat coloration, given that its closest phylogenetic genus, *Chiropotes* ^8^, shares this with black uakaris ^24^. Considering this, our results would match the expectation that bald uakaris should be the group to have diverged from black uakaris, which on the contrary share the ancestral state with the *Pithecia pithecia* reference genome. We are aware that our results do not imply causality for the phenotype of bald uakaris and that further targeted functional analyses should be addressed in order to confirm the link between this mutation and the observed variation in coat coloration. Nevertheless, the genetic landscape of non-human primates has been inspected before with the aim of identifying precise genetic patterns to explain light-dark phenotypic transitions in the phylogeny without concluding results ^37,38,50^, hence we consider this to be a relevant advance and starting point for potential further research. On the other hand, regarding the functional categories enriched in these sets of genes, we found these were less specific than in the common set. Nonetheless, Malaria-related categories for bald (Table 1, Figure S15) and integrin pathway-related categories for black uakaris (Table 1, Figure S16) arised more than once in independent databases. Although the underlying mechanisms leading to the enrichment of integrin pathway-related categories only in black uakaris remain unresolved in this study, differential Platyrrhini-specific variation inside the primate lineage has been identified in the integrin transmembrane receptor family of proteins, which beyond being linked to immune function, have been recognized specifically as receptors to viruses in primates ^51^. On the other hand, a specific species of *Plasmodium* that is regarded as zoonotic Malaria in South American primates, *Plasmodium brasilianum*, is currently known and was first ever reported on the blood of a bald uakari in 1908 ^52^. Regarding this, black uakari species have been hypothesized to be at lower risk of infection ^53^ given that in their common habitat close to blackwater rivers, the density of Malaria-vector mosquitoes is low ^54^. On the contrary, bald uakaris mostly inhabit areas nearby whitewater rivers - with the only exception of the *Jutaí River* ^33^ -, specifically flooded areas which are considered Malaria hotspots ^55^. We hypothesize higher exposure to the parasite throughout time in bald uakaris would explain the observed enrichment for Malaria-related categories in this group in contrast to black. Overall, we consider these are preliminary results on the functional differentiation between bald and black uakaris and encourage future studies to further explore this topic.

#### Conservation perspective

Connectivity is broadly widespread between populations of the same *Cacajao* group (Figure 4), which is particularly relevant for bald uakaris which had been regarded as mostly isolated populations ^23^. This indicates connectivity is a relevant factor in these genus’ population dynamics and to maintaining the health of *Cacajao* populations in the wild ^6^. Climate change effects combined with anthropogenic threats have led to population decline in the past in lemur ^56^, langur ^57^ and tamarin ^58^ lineages via habitat fragmentation and changes in connectivity ^7^. In the case of *Cacajao* populations, black uakaris have been categorized by the IUCN red list of endangered species as low concern (ML ^59^, AY ^60^) and vulnerable (HS ^61^), and display a wider and rather continuous distribution range in contrast to bald uakaris ^22^, being reported to migrate to different habitat types ^19^. Despite these populations show higher habitat versatility and are found in the northern distribution in the Amazon Rainforest - further from the deforestation focus ^62^ - their conservation status is still of concern, in particular, because of the high hunting pressure they are exposed to ^61^.

Bald uakaris are largely specialists of flooded forests, which makes them highly dependent on the wellbeing of this specific habitat ^19,20^. The distribution of bald uakaris is disjunct (Figure 1A), with the most extreme example being the sampled population of CV-MSDR, which is known to be found in a flooded forest area limited by the Solimões and Japurá Rivers isolated from the remaining bald populations ^20^, and effectively isolated from a genomics perspective as observed in this study too. The *Cacajao* genus has been under thorough taxonomic review in the recent past ^16,20^, probably the reason why all bald uakaris are still assessed as subspecies of *C. calvus*, following the taxonomic classification of Hershkovitz (1987). Accordingly, two taxa were categorized as Vulnerable – *ucayalii* ^63^ and *novaesi* ^64^ and two more – *calvus* ^65^ and *rubicundus* ^66^ – as Least Concern, likely in relation to their northern distributions ^62^. Also, *Cacajao amuna* is the only taxon not assessed yet as it was recently described ^20^.

Although habitat loss is currently not an immediate threat for all these populations, particularly for those of northern distributions (e.g. *C. calvus* and *C. rubicundus)* ^62^, it is relevant to acknowledge their vulnerability linked to habitat specificity ^20^. There is yet a relevant exception in populations of *C. ucayalii* (UC was sampled in this study), which are the only bald uakaris to be reported at high altitude non-flooded forests and show the largest distribution range ^23,67^. In accordance with field observations ^23^, the UC population sampled in this study showed no significant signals of interbreeding with any other bald population, which significantly may suggest a crucial interplay between their habitat versatility and such high genetic diversity levels, highest in the *Cacajao* genus, despite being reproductively isolated from the rest of the populations in this study. Contrary to this, CV and RB populations are the most flooded-forest specialists in the genus and present the most restricted distribution ^23^, accordingly, in this study the sampled populations from these species show the lowest genetic diversity and estimated *Ne* (CV) in bald uakaris.

*Cacajao* is however one more case in the global panorama, where we find more than half of the extant primate lineages are under extinction threat ^7,8^. With the aim of starting to fill the notable gap of knowledge in non-Catarrhini (mostly non-great ape) wild populations’ genomics 8,10–12, we hope the understanding of uakari wild population dynamics and their respective degree of genetic diversity may contribute to the implementation of effective conservation actions to face increasing rates of wild biodiversity loss in the near future 2,6,7,68.

## Methods

### Sample processing & Data generation

We used blood and tissue samples from 48 *Cacajao* individuals (N_bald=30, N_black=18) and 1 *Pithecia* from wild populations on the Brazilian Amazon with known geographical coordinates. The latter was added as an outgroup in certain analyses. All samples in this study were muscle tissue samples from wild-caught uakaris preserved in 70%-90% ethanol obtained from Brazilian zoological collections in Instituto Nacional de Pesquisas da Amazônia (INPA), Universidade Federal do Amazonas (UFAM) and Instituto de Desenvolvimento Sustentável Mamirauá (IDSM). The individuals analyzed were either sampled during multiple commissioned large field surveys of the biodiversity of the Amazon rainforest by the Brazilian government (2000-2017) or as part of monitoring programmes, from hunted individuals by local communities. Collection permits were obtained from the Biodiversity Authorization and Information System (SISBIO; permit nos. 55777, 42111, 32095–1, 7795–1) and exported under CITES permits (19BR033597/DF and 15BR019039/DF). We extracted the DNA with the MagAttract HMW DNA extraction kit and prepared short-inserted paired-end libraries for the whole genome with the KAPA HyperPrep kit (Roche) PCR-free protocol. From these, we sequenced paired-end reads with a 2×151+18+8bp length on NovaSeq6000 (Illumina) to reach at ~30X coverage per sample. For more details on the data generation steps see Supplementary Methods in Kuderna *et al.* (2023) ^8^.

### Data processing

We interleaved the reads with seqtk mergepe (v1.3) ^69^ and subsequently trimmed the adapters with cutadapt (v3.4, –interleaved) ^70^. We then mapped the trimmed reads using bwa-mem (v0.7.17, default settings) ^71^ to the reference assembly of the *Pithecia pithecia* species at the scaffold level presented in Shao *et al* 2023 ^13^. We used a reference genome from a genus outside *Cacajao* for the following reasons, i) given that *Cacajao* is a non-model organism *Pithecia pithecia* was the closest species with an available reference genome and ii) we considered using a sister but external taxon as reference to our analyses was a necessary direction in order to minimize biases in our work related to reference bias and ancestry sharing. The reference genome had a length of 2.72 Gbp and a 10,87 Mbp scaffold N50. We proceeded to mark duplicates and add read groups employing bbmarkduplicates (default settings) from biobambam (v2.0.182) ^72^ and the AddOrReplaceReadgroups (default settings) from PicardTools (v2.8.2) ^73^ respectively. The mode of the depth was calculated for all samples independently using MOSDEPTH (v0.3.3) ^74^.

We employed the resulting cram files for the calling of variants in a three-step manner using the GATK (v4.1.7.0) ^75^ toolkit. We generated eighty-nine equal-size windows from the reference genome to parallelize the process for the sake of computation time. At first, we generated a GVCF file using the HaplotypeCaller algorithm (-ERC BP_RESOLUTION) per each of the samples and chunks respectively. Secondly, we combined all variants called in all samples for a given chunk in the reference genome using CombineGVCFs (default settings) to be finally jointly genotyped employing GenotypeGVCFs (default settings).

Hard filtering was applied to the resulting SNPs in each chunk by removing those variants that were not biallelic (vcflib vcfbilallelic (v10.5) ^76^), those with depth values outside the range between 70 (double the median maximum coverage across samples) and 5 (one third of the median minimum coverage across samples), and missingness higher than 60% using vcftools (v0.1.12) ^76,77^. Furthermore, following GATK Best Practices Protocol, we excluded those SNPs not complying with the following expression: “QD < 2 | FS > 60 | MQ < 40 | SOR > 3 | ReadPosRankSum < −8.0 | MQRankSum < −12.5” employing GATK VariantFiltration. We merged the remaining SNPs in each chunk-VCF into a single dataset which we then filtered for allele imbalance by keeping variants with frequencies within the range of 0.25-0.75 (Supplementary Methods (Section 1.2), Figure S1). Lastly, we removed SNPs in scaffolds belonging to sex chromosomes (identified in Kuderna *et al.* (2023) ^8^) and those in scaffolds shorter than 0,5Mb from the dataset, thus keeping 367 scaffolds.

### Data analysis

#### Genome-wide heterozygosity

We calculated genome-wide heterozygosity in windows of 100kbp. For each sample independently, we extracted the number of callable base pairs in the given window region together with the number of heterozygous positions. These values were used (i) to calculate the mean heterozygosity per individual in the dataset as well as (ii) to plot the heterozygosity distribution across the genome. Plots were generated using the ggplot2 ^78^ package in R (v4.2.2) ^79^.

#### Relatedness and Inbreeding

We used NGSRelate2 (v2.0) ^80^ to calculate relatedness and inbreeding in the dataset, which employs the Jacquard coefficients to estimate the kinship and inbreeding coefficients respectively. We assessed relatedness based on the *theta* parameter (Figure S2). We then split the full dataset into sub-datasets based on the taxonomy to calculate relatedness and inbreeding so these were not overestimated due to population structure. Plots were generated using the ggplot2 package in RStudio.

At this point, we independently filtered the full dataset and the subset of bald individuals using PLINK (v2.0, --maf 0.019 --geno 0.4) ^81^. Furthermore, we only filtered the dataset including the black species by --maf 0.05 --geno 0.4, employing a different minimum allele frequency filter due to the difference in number of individuals. These datasets were used to run EEMS and SMC++ (See below). Then, in all three datasets, we obtained independent sites by pruning the datasets of linkage disequilibrium using the default settings (--indep-pairwise 50 5 0.5). These datasets were employed to run PCA and ADMIXTURE (See below).

#### Population structure

##### PCA

We generated three PCA plots using smartPCA from EIGENSOFT (v7.2.1) ^82^. First, we calculated the eigenvalues and eigenvectors on the full dataset (bald and black samples), on the bald species subset, and on the black species subset. Plots were generated using the ggplot2 package in R (v4.2.2). When compared, Figure 1C and Supplementary Figure 7A depict differences before and after applying the above mentioned LD pruning filter. Supplementary figures were generated with PLINK (v2.0) *--pca* and plotted in the same way as the main figures.

##### ADMIXTURE

ADMIXTURE ^83^ software was run on the full dataset (bald and black), and on both partial datasets independently. We generated fifty different random seeds for the runs on the different datasets so as to compare the results of independent runs and assess the model convergence in each of the three cases. The tested number of ancestral populations (K) ranged from 2-13 in the full dataset and from 2-8 in the partial datasets. Plots were generated with the ggplot2 package in R (v4.2.2) using the data in those runs with the lowest cross-validation error.

#### Connectivity and gene flow

##### Testing deviations from *Isolation by Distance*

Using the Effective Migration Surfaces (EEMS) ^84^ software, we estimated migration and diversity rates in our dataset by looking for deviations from an isolation by distance model. We run this three times, one on the full dataset (bald and black), and the other two on the partial datasets. We generated dissimilarity matrices with EEMS’ bed2diffs_v1 program. We defined a geographic range in each of the cases based on geographic distribution knowledge obtained from the IUCN website. The software was run for 9M iterations in each of the cases with a number of thinning iterations of 9999. We assessed the convergence of the MCMC and completeness in each of the times by placing parallel and independent runs. Plots were generated in R (v4.2.2) using the reemsplots2 package ^85^.

##### Gene flow quantification using D & fstatistics

To calculate D and *f* statistics we ran the DSUITE software ^86^ on 191,4M bi- and multi-allelic SNPs. This was the only analysis where both types of variants were used - bi- and multi-allelic SNPs were employed here given the software allowed it with the aim of increasing statistical power. We also calculated the *f*-branch metric ^82^ as a simplification of *f* statistics in large phylogenies with a high proportion of correlated values. We used the sequenced *Pithecia pithecia* sample (different from the reference) as an outgroup. To obtain *f*-branch values depicting gene flow evidence in the dataset ^82^, the *Dtrios, Fbranch* and *dtools.py* tools from DSUITE were employed.

#### Phylogenetics

##### Whole genome phylogeny

From each of the samples and the one *Pithecia pithecia* outgroup sample above mentioned, we generated 2144 1Mb-long and 9313 250kb-long windows based on the autosomal regions in the *Pithecia pithecia* reference genome. These covered 78.78% and 85.55% of the reference genome respectively. We obtained consensus sequences using ANGSD (v0.931, -doFasta 1) ^87^. The sequences corresponding to each window and respective length were then piled up thus forming a multiple sequence alignment (MSA) which was trimmed with trimAL (v1.4.1) ^88^. We then independently inputted these to iqTree2 (v2.1.2, -B 1000) ^89^ to generate a maximum likelihood (ML) tree per window after 1000 bootstrap replicates and automatic model selection. Two consensus trees were built by merging trees obtained from windows of a given length using BEAST’s TreeAnnotator ^90^.

##### Topological distance between 1Mb Window trees vs. Consensus tree

To assess the distribution of branch support across the genome, we calculated the topological distances from each 1Mb window tree to the respective consensus tree. Two metrics were applied using ape.dist.topo function from ape R package (v5.6-2) ^91^: PH85 and score. The latter metric takes into account branch lengths.

##### Mitochondrial phylogeny

To run MitoFinder ^92^, we used a subset of the trimmed raw reads representing 4% of the total set per sample. Inside MitoFinder, we employed MetaSPADES to assemble into and Arwen ^93^ to annotate the mitochondrial genomes of each of the samples. We reordered all the contigs with Fasta-tools SHIT ^94^ option so all start at the CYTB gene. MAFFT ^92^ was then used to generate a MSA of the mitochondrial contigs, which was in turn trimmed using trimAL to generate a phylogenetic tree using iqTree2 (v2.1.2, -B 1000).

All figures of phylogenetic trees were built using FigTree (v1.4.4)^95^.

#### Genetic differentiation using Fst

##### Average differentiation genome wide

We generated PLINK (v2.0) binary files with a 0.6 allowed missingness which was fed to the KRIS R package ^96^. The fst.hudson function was used to calculate the average pairwise Hudson *F*_ST_ ^97^ along the genome among all the identified populations in the dataset combining taxonomic data and observed population structure.

##### Differentiation in coding regions, analysis of non-synonymous variants

To calculate Hudson *F_ST_* values per SNP in each pairwise comparison between the studied populations we used PLINK (v2.0). After this, we kept variants in the 99%-upper quantile of the *F*_ST_ distribution detected in all inter-group comparisons. As the employed VCF here was biallelic, these corresponded to differentially fixed variants in each of the two groups. Out of these, we investigated those variants in coding sequence (CDS) regions. These were identified using the annotation of the respective assembly based on human orthologs (Valenzuela-Seba et al. [in preparation] ^98^). We retrieved the sequence of those codons containing the kept variants both (i) in the REF state from the original reference genome employing Samtools faidx ^99^ and (ii) in the ALT state from the filtered VCF employing Bcftools consensus ^99^. We then proceeded to translate these codons and kept only those that contained a variant causing a non-synonymous mutation (6442 SNPs). These were then filtered based on their alignability and on their conservation in the primate order. First, given that the quality of the genomic annotation employed was low, after filtering we only kept codons that i) were confidentially aligned throughout the primate lineage and ii) those falling in genes without frameshifts (99.99% of the total) (3992 SNPs). The MSA including most primate lineages was obtained from Valenzuela-Seba et al. [in preparation] ^98^. We then grouped the filtered codons by gene, getting a total of 3143. Second, by assessing the amino acidic conservation across the primate lineage based on the Shannon entropy coefficient calculated in (Valenzuela-Seba et al. [in preparation] ^98^), we kept only those positions in our dataset with lower levels of conservation across the whole primate phylogeny (positions in the top 75% quantile of the Shannon entropy distribution) in order to filter out positions so highly conserved at the order level that would show no signal at the genus level. By doing this, the list was reduced to 1126 positions, which fell into 956 genes. We divided this list of genes into three groups: genes which showed non-synonymous mutations only in bald uakaris (336) only in black (406) and finally those affected in both groups - although by different variants (98) (Supplementary Table S2). We then employed the Over-Representation Analysis (ORA) option from the WebGestalt tool ^100^ with default parameters to inspect functional enrichment in each of these gene lists independently. Here, we used the initial set of ortholog genes as background - excluding the genes with frameshifts. The following databases were tested for significant over-representation: GO (Biological Process non-redundant), KEGG, PANTHER pathway, OMIM and HPO. We report significant results (FDR ≤ 0.05).

#### Modeling demographic history

##### Maximum Likelihood modeling with *fastsimcoal2*

We generated a VCF including monomorphic positions by adding --select-variant-type NO_VARIATION with the GATK (v4.1.7.0) VariantFiltration option including those autosomal scaffolds longer than 0,5Mb (367). No missingness was allowed across all samples and only biallelic sites were included. In order to minimize intra-population structure, widespread gene flow and coherent sample sizes between populations, we only included a subset of the samples in the general dataset (Supplementary Table S5). Coding regions predicted on the *Pithecia pithecia* reference genome were removed in order to keep putatively neutral positions based on the annotation from (Valenzuela-Seba *et al.* [in preparation] ^98^). Folded multiSFS to be used as input for fastsimcoal2 ^101^ were generated with the associated vcf2sfs.py and foldSFS.py scripts based on 162,75M SNPs and 1.953M monomorphic positions.

##### Models details

We generated three models, a general one including bald and black samples, and two more for each of the groups alone. We included seven samples from the population of *C. calvus* from *Jutaí River* (CV-Jutaí) and six from *C. melanocephalus* from *Japurá River* (ML-JapuráR) in the bald and black model. Then, we sampled seven individuals from *C. calvus* from *Jutaí River* (CV-Jutaí) and *C. amuna* (AM) representing white bald uakaris and six from *C. novaesi* (NV) for red bald uakaris for the bald model. Finally, six samples from *C. melanocephalus* from *Japurá* (ML-JapuráR), five from *Negro* river’s (ML-NegroR) and four samples from both *C. hosomi* populations - given their very low genetic differentiation - as HS, were included in the black model. For more details about the models see Supplementary Methods (Section 1.3.1).

The mutation rates used in these models were calculated in Kuderna *et al*. (2023) ^8^: the mean between *C. calvus* and *C. melanocephalus’* mutation rates was used for the bald and black model, *C. calvus*’ mutation rate was used in the bald model and the mean between *C. hosomi*’s and *C. melanocephalus*’ in the black one.

##### SMC++ demographic inference

To compensate for the fact that *Ne* is assumed constant in the previous models, we used the SMC++ software to investigate the fluctuations of *Ne* through time. Making use of the four populations defined and respective sample sets used for the group-specific maximum likelihood models (White bald uakaris, red bald uakaris, south bank Negro River black uakaris and north bank Negro River black uakaris), we created a .smc file employing the vcf2smc program from the SMC++ (v. 1.15.2) ^102^ software per group and each of the 367 remaining scaffolds after filtering in previous steps. After that, using the associated estimate program (--spline cubic --timepoints 0 100000), we generated a .model file per group applying an averaged mutation rate per generation across all Cacajao estimates in Kuderna et al. (2023) ^8^. We built one single plot with all group estimates together using the plot script from SMC++ and indicating a generation time of 10 years based on Kuderna et al. (2023) ^8^.

## Supporting information

Supplementary Table S1: Sample metadata

Supplementary Table S2: Sets of genes showing non-synonymous (NS) variation considered for ORA analyses.

## Acknowledgements

T.M.B gratefully acknowledges the financial support from the European Research Council (ERC) under the European Union’s Horizon 2020 research and innovation programme (grant agreement No. 864203), (PID2021-126004NB-100) (MICIIN/FEDER, UE) and from the Secretaria d’Universitats i Recerca and CERCA Programme del Departament d’Economia i Coneixement de la Generalitat de Catalunya (GRC 2021 SGR 00177). J.P.B. gratefully acknowledges the financial support from the Natural Environment Research Council (NERC) (NE/T000341/1). F.E.S. gratefully acknowledges the financial support from the European Union’s Horizon 2020 research and innovation programme under the Marie Skłodowska-Curie grant agreement (801505), the Fonds National de la Recherche Scientifique (F.R.S.-FNRS, Belgium; grant 40017464) Brazilian National Council for Scientific and Technological Development (CNPq) (Processes 303286/2014-8, 303579/2014-5, 200502/2015-8, 302140/2020-4, 300365/2021-7, 301407/2021-5, #301925/2021-6), the International Primatological Society (Conservation grant). The Rufford Foundation (14861-1, 23117-2, 38786-B), the Margot Marsh Biodiversity Foundation (SMA-CCO-G0023, SMA-CCOG0037), the Primate Conservation Inc. (1713 and 1689) and the Gordon and Betty Moore Foundation (Grant 5344) (Mamirauá Institute for Sustainable Development). N.H.-A. gratefully acknowledges the financial support from the Government of Catalonia | Agència de Gestió d’Ajuts Universitaris i de Recerca (Agency for Management of University and Research Grants) (-FI_00040).

## Competing interests

The authors declare that they have no competing interests.

## Data and materials availability

All data needed to evaluate the conclusions in the paper are present in the paper and/or the Supplementary Materials as well as all the necessary parameters to run the used programs. For further doubts or inquiries contact. Reference genome assembly for *Pithecia pithecia* is available in GenBank with ID GCA_023779675.1. Raw sequencing data for the 48 *Cacajao* samples analyzed will be available in a public repository by the time of publishing.

## Notes

### Competing Interest Statement

The authors have declared no competing interest.

### Summary of Updates

The changes suggested by reviewers. These include the addition of a demographic inference analysis, the exploration of the genetics of coat coloration differences between uakari groups, the correction of format errors and addition of clarifications to certain statements.

