## Supplementary Material for "Whole genomes of the amazonian *Cacajao* reveal complex connectivity and fast differentiation driven by high environmental dynamism"


Supplementary Information 1
1. Methods 3
1.1. Labeling system 3
1.2. Data processing 4
1.3. Data analysis 5
1.3.1. Modeling demographic history 5
Models details 5
Bald and black 5
Bald 5
Black 5
Fastsimcoal2 run details 7
2. Results 8
2.1. Population structure and genus description 9
2.1.1. Relatedness 9
2.1.2. Heterozygosity 10
2.1.3. Structure analysis 11
2.1.4. Principal Component Analysis 15
2.2. Phylogenetics 18
2.2.1. Whole genome - 250kb windows 18
2.2.2. Whole genome - 1Mb windows 20
2.2.3. Whole mitochondrial genome 26
2.3. Gene flow 28
2.3.1. Fst 28
2.3.2. EEMS 29
2.4. Demographic inference 30
2.4.1. fastsimcoal2 30
2.4.2. SMC++ demographic inference 31
2.5. Genetic differentiation 32

### 1. Methods

#### 1.1. Labeling system

The labeling system includes 12 populations which were named by considering i) the taxonomy, ii) the geographical location of the given population (Figure 1A) and iii) the potential intra-species genomic differentiation. In the text, we employ the labels for the sampled populations unless we refer to populations not sampled in this study.

| **Sampled population label** | **Group** | **Taxonomy** | **Specific sampling area** |
|---|---|---|---|
| AM | Bald | *Cacajao amuna* | Tarauacá River, right bank, Brazil |
| AY | Black | *C. ayresi* | Aracá River, right bank, Brazil |
| CV-MSDR | Bald | *C. calvus* | Mamirauá Sustainable Development Reserve, Brazil |
| CV-Jutaí | Bald | *C. calvus* | Jutaí River, right bank, Brazil |
| HS-SaImeri | Black | *C. hosomi* | Serra do Imeri, Brazil |
| HS-LowV | Black | *C. hosomi* | Lowland forests, south Venezuela |
| ML-NegroR | Black | *C. melanocephalus* | Low Japurá and Negro interfluve, Brazil |
| ML-JapuráR | Black | *C. melanocephalus* | Upper Japurá River, Brazil |
| NV | Bald | *C. novaesi* | Tarauacá River, left bank, Brazil |
| RB-JutaíSolimões | Bald | *C. rubicundus* | Jutaí-Solimões interfluve, Brazil |
| RB-JacurapáR | Bald | *C. rubicundus* | Jacurapá channel, Brazil |
| UC | Bald | *C. ucayalii* | Serra do Divisor National Park, Brazil |

**Table S3.** Description of the labeling system designed for the study.

#### 1.2. Data processing

After mapping the reads to the reference genome and filtering the called variants in our dataset by coverage (min: 5, max: 70) and "QD < 2 | FS > 60 | MQ < 40 | SOR > 3 | ReadPosRankSum < -8.0 | MQRankSum < -12.5", we obtained the following allele balance distributions (Supp Fig*). These were then filtered for allele imbalance by keeping variants with frequencies within the range of 0.25-0.75.

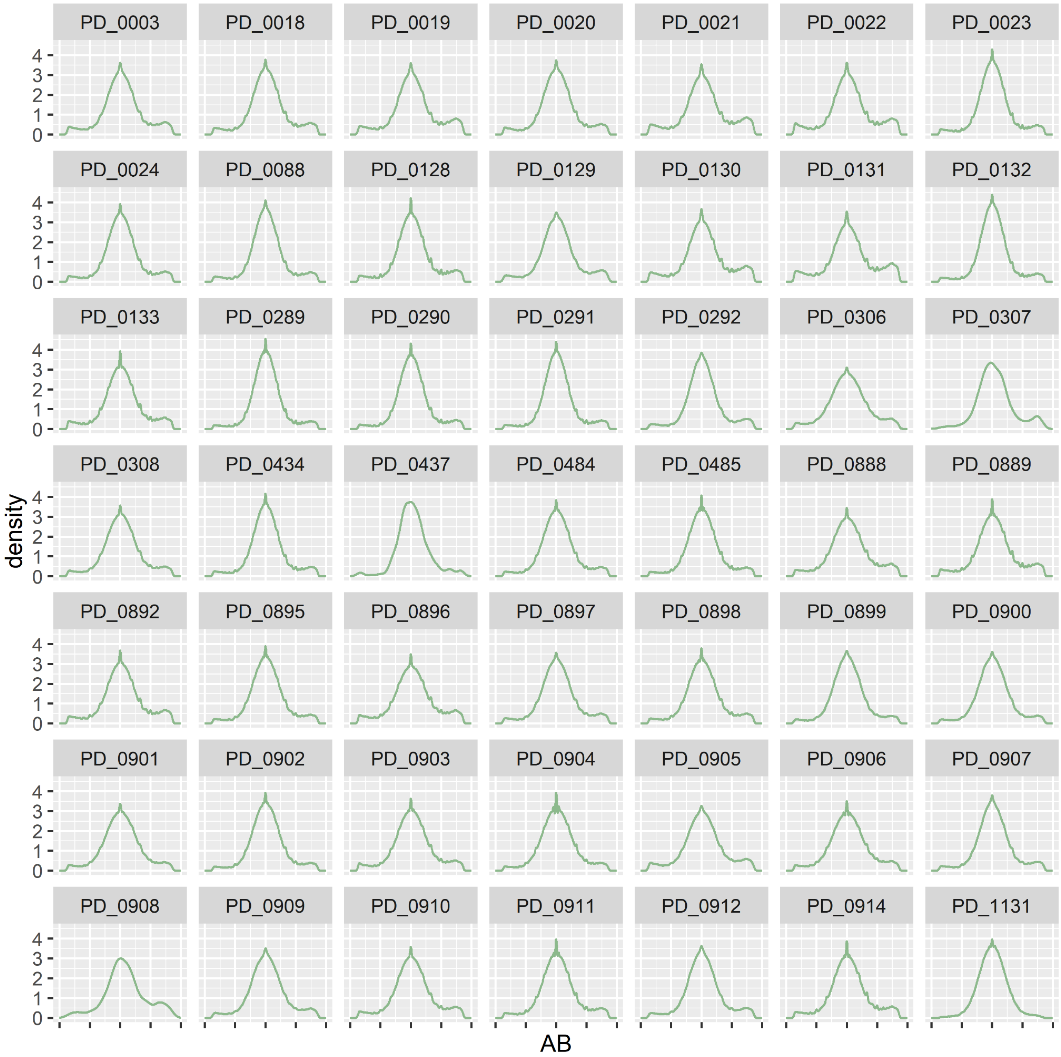


**Figure S1.** Allele balance distribution in dataset and outgroup sample.

### 1.3. Data analysis

#### 1.3.1. Modeling demographic history

##### Models details

We generated three models, a general one including bald and black samples, and two more for each of the groups alone. With these models we estimated divergence times, constant effective population sizes (*Ne*) and in the bald and black models, also effective migration rates per generation were estimated as nuisance parameters. In total, we estimated twenty-one parameters, where *Ne* for the ancestors of bald and black uakaris were estimated twice: The aim of the general bald and black model was to estimate the divergence between these two and the *Ne* of the ancestor of *Cacajao*, for this reason, to inspect further parameters such as the *Ne* of bald and black ancestors, that of extant populations and intra-group divergence times we refer to each of the particular and more complete models. In order to minimize intra-population structure, widespread gene flow and coherent sample sizes between populations, we only included a subset of the samples in the general dataset (Table S5).

###### Bald and black

Samples from the populations of *C. calvus* from *Jutaí* river (CV-Jutaí) and *C. melanocephalus* from *Japurá* river (ML-JapuráR) were considered for this model. Seven and six individuals were sampled from each population respectively. With this data four parameters were estimated. These included the effective population sizes for (i) the ancestor of *bald* and *black* uakaris, (ii) the ancestor of *bald* uakaris and (iii) the ancestor of *black* uakaris and the time split between the ancestors of *bald* and *black* uakaris.

###### Bald

Samples from the populations of *C. calvus* from *Jutaí* river (CV-Jutaí) and *C. amuna* (AM) also representing white bald uakaris and *C. novaesi* (NV) for red bald uakaris. Seven individuals were sampled from the CV-Jutaí and AM populations six from NV. With this data eight parameters were estimated, including:  1) splits between (i) *red* and *white bald* uakaris and (ii) inside *white* uakaris into the respective sampled populations; 2) effective population sizes of (i) the ancestor of *bald* uakaris, (ii) the ancestor of *white bald* uakaris, (iii) AM, (iv) CV-Jutaí, (v) NV; and finally, the 3) effective migration rate per generation between AM_NV which was employed as a nuisance parameter.

###### Black

Four samples from both *C. hosomi* populations were used as HS given their low genetic differentiation. Six samples from *C. melanocephalus* from *Japurá* (ML-JapuráR) and five from *Negro* (ML-NegroR) river's were also used. With these, nine parameters were estimated, including:. These include  1) time splits (i) the ancestor of *black* uakaris into the respective sampled populations and that of (ii) the ancestor of ML into the two sampled populations; effective population sizes of (i) the ancestor of *black* uakaris, (ii) the ancestor of ML populations, (iii) HS, (iv) ML-JapuráR, (v) ML-NegroR; and finally, the effective migration rate per generation between (i) HS and the ancestor of the sampled ML populations, and (ii) between the two ML populations which were employed as nuisance parameters.

| Model | Sampled populations | Samples per population | Parameters | Topology |
|---|---|---|---|---|
| Bald and black | CV-Jutaí<br>ML-JapuráR | 7<br>6 | Tsplit bald/black<br>*Ne* Ancestral *Cacajao*<br>*Ne* bald<br>*Ne* black | 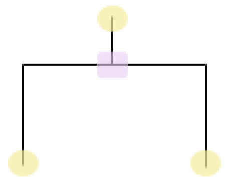 |
| Bald | AM<br>CV-Jutaí<br>NV | 7<br>7<br>4 | Tsplit Ancestral bald<br>Tsplit white bald<br>*Ne* Ancestral bald<br>*Ne* Ancestral white bald<br>*Ne* AM<br>*Ne* CV-Jutaí<br>*Ne* NV<br>Migration rate AM_NV | 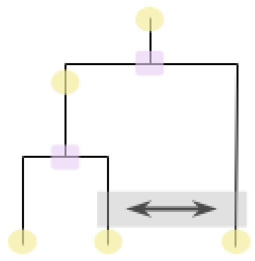 |
| Black | HS (Salmeri + LowV)<br>ML-JapuráR<br>ML-NegroR | 4<br>6<br>5 | Tsplit Ancestral black<br>Tsplit Ancestral ML<br>*Ne* Ancestral black<br>*Ne* Ancestral ML<br>*Ne* HS<br>*Ne* ML-JapuráR<br>*Ne* ML-NegroR<br>Migration rate HS_Ancestral ML<br>Migration rate ML populations | 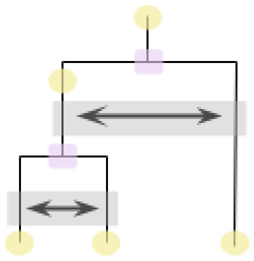 |

**Table S4.** Maximum likelihood models summary: sampled populations, number of samples per population, estimated parameters and model topologies.

| Sample ID | Population | Included in model |
|---|---|---|
| PD_0289 | AM | Bald |
| PD_0291 | AM | Bald |
| PD_0484 | AM | Bald |
| PD_0485 | AM | Bald |
| PD_0899 | AM | Bald |
| PD_0903 | AM | Bald |
| PD_0904 | AM | Bald |
| PD_0290 | CV-Jutaí | Bald, Bald and black |
| PD_0306 | CV-Jutaí | Bald, Bald and black |

| | | |
|---|---|---|
| PD_0307 | CV-Jutaí | Bald, Bald and black |
| PD_0308 | CV-Jutaí | Bald, Bald and black |
| PD_0437 | CV-Jutaí | Bald, Bald and black |
| PD_0888 | CV-Jutaí | Bald and black |
| PD_0895 | CV-Jutaí | Bald, Bald and black |
| PD_0003 | HS-Salmeri | Black |
| PD_0889 | HS-Salmeri | Black |
| PD_0129 | HS-LowV | Black |
| PD_0914 | HS-LowV | Black |
| PD_0128 | ML-JapuráR | Black, Bald and black |
| PD_0898 | ML-JapuráR | Black, Bald and black |
| PD_0905 | ML-JapuráR | Black, Bald and black |
| PD_0906 | ML-JapuráR | Black, Bald and black |
| PD_0911 | ML-JapuráR | Black, Bald and black |
| PD_0912 | ML-JapuráR | Black, Bald and black |
| PD_0018 | ML-NegroR | Black |
| PD_0907 | ML-NegroR | Black |
| PD_0908 | ML-NegroR | Black |
| PD_0909 | ML-NegroR | Black |
| PD_0910 | ML-NegroR | Black |
| PD_0023 | NV | Bald |
| PD_0132 | NV | Bald |
| PD_0900 | NV | Bald |
| PD_0902 | NV | Bald |

**Table S5.** Samples included in particular models.

##### Fastsimcoal2 run details

fastsimcoal2 starts with random initial parameter values sampled from a specified distribution and performs a series of expectation conditional maximization (ECM) optimization cycles. We set uniform priors for effective population sizes and migration rates borned by arbitrary values (100-100,000 and 0.000001-0.0001 respectively, with the exception of the ancestral effective population size for each model, which upper bound was set to 1,000,000), and uniform priors for divergence times borned by the lower and upper bounds of previous ddRAD-seq estimates (*20*). To avoid local maxima, the same demographic scenario was simulated several times with varying seeds. We performed 600,000 simulations, 65 ECM cycles and 100 replicate runs starting from different random initial values. To limit overfitting, only SFS entries with more than 5 counts were considered (*-C 5*).

To maximize the fit between the expected and observed SFS, we used the approach described in Choin, Mendoza-Revilla, Rubio-Arauna et al. 2021 (*99*) . To obtain the maximum-likelihood (ML) estimates of demographic parameters for a given model, we first selected the 10 runs, among the 100 replicate runs, with the highest likelihood. To account for the stochasticity inherent to the approximation of the likelihood using coalescent simulations, we re-estimated the likelihood of each of the 10 best runs, using 100 expected SFS obtained using 600,000 simulations. Finally, we refined the likelihood of the three runs with the highest average, re-estimated log10(likelihood) using 10,000,000 simulations, and considered the run with the highest likelihood as the ML run. To correct for the different number of SNPs in the expected and observed SFS, we rescaled the parameters by a rescaling factor (RF) defined as $S_{obs}/S_{exp}$: effective population sizes and split times were multiplied by RF, while migration rates were divided by RF.

We calculated confidence intervals with a non-parametric block bootstrap approach. We generated 100 bootstrapped folded-SFS with the associated scripts vcf2sfs.py and foldSFS.py, and re-estimated parameters using the same settings as for the original dataset with 20 replicate runs. To obtain the 95% confidence intervals, we calculated the 2.5% and 97.5% percentiles of the estimates' distribution obtained by non-parametric bootstrap. The estimates derived from the 20 runs on the 100 bootstrapped folded-SFS of the *black* model showed a bimodal distribution, probably due to a lack of convergence in some bootstrap datasets. For this reason, additional 80 runs were performed per replicate. Out of these 100 runs, the 20 with the best likelihood were chosen for each bootstrap replicate to obtain the 95% confidence intervals of the model estimates. In this way, bimodality in the parameter estimate distributions was removed.

### 2. Results

Here, we generated 48 whole genomes at a mean coverage of 31x ranging from 14x to 43x (Table S1) in which we discovered a total of 191,4M bi and multi-allelic SNPs.

#### 2.1. Population structure and genus description

##### 2.1.1. Relatedness

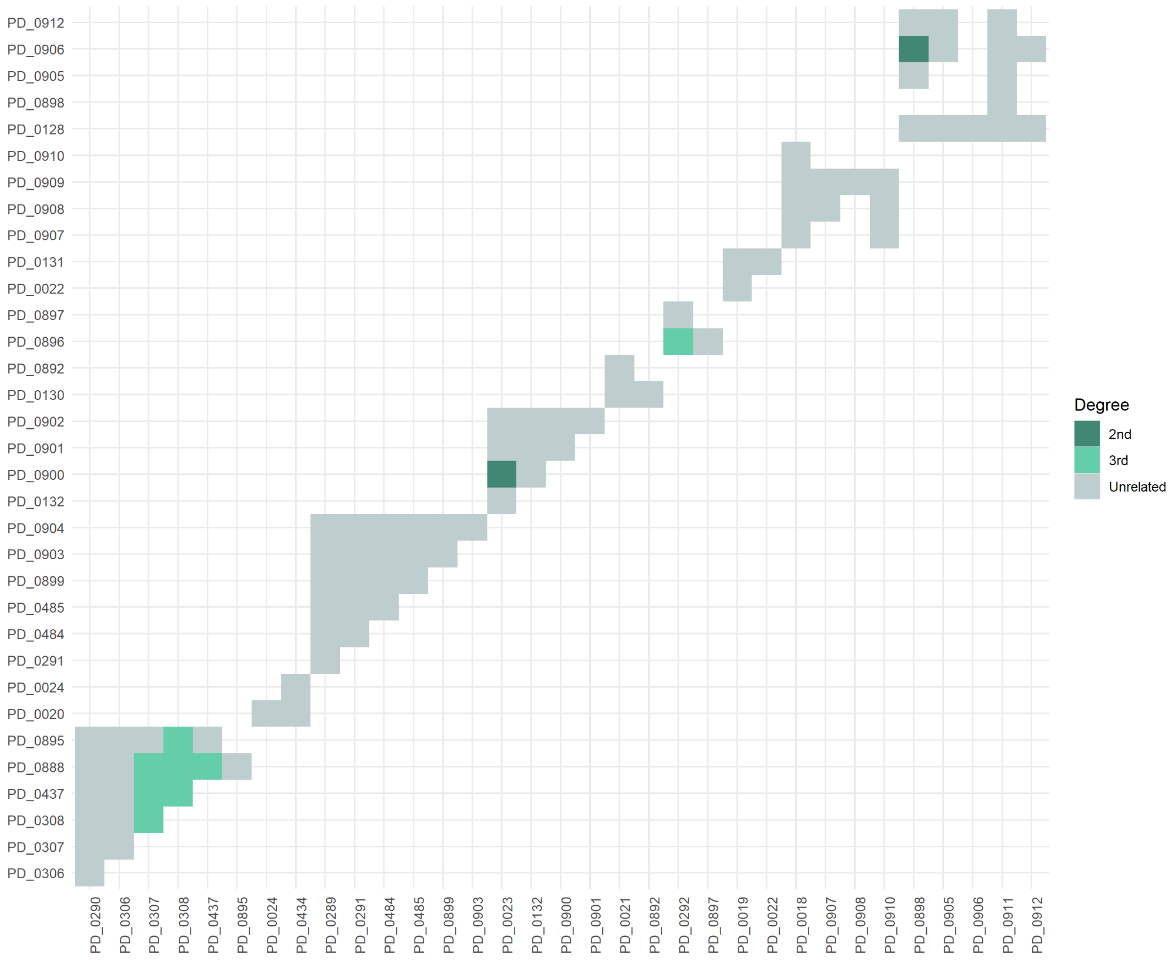


**Figure S2.** Relatedness degree from NGSRelate2 run independently per population and merged to plot, see sample clusters. Colors in the outer x and y axis indicate population. Five categories to depict relatedness: $Self \geq 0.375,\ 1st \geq 0.187,\ 2nd \geq 0.062,\ 3rd \geq 0.001,\ unrelated\ = 0$ based on Wang *et al* (2010, COANCESTRY: a program for simulating, estimating and analyzing relatedness and inbreeding coefficients).

#### 2.1.2. Heterozygosity

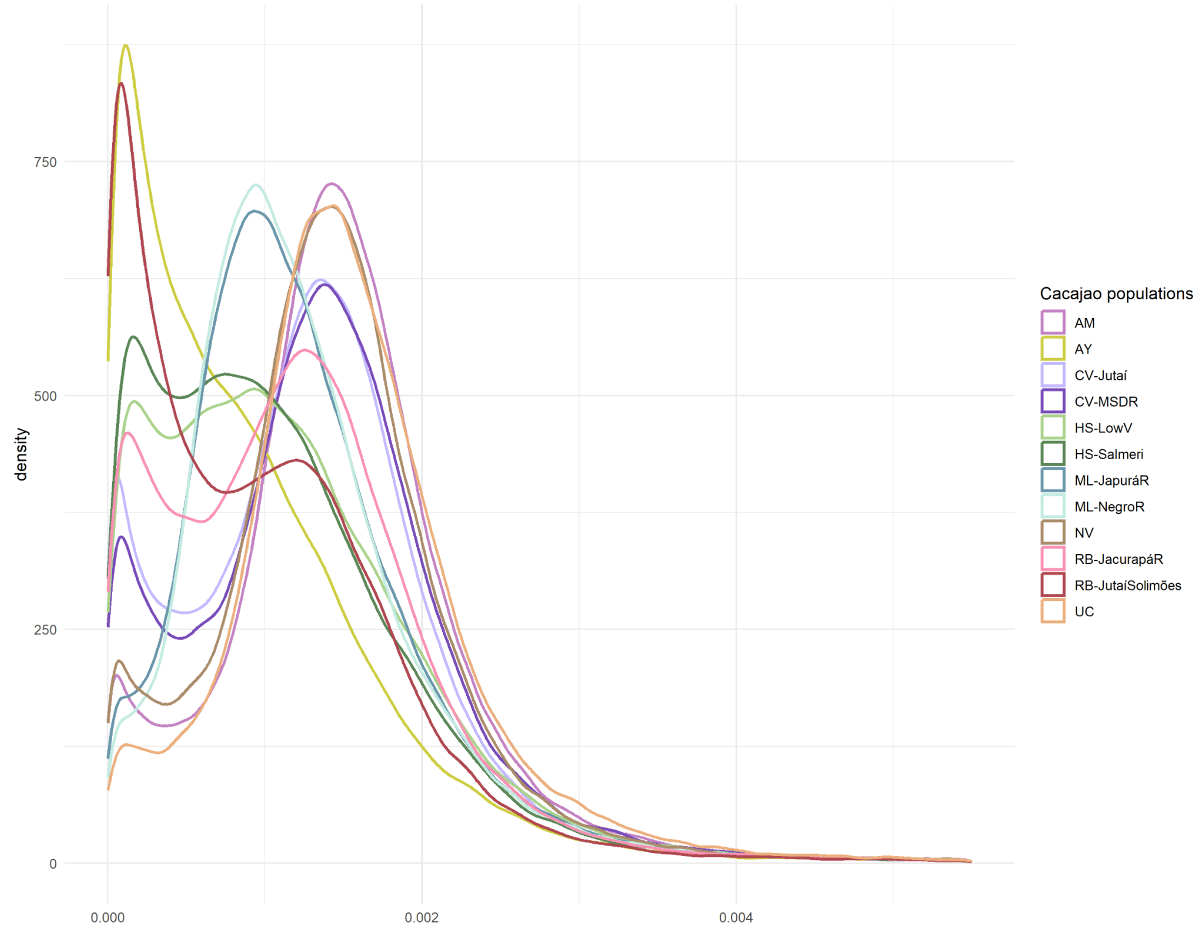


**Figure S3.** Heterozygous sites in 1kb window density distribution across the genome per *Cacajao* population. Note visible intra-species variability particularly between HS and RB populations. Overall, heterozygous distributions in bald uakaris show higher means than black's, in accordance with mean heterozygosity values in the groups.

##### 2.1.3. Structure analysis

In accordance with Principal Component Analyses, ADMIXTURE on the full dataset (Figure S4B) shows that population structure in the genus follows patterns described by pelage coloration: the largest differences being found between bald and black populations, followed by differences within each group ((bald*:* red bald uakaris - white bald uakaris), *(* black*:* ML - AY,+HS)).

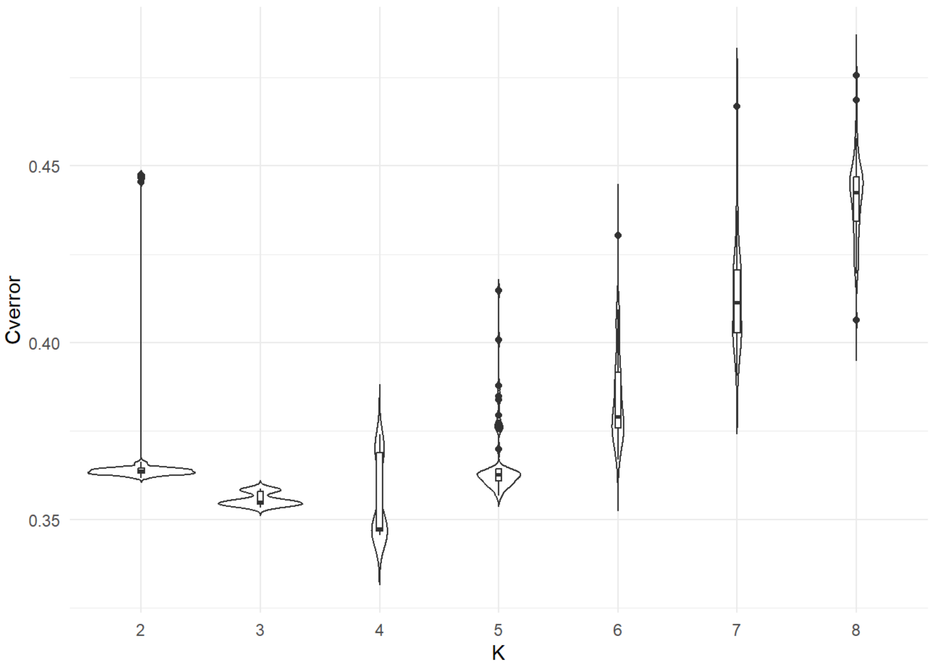


**Figure S4A.** ADMIXTURE cross-validation error for all *Cacajao* populations (n = 10.46M SNPs) over 50 random-seed replicates per K. Best $K = 4$.

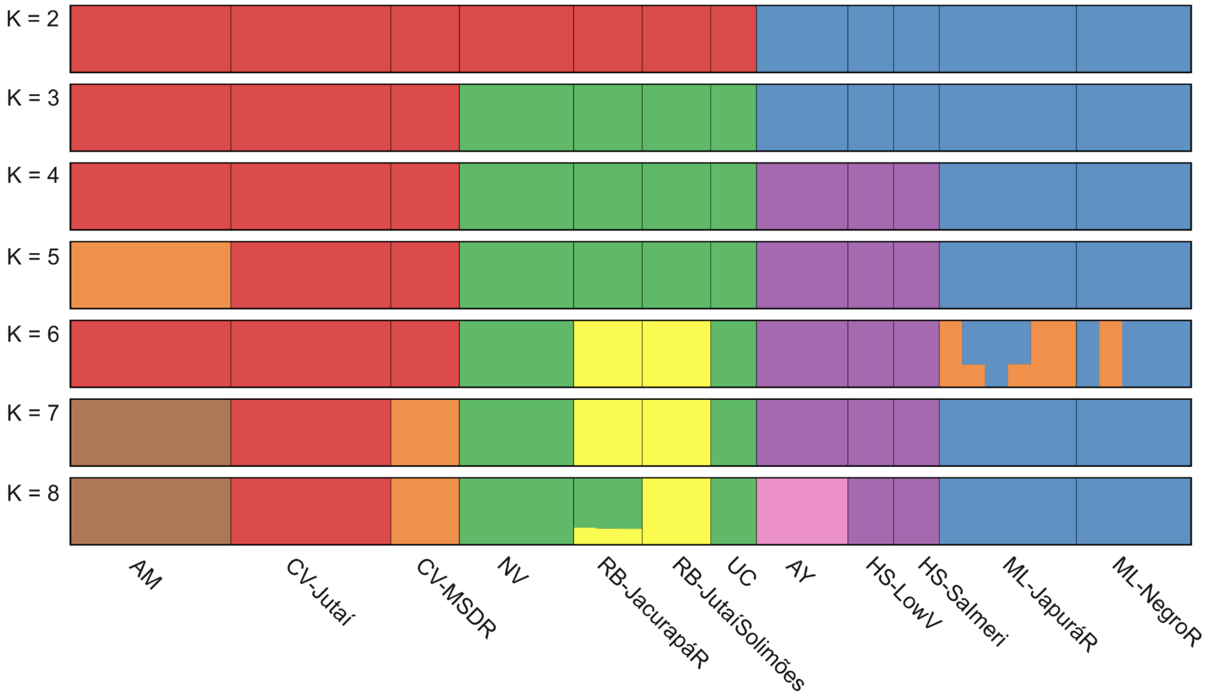


**Figure S4B.** ADMIXTURE proportions for all *Cacajao* populations (n = 10.46M SNPs) in the best run per K. Best K ($K = 4$). clusters white bald uakaris, red bald, northern Negro river black uakaris and southern populations independently.

In bald uakaris, the intra-group structure observed analyzing the whole dataset (Figure S4B) is maintained (Figure S5B). Results suggest CV-MSDR shares a small ancestry component with AM, we nonetheless propose that given that his signal was not captured by any other method, CV-MSDR represents a unique ancestral component on its own.

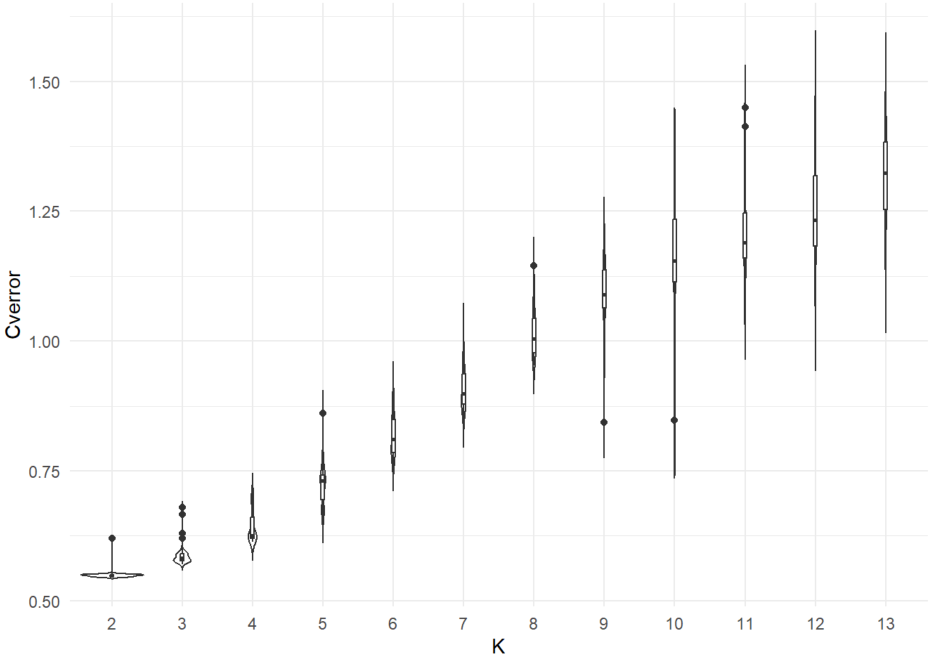


**Figure S5A.** ADMIXTURE cross-validation error for *Cacajao* bald populations (n = 6.10M SNPs) over 50 random-seed replicates per K. Best $K = 2$.

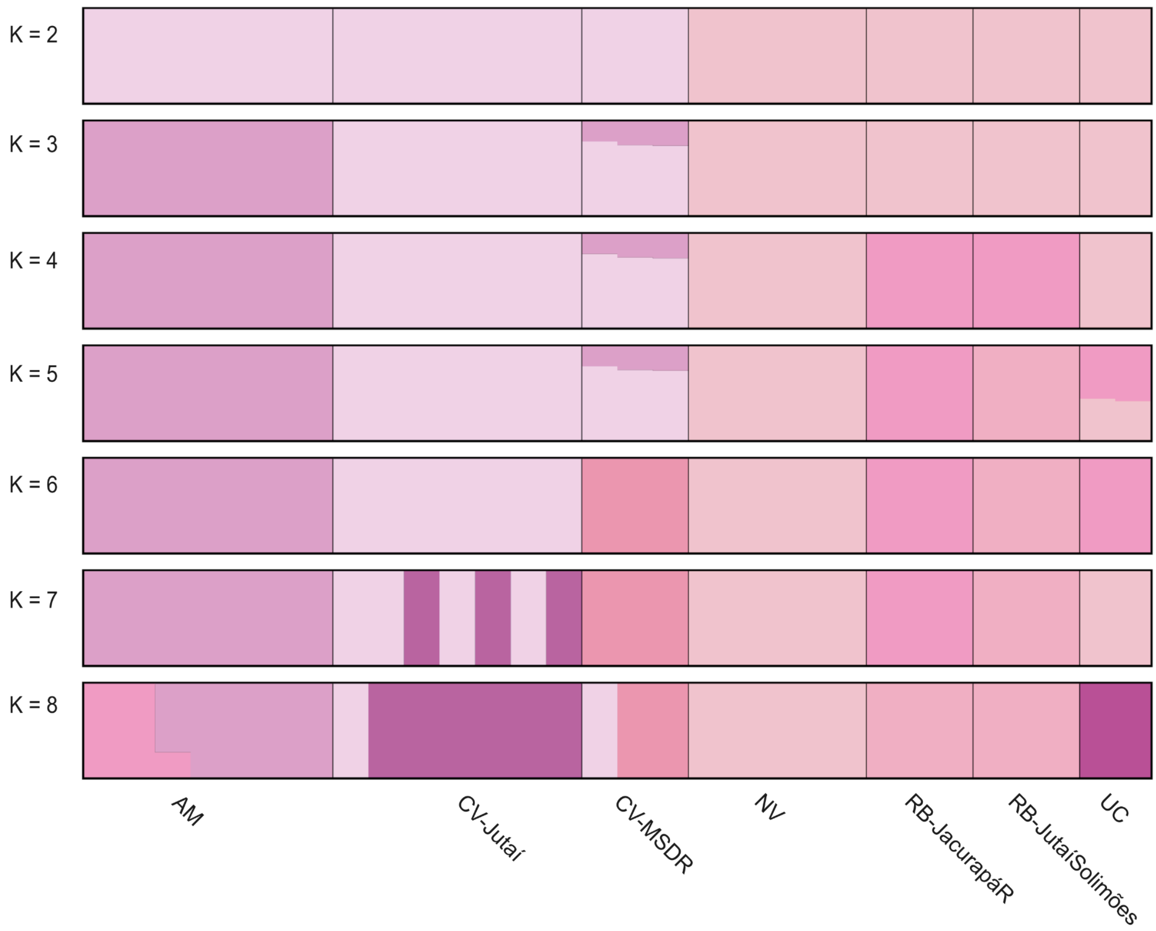


**Figure S5B.** ADMIXTURE proportions for *Cacajao* bald populations (n = 6.10M SNPs) in the best run per K. Best K ($K = 2$) clusters red bald uakaris and white bald uakaris independently.

As for bald uakaris, in black, the intra-group structure observed analyzing the whole dataset (Figure S4B) is maintained (Figure S6B) and no admixed individuals are identified.

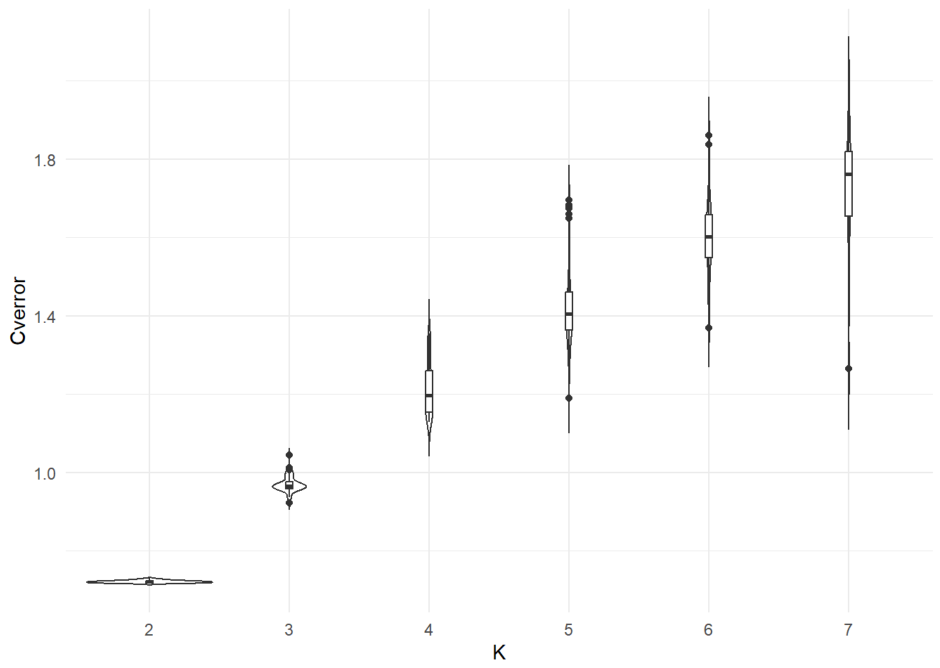


**Figure S6A.** ADMIXTURE cross-validation error for *Cacajao* black populations (n = 2.24M SNPs) over 50 random-seed replicates per K. Best $K = 2$.

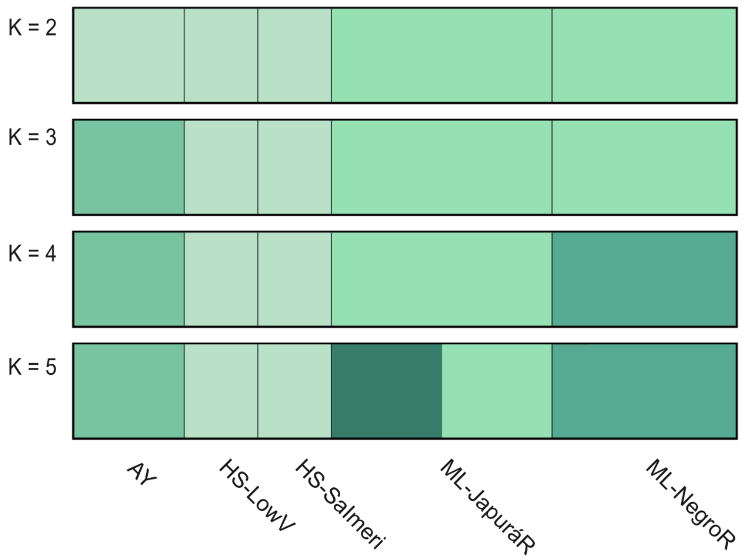


**Figure S6B.** ADMIXTURE proportions for *Cacajao* black populations (n = 2.24M SNPs) in the best run per K. Best K ($K = 2$) clusters northern Negro river black uakaris and southern populations independently.

##### 2.1.4. Principal Component Analysis

The final set employed to run STRUCTURE, PCA and EEMS included the pruning of regions in linkage disequilibrium (LD), after which we kept roughly between 30% - 20% of the variants. Nonetheless, PCA was run as well without this filter, thus observing that in contrast to PC1 which splits apart bald from black uakaris with both filtering sets, PC2 finds more variance in black uakaris without considering LD (Figure S6A), while this is the case for bald uakaris when LD is removed (Figure 1C). Therefore, based on Figure 1C nor Figure S6A one can not make any assumption on which intra-group variance is stronger. When the latter is analyzed (Figure S6B,C), no major differences are detected to the LD pruned dataset.

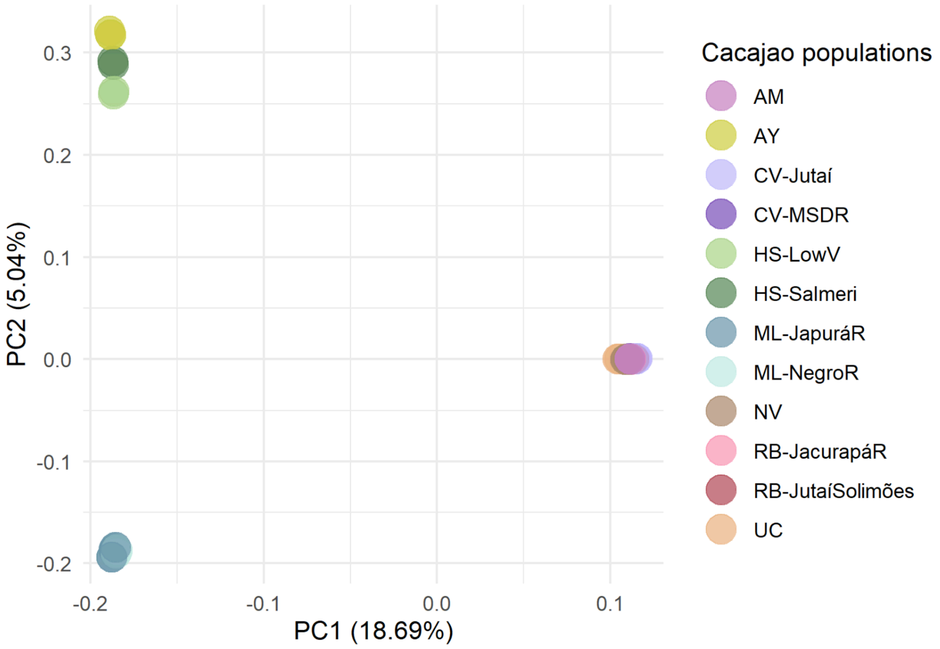


**Figure S7A.** Principal component analysis for all *Cacajao* populations (n = 33.1M SNPs).

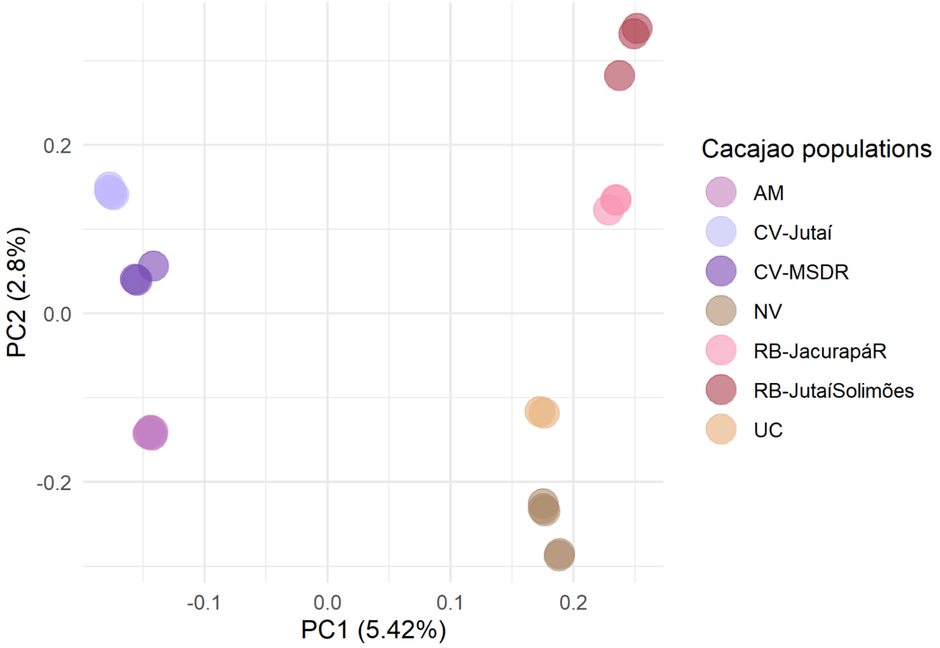


**Figure S7B.** Principal component analysis for *Cacajao* bald populations (n = 20.6M SNPs).

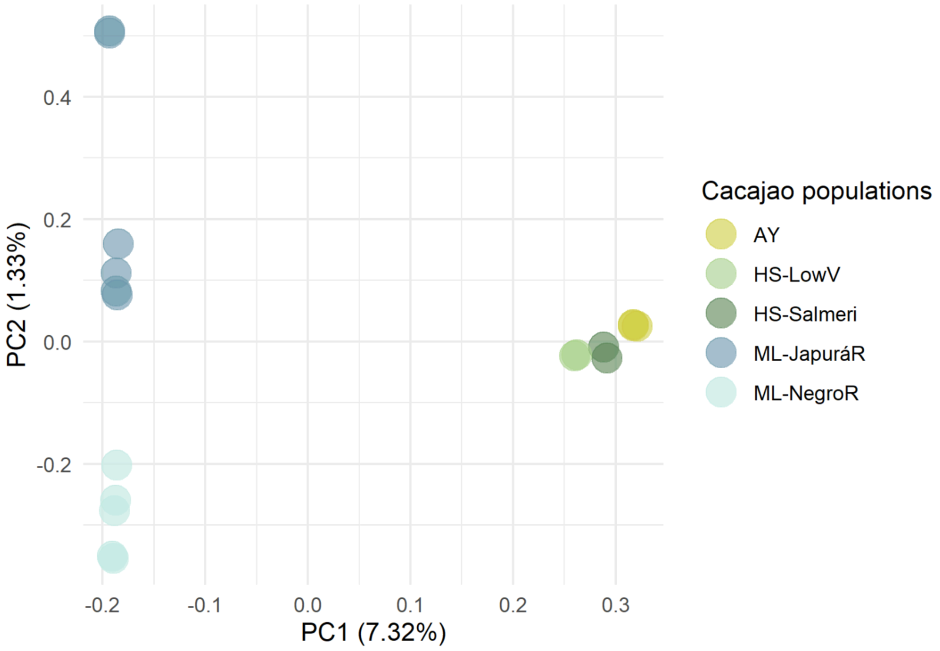


**Figure S7C.** Principal component analysis for *Cacajao* black populations (n = 12.4M SNPs).

#### 2.2. Phylogenetics

Two whole genome phylogenies were built, based on 250kb and 1Mb-long windows respectively and an additional one based on whole mitochondrial genomes. For the sake of visualization, two figures were generated from each of these, one depicting an ultrametric tree and the other a phylogram where branch lengths are proportional to the number of substitutions in the given branch.

##### 2.2.1. Whole genome - 250kb windows

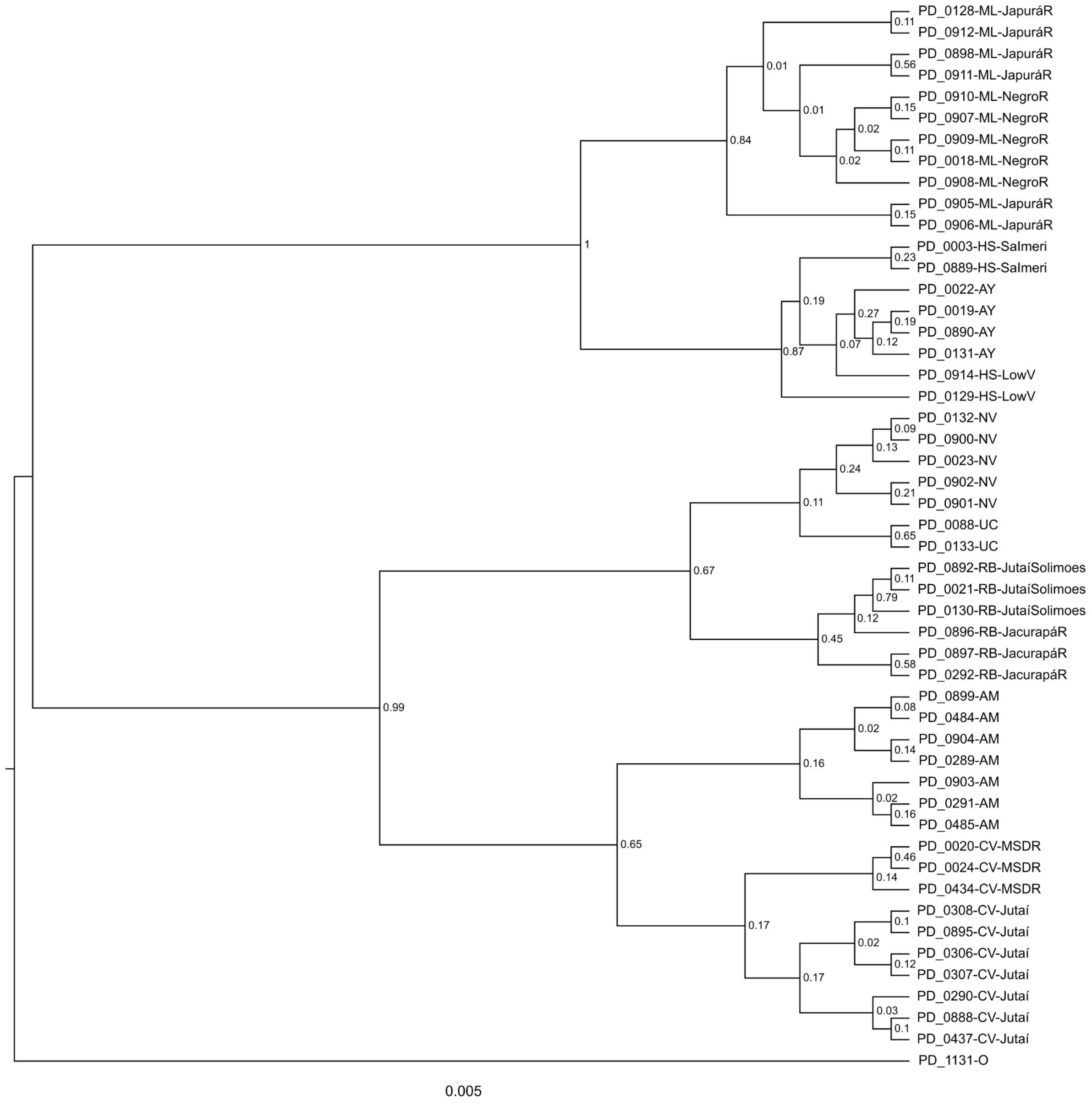


**Figure S8A** . Phylogenetic tree based on 250kb-windows phylogram

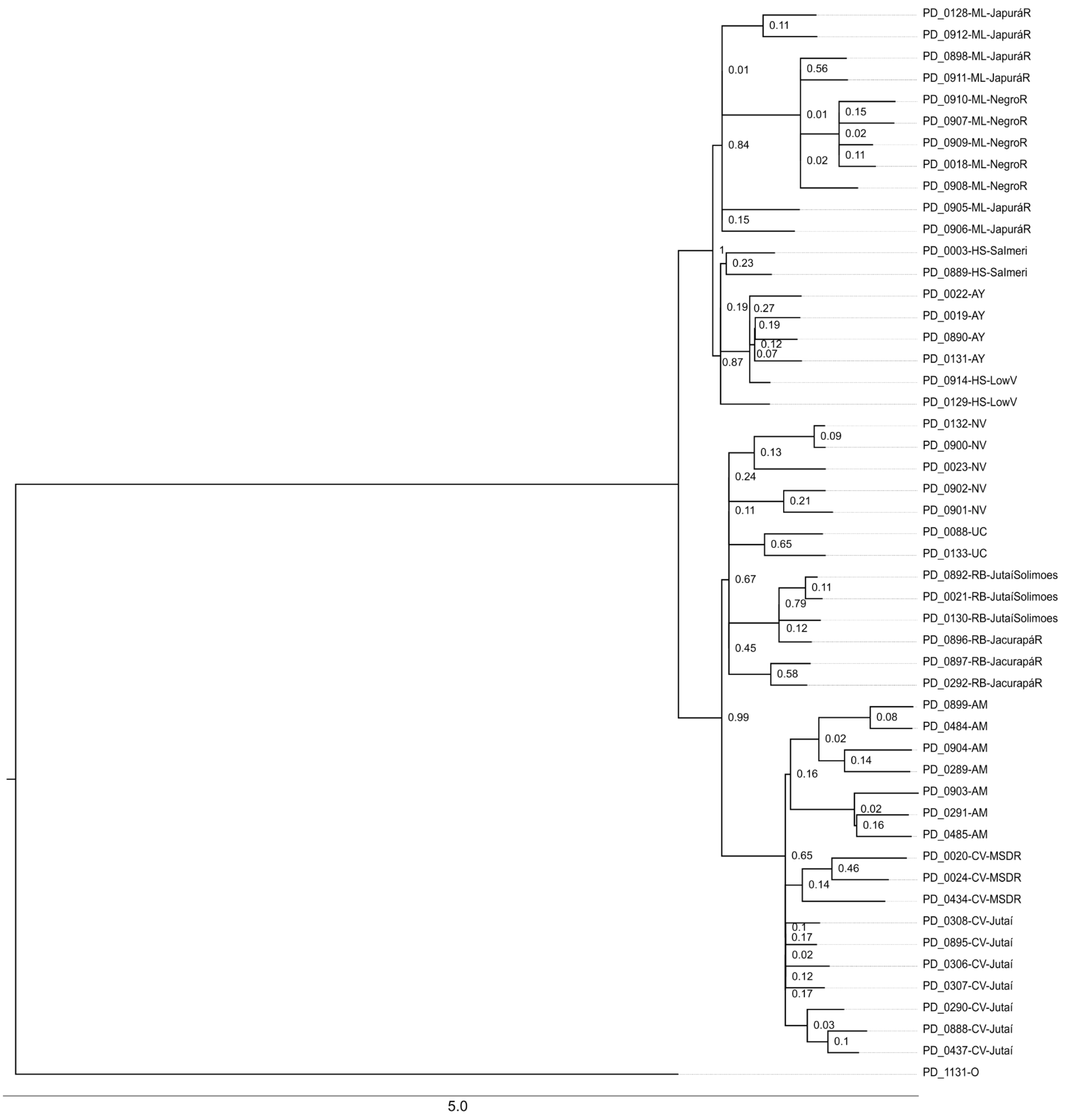


**Figure S8B.** Ultrametric phylogenetic tree based on 250kb-windows

#### 2.2.2. Whole genome - 1Mb windows

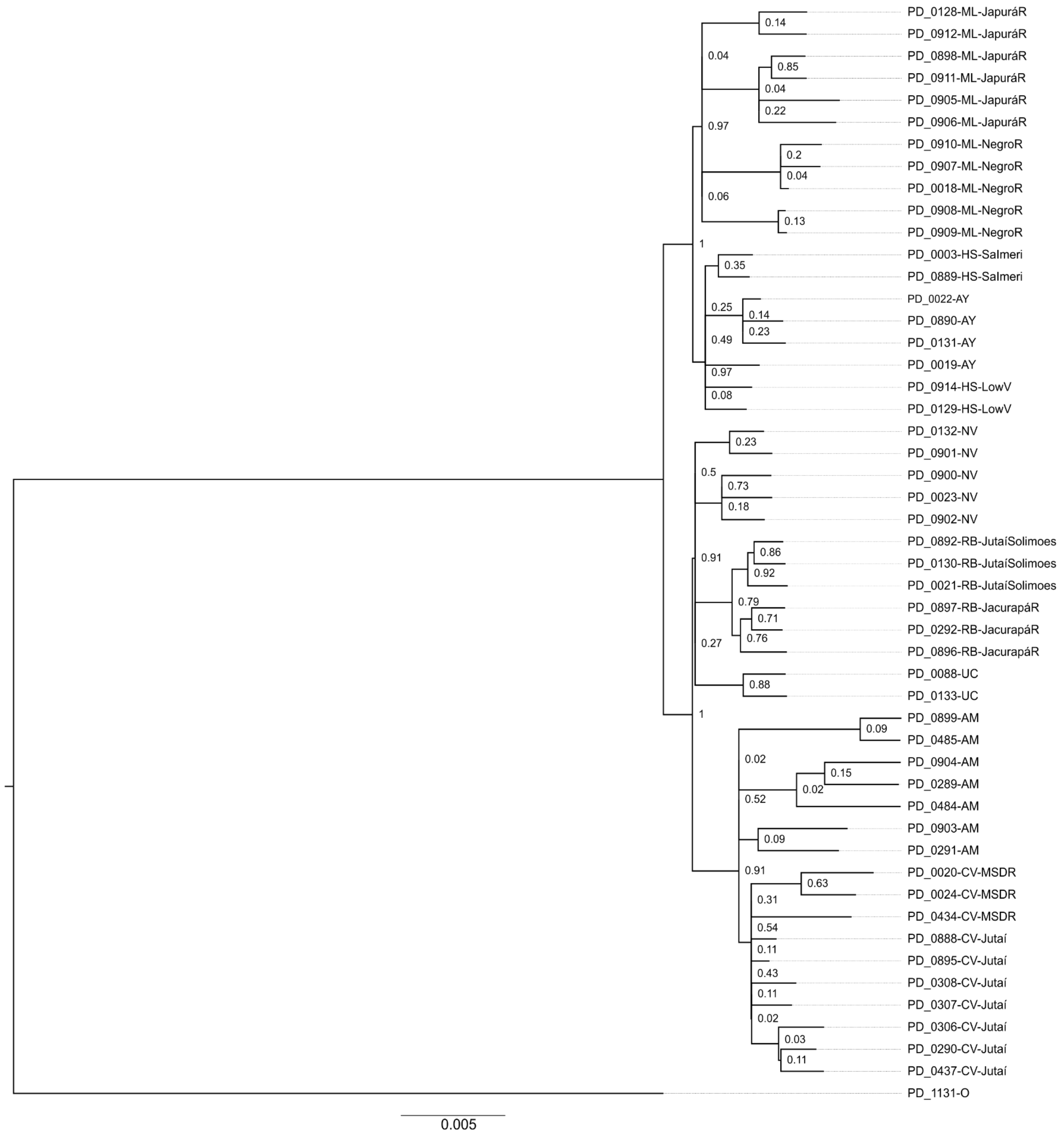


**Figure S9A.** Ultrametric phylogenetic tree based on 1Mb-windows

We calculated topological distances from each 1Mb window tree to consensus. Two strategies using *ape.dist.topo* function: *PH85* or *score.* The latter takes into account branch lengths. If these are not considered, a normal distribution is obtained where the mean goes towards a high differentiation from the consensus. On the other hand, considering branch lengths, we see a skewed normal distribution where most of the window trees are quite similar to the consensus while some of these show high differentiation. When considering the distance representations applying each of the metrics (Figure S9B1, S9C1), these have in common a few strong differentiation signals, for example that in the first window of the tarseq_28 scaffold, the tree 713. These outliers are not found contiguously in the genome, hence we propose these may correspond to naturally high variable regions (structural regions such as centromeric), thus not meaningful in this context. Overall, when the two metrics are compared the background pattern is conserved, describing a generally homogenous distance from individual window to consensus trees. Nonetheless, the intensity of the signals is reduced by considering branch length, likely by removing background noise.

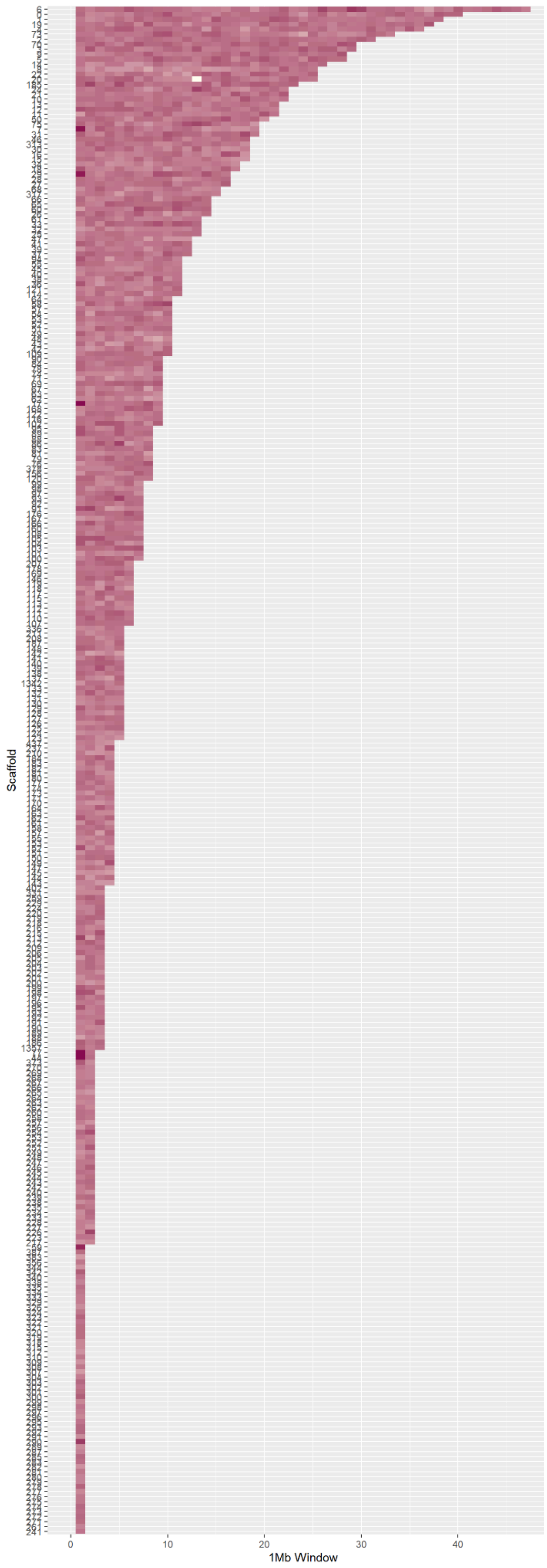


**Figure S9B1.** Topological distances between consensus tree and each 2144 1Mb-based tree not considering branch lengths and not-normalized (d_PH85)

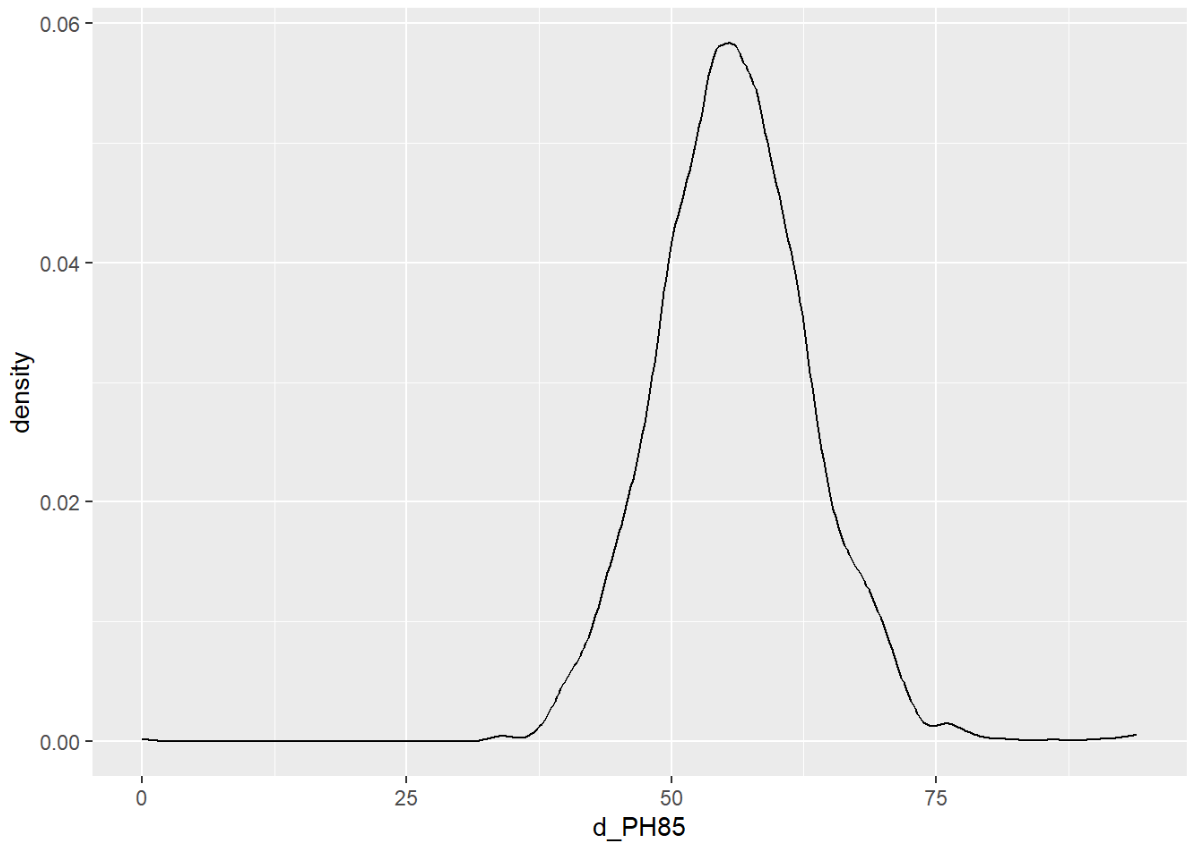


**Figure S9B2.** Density distribution of d_PH85 statistic which evaluates topological distance not considering branch length.

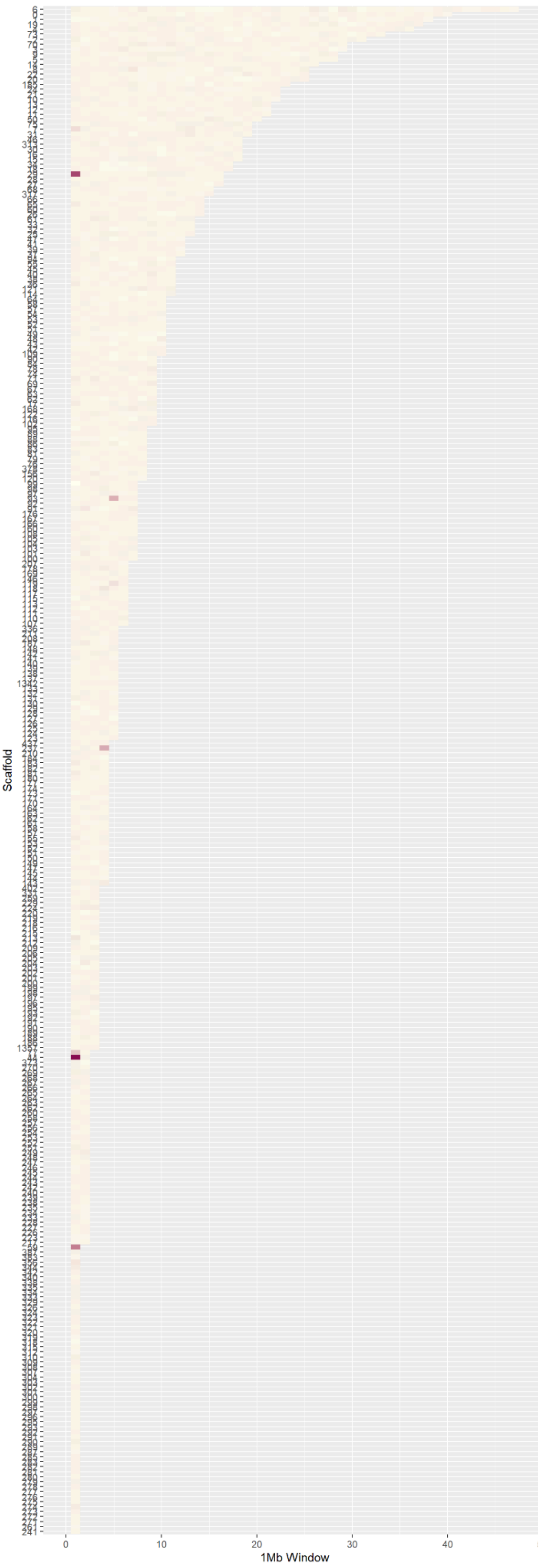


**Figure S9C1.** Topological distances between consensus tree and each 2144 1Mb-based tree considering branch lengths and not-normalized (score)

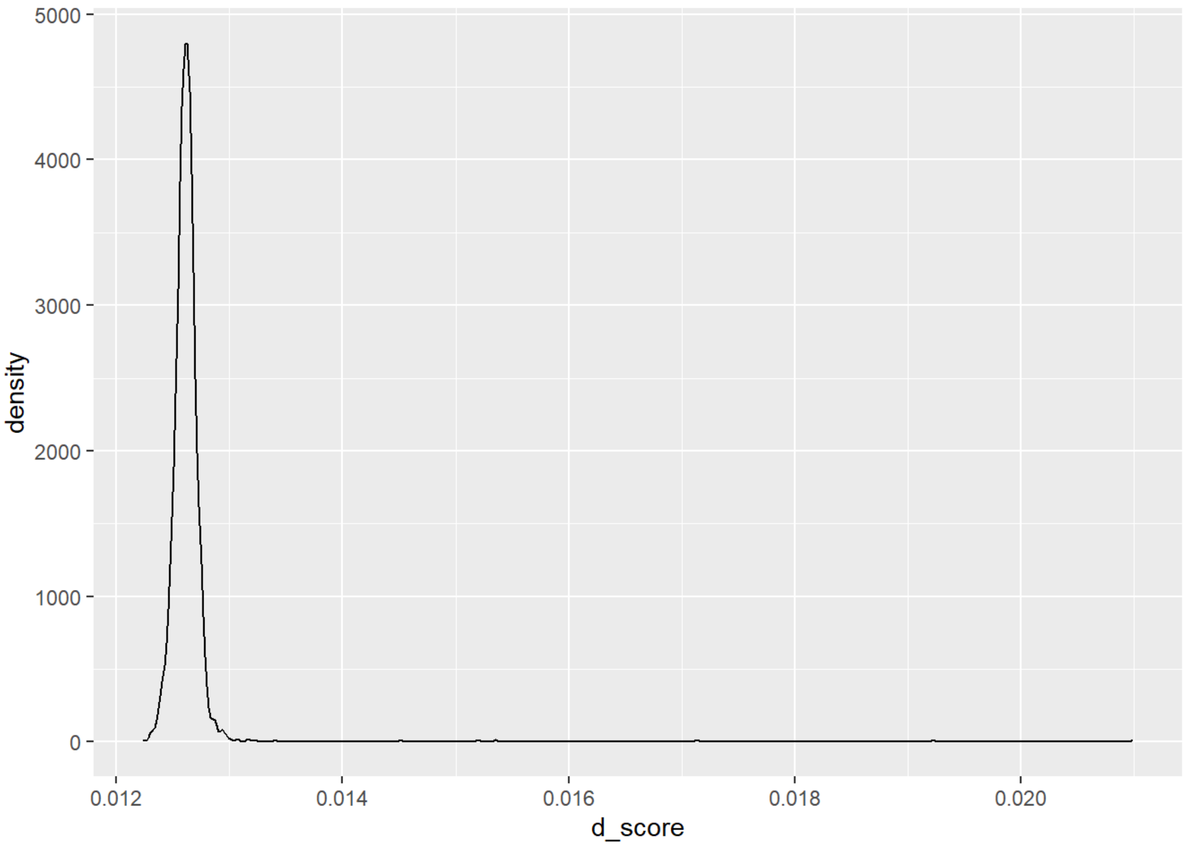


**Figure S9C2.** Density distribution of score statistic which evaluates topological distance considering branch length, which is removing the background noise in contrast to PH85, only keeping differentiation signals. Most of the windows show very close topologies to the consensus tree and few outliers can be seen in the right tail of the distribution.

#### 2.2.3. Whole mitochondrial genome

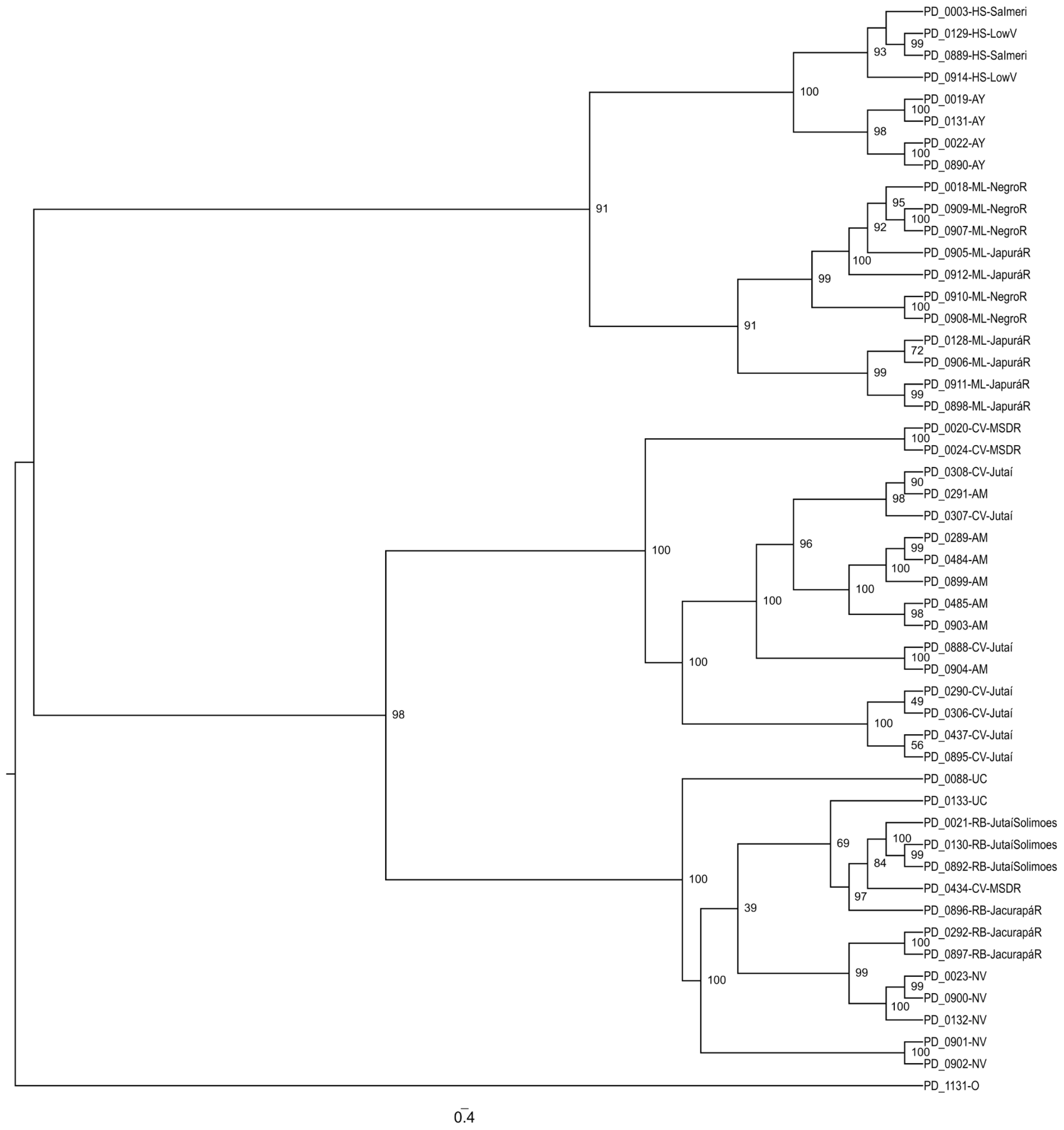


**Figure S10A.** Whole mitochondrial genomes-based phylogeny phylogram

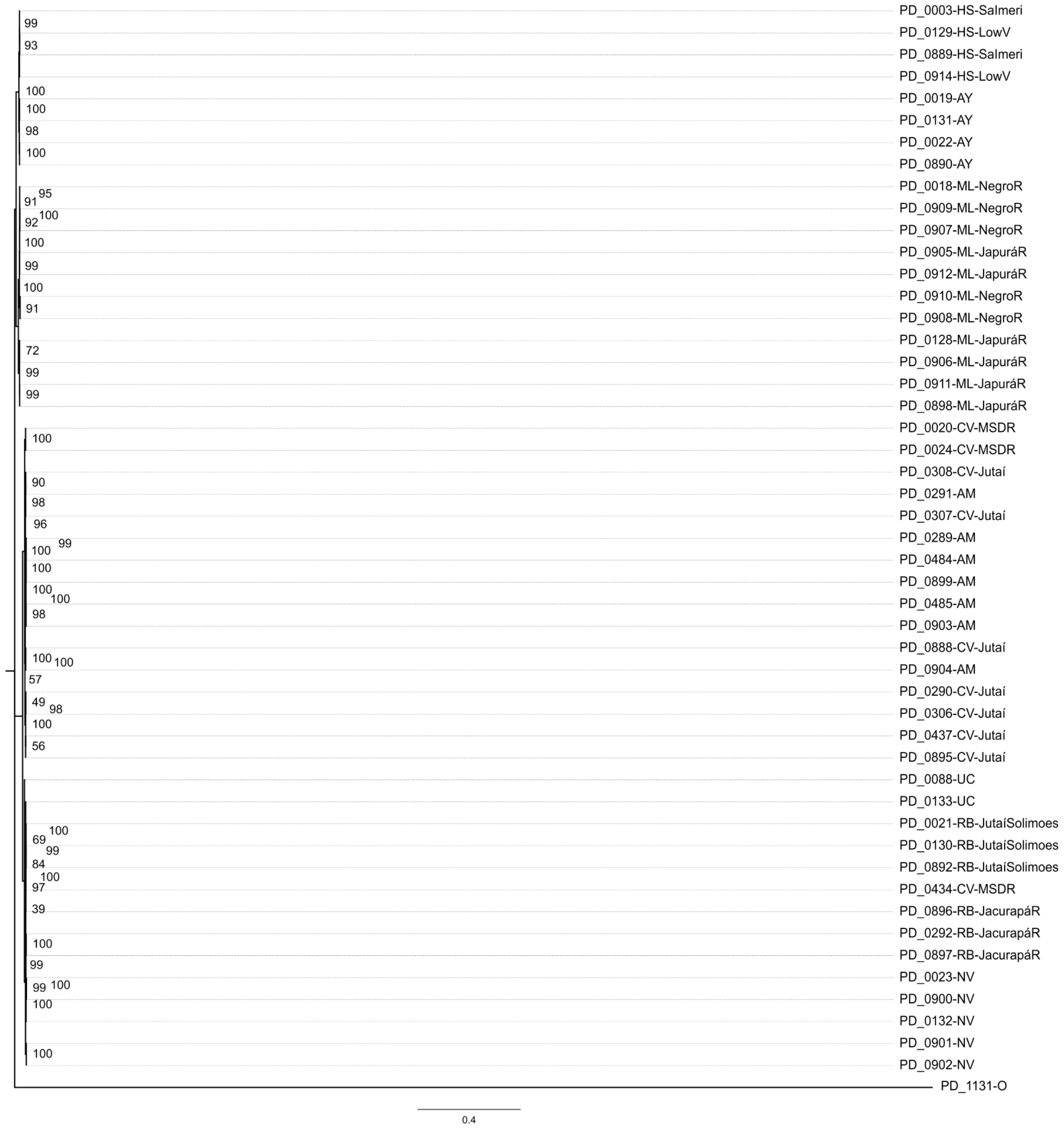


**Figure S10B.** Ultrametric whole mitochondrial genomes-based phylogeny

#### 2.3. Gene flow

##### 2.3.1. Fst

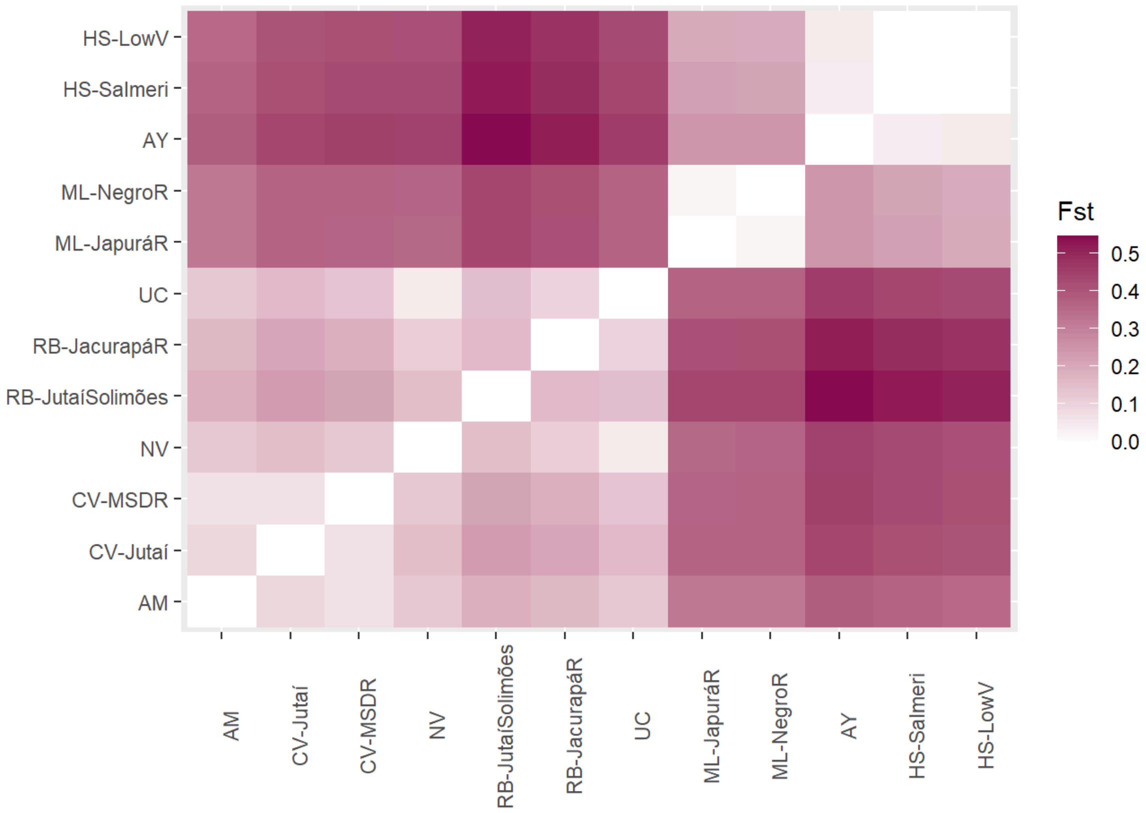


**Figure S11.** Average Hudson Fst across the whole genome as a proxy of genetic dissimilarity between *Cacajao* populations. Note clear dissimilarity between bald and black sample clusters, as well as intra-group structure (bald: red/white, black: north/south Negro River).

#### 2.3.2. EEMS

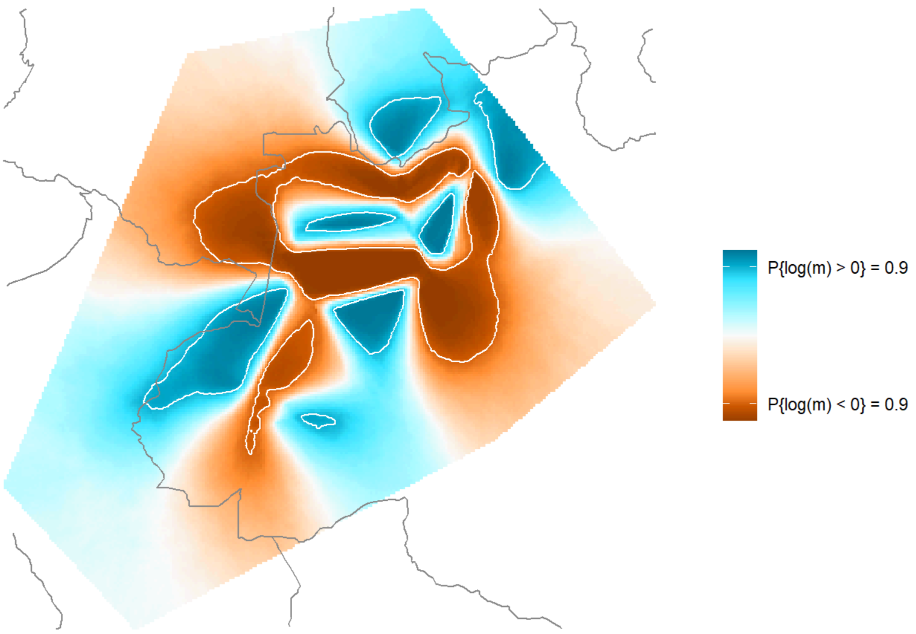


**Figure S12A**. Map of probability of estimated effective migration rates (m) across all studied *Cacajao* populations' habitat distribution by EEMS. Higher intensity indicates higher probability.

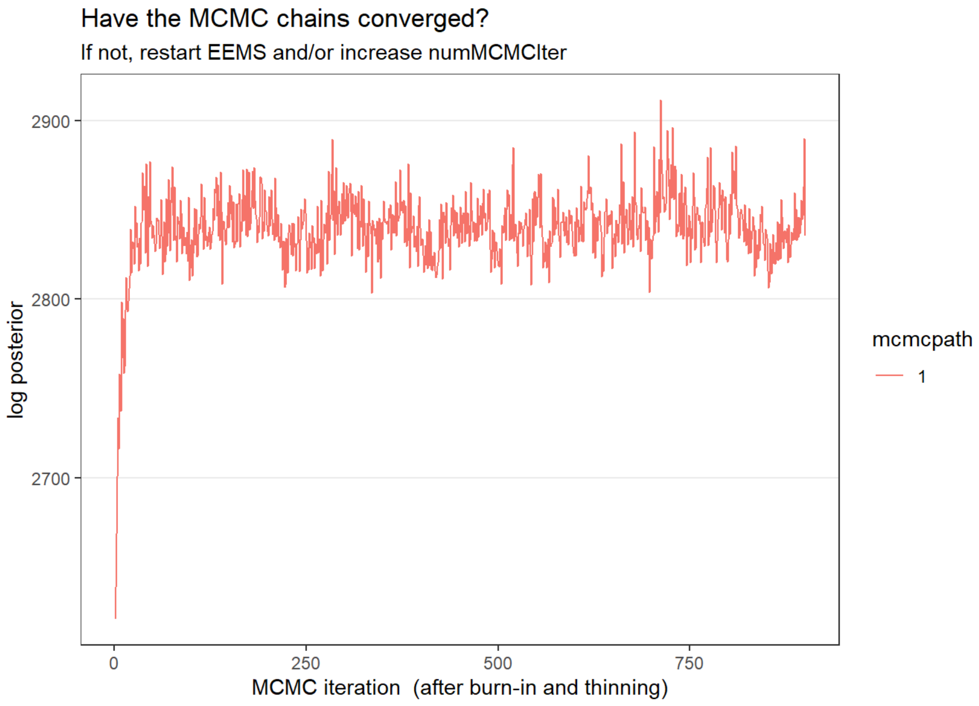


**Figure S12B.** Monte Carlo Markov Chain convergence assessment of the expectation-maximization EEMS algorithm.

#### 2.4. Demographic inference

##### 2.4.1. fastsimcoal2

| **Parameter** | **Description** | **Lower CI 95%** | **Best Estimate** | **Upper CI 95%** | **Model** |
|---|---|---|---|---|---|
| BALD | *Ne* Ancestor bald | 54970.46 | 56410.51 | 59382.99 | Bald and black |
| BLACK | *Ne* Ancestor black | 53228.77 | 55556.04 | 57783.41 | Bald and black |
| CACA | *Ne* Ancestor *Cacajao* | 152067.11 | 160745.1 | 167113.82 | Bald and black |
| TsCac | Split time *Cacajao* | 885690.29 | 922911.28 | 976642.21 | Bald and black |
| AMUNA | *Ne* AM | 108750.31 | 112983 | 119996.30 | Bald |
| BALD | *Ne* Ancestor bald | 83500.93 | 86749.92 | 89531.44 | Bald |
| CALVUS | *Ne* CV | 57298.12 | 60047.79 | 62426.66 | Bald |
| NOVAESI | *Ne* NV | 109791.67 | 114439.3 | 117930.79 | Bald |
| NV_AM | Migration rate NV - AM | 2.29E-06 | 2.80E-06 | 2.93E-06 | Bald |
| TsBald | Split time bald | 640065.44 | 665896.67 | 681795.10 | Bald |
| TsWhite | Split time white bald | 165112.08 | 166506.97 | 178077.88 | Bald |
| WHITE | *Ne* Ancestor white bald | 229891.36 | 236557 | 260214.51 | Bald |
| BLACK | *Ne* Ancestor black | 34633.58 | 45362.05 | 474148.07 | Black |
| HOSOMI | *Ne* HS | 20768.29 | 33889.63 | 34675.02 | Black |
| HS_ML | Migration rate HS - ML | 7.95E-06 | 8.27E-06 | 2.91E-04 | Black |
| Japurá | *Ne* ML-JapuráR | 40048.57 | 56237.05 | 61288.83 | Black |
| MELANO | *Ne* ML | 2455.56 | 41648.81 | 43813.32 | Black |
| Negro | *Ne* ML-NegroR population | 46309.28 | 73299.26 | 80189.77 | Black |

| | | | | | |
|---|---|---|---|---|---|
| Ng_Ja | Migration rate ML-NegroR - ML-JapuráR | 2.43E-06 | 3.19E-06 | 1.17E-04 | Black |
| TsBlack | Split time black | 273616.95 | 662471.94 | 826289.57 | Black |
| TsML | Split time ML | 29122.42 | 31167.26 | 152844.95 | Black |

**Table S6.** Maximum likelihood estimates and confidence intervals obtained in three independent model topologies ((i) bald and black, ii) bald and iii) black) with fastsimcoal2 (See Supplementary Methods (1.3.1)).

##### 2.4.2. SMC++ demographic inference

The following SMC++ analysis was run to i) investigate the trajectory of the effective population sizes (*Ne*) of different groups of species in the genus in relation to current heterozygosity patterns, and ii) to compare and support the constant *Ne* and split time estimates obtained in the Maximum Likelihood approach.

We found the results of the two methods overall agree: in both methods, bald uakaris' split is identified to be older than black's and present higher *Ne* estimates too; the main disagreement appears at the intragroup level with white and red bald uakaris' *Ne*. Differences are nonetheless to be expected, different parameters are being estimated (constant *Ne* vs. *Ne* trajectory) through completely different approaches - moreover, intra-group estimates are based on a smaller number of samples, from which higher variance can be expected.

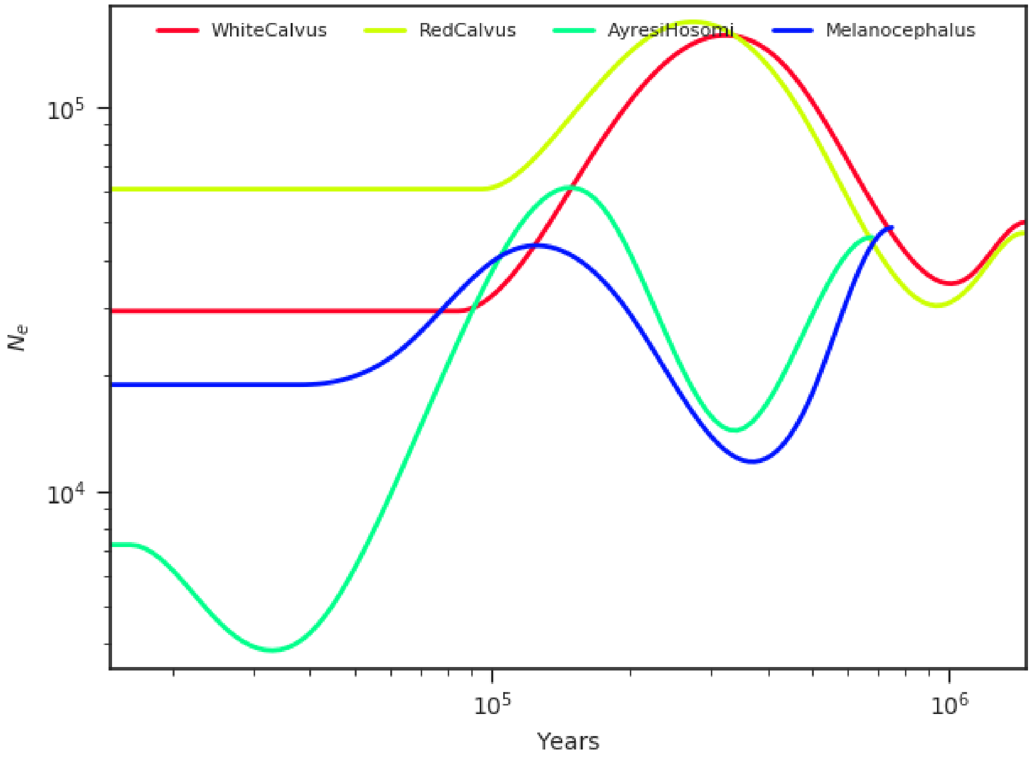


**Figure S13.** SMC++ output of *Ne* trajectories in the last million years (approximately estimated divergence time between bald and black uakaris by maximum likelihood (ML)) modeling. The same four groups of samples used in the ML estimation of divergence times and effective population sizes were used. The results show approximately consistent divergence times compared to the ML approach. Also, the order of *Ne* estimates' size is consistent among analyses, and it changes in accordance to major split times identified by ML.

#### 2.5. Genetic differentiation

See table with genes with differentially fixed non-synonymous variants passing all applied filters (See Methods, “Fst - Coding non-synonymous variation” section): 336 genes for bald uakaris, 406 for black and 98 for genes affected in both groups by different variants (Supplementary Table S2). The three gene lists were inspected independently for functional overrepresentation with the following results. In the following figures % depicts category enrichment and bar color, significance: dark blue indicates significance (FDR <= 0.05) while light blue refers to non-significance.

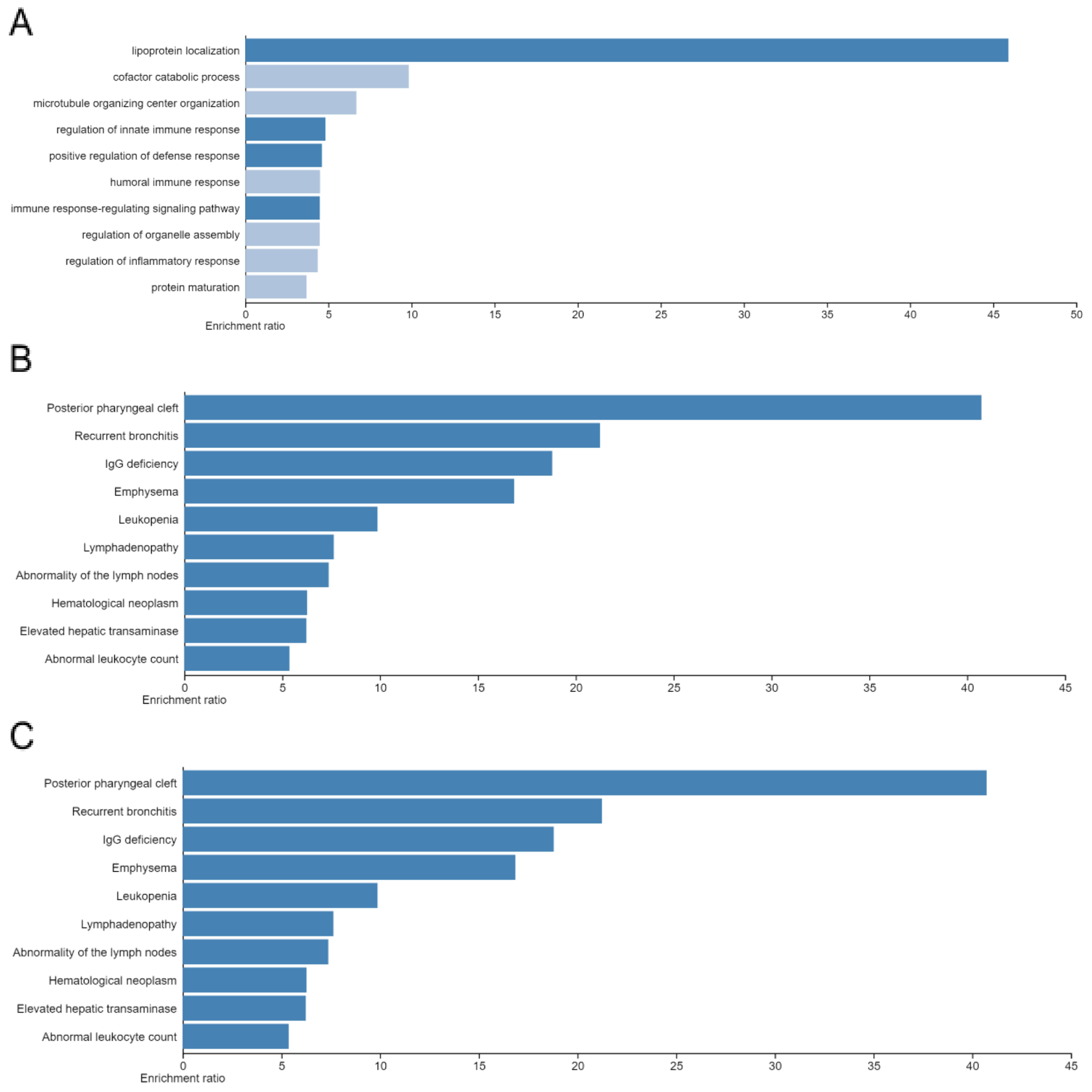


**Figure S14.** Enrichment of 98 genes affected in both groups by different non-synonymous variants. Results for **A)** GO (Biological Process non-redundant), **B)** Human Ontology Phenotype and **C)** OMIM.

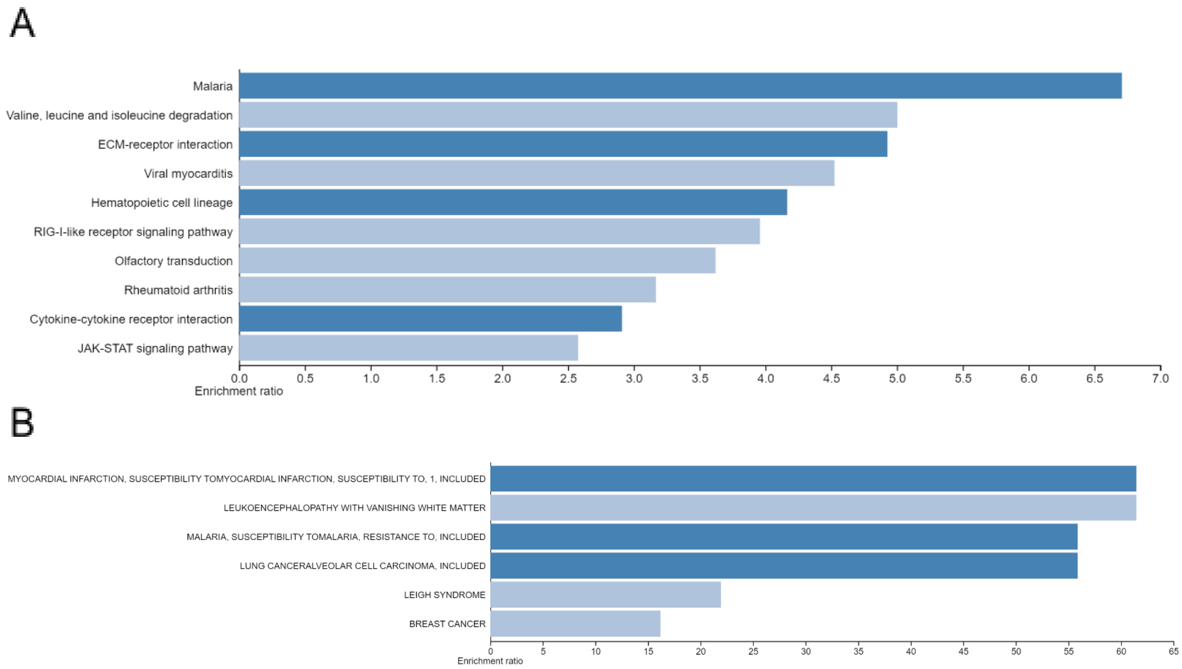


**Figure S15.** Enrichment of 336 genes with non-synonymous variants only in bald uakaris. Results for **A)** KEGG Pathway, **B)** OMIM.

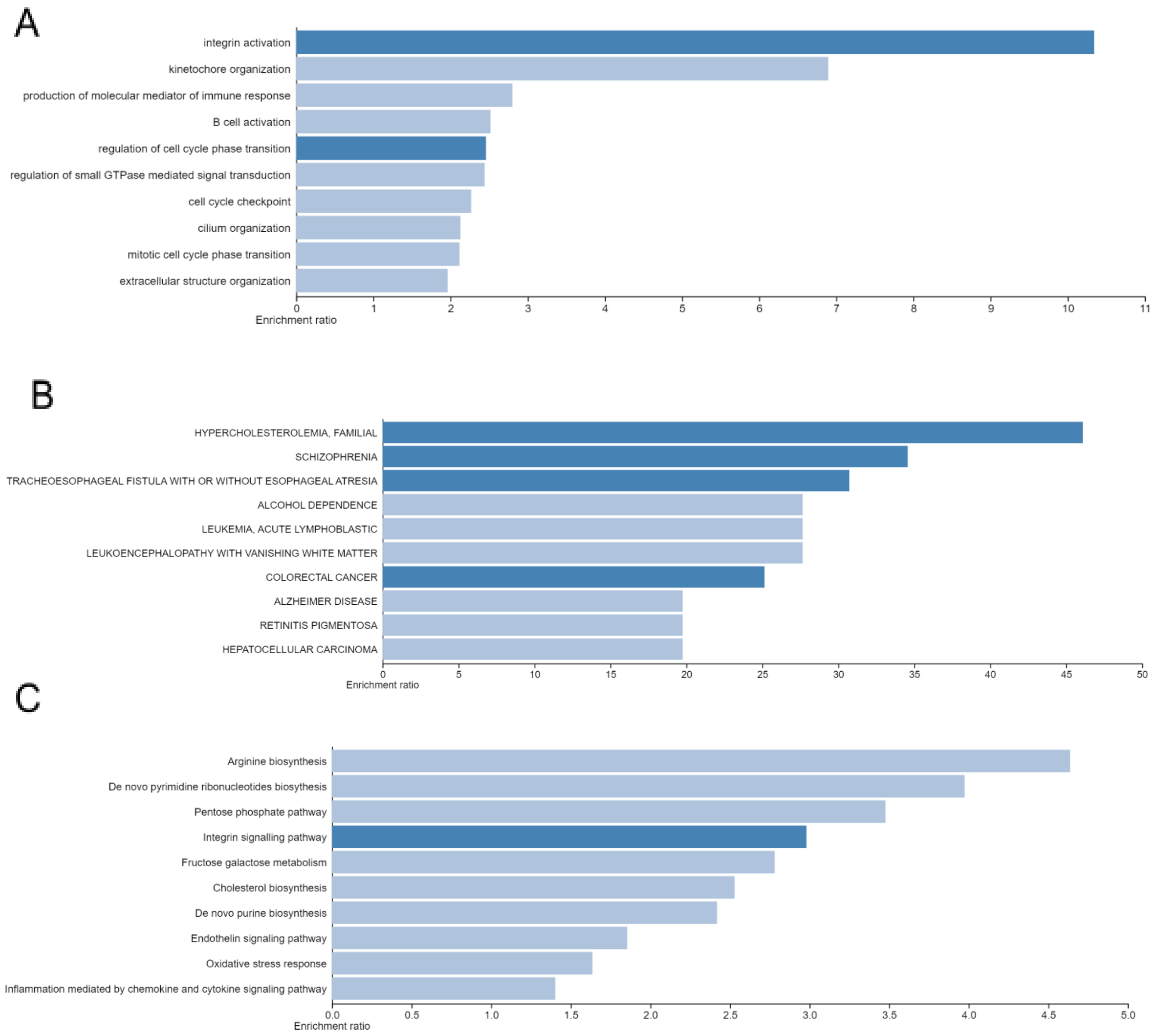


**Figure S16.** Enrichment of 406 genes with non-synonymous variants only in black uakaris. Results for **A)** GO (Biological Process non-redundant), **B)** OMIM, **C)** PANTHER Pathway.
