## Supplementary Table S1: Sample metadata for "Whole genomes of the amazonian *Cacajao* reveal complex connectivity and fast differentiation driven by high environmental dynamism"

| ID | Genus | Species | Population | Group | Coverage_mode |
|---|---|---|---|---|---|
| PD_0003 | *Cacajao* | *hosomi* | HS-Salmeri | Black | 30 |
| PD_0018 | *Cacajao* | *melanocephalus* | ML-NegroR | Black | 33 |
| PD_0019 | *Cacajao* | *ayresi* | AY | Black | 34 |
| PD_0020 | *Cacajao* | *calvus* | CV-MSDR | Bald | 32 |
| PD_0021 | *Cacajao* | *rubicundus* | RB-JutaíSolimões | Bald | 34 |
| PD_0022 | *Cacajao* | *ayresi* | AY | Black | 34 |
| PD_0023 | *Cacajao* | *novaesi* | NV | Bald | 37 |
| PD_0024 | *Cacajao* | *calvus* | CV-MSDR | Bald | 33 |
| PD_0088 | *Cacajao* | *ucayalii* | UC | Bald | 38 |
| PD_0128 | *Cacajao* | *melanocephalus* | ML-JapuráR | Black | 35 |
| PD_0129 | *Cacajao* | *hosomi* | HS-LowV | Black | 31 |
| PD_0130 | *Cacajao* | *rubicundus* | RB-JutaíSolimões | Bald | 35 |
| PD_0131 | *Cacajao* | *ayresi* | AY | Black | 35 |
| PD_0132 | *Cacajao* | *novaesi* | NV | Bald | 39 |
| PD_0133 | *Cacajao* | *ucayalii* | UC | Bald | 32 |
| PD_0289 | *Cacajao* | *amuna* | AM | Bald | 38 |
| PD_0290 | *Cacajao* | *calvus* | CV-Jutaí | Bald | 36 |
| PD_0291 | *Cacajao* | *amuna* | AM | Bald | 39 |
| PD_0292 | *Cacajao* | *rubicundus* | RB-JacurapáR | Bald | 36 |
| PD_0306 | *Cacajao* | *calvus* | CV-Jutaí | Bald | 23 |
| PD_0307 | *Cacajao* | *calvus* | CV-Jutaí | Bald | 30 |
| PD_0308 | *Cacajao* | *calvus* | CV-Jutaí | Bald | 26 |
| PD_0434 | *Cacajao* | *calvus* | CV-MSDR | Bald | 34 |
| PD_0437 | *Cacajao* | *calvus* | CV-Jutaí | Bald | 43 |
| PD_0484 | *Cacajao* | *amuna* | AM | Bald | 31 |
| PD_0485 | *Cacajao* | *amuna* | AM | Bald | 28 |
| PD_0888 | *Cacajao* | *calvus* | CV-Jutaí | Bald | 26 |
| PD_0889 | *Cacajao* | *hosomi* | HS-Salmeri | Black | 29 |
| PD_0892 | *Cacajao* | *rubicundus* | RB-JutaíSolimões | Bald | 30 |
| PD_0895 | *Cacajao* | *calvus* | CV-Jutaí | Bald | 29 |
| PD_0896 | *Cacajao* | *rubicundus* | RB-JacurapáR | Bald | 26 |
| PD_0897 | *Cacajao* | *rubicundus* | RB-JacurapáR | Bald | 29 |
| PD_0898 | *Cacajao* | *melanocephalus* | ML-JapuráR | Black | 28 |
| PD_0899 | *Cacajao* | *amuna* | AM | Bald | 29 |
| PD_0900 | *Cacajao* | *novaesi* | NV | Bald | 28 |
| PD_0901 | *Cacajao* | *novaesi* | NV | Bald | 24 |
| PD_0902 | *Cacajao* | *novaesi* | NV | Bald | 29 |
| PD_0903 | *Cacajao* | *amuna* | AM | Bald | 27 |
| PD_0904 | *Cacajao* | *amuna* | AM | Bald | 24 |
| PD_0905 | *Cacajao* | *melanocephalus* | ML-JapuráR | Black | 27 |
| PD_0906 | *Cacajao* | *melanocephalus* | ML-JapuráR | Black | 22 |
| PD_0907 | *Cacajao* | *melanocephalus* | ML-NegroR | Black | 31 |
| PD_0908 | *Cacajao* | *melanocephalus* | ML-NegroR | Black | 28 |
| PD_0909 | *Cacajao* | *melanocephalus* | ML-NegroR | Black | 28 |
| PD_0910 | *Cacajao* | *melanocephalus* | ML-NegroR | Black | 28 |
| PD_0911 | *Cacajao* | *melanocephalus* | ML-JapuráR | Black | 28 |
| PD_0912 | *Cacajao* | *melanocephalus* | ML-JapuráR | Black | 31 |
| PD_0914 | *Cacajao* | *hosomi* | HS-LowV | Black | 29 |
| PD_1131 | *Pithecia* | *pithecia* | O | Outgroup | NA |

| Heterozygosity_median | Inbreeding_theta | Longitude | Latitude |
|---|---|---|---|
| 0.0017 | 0 | -65.270 | 0.490 |
| 0.0019 | 0 | -64.5 | -2.5 |
| 0.0014 | 0 | -62.91 | -0.54 |
| 0.0020 | 0 | -64.935 | -2.912 |
| 0.0016 | 0 | -67.423 | -3.201 |
| 0.0015 | 0 | -62.95 | -0.38 |
| 0.0020 | 0.2123 | -70.196 | -6.864 |
| 0.0020 | 0 | -64.854 | -3.071 |
| 0.0020 | 0 | -73.668 | -7.461 |
| 0.0017 | 0 | -67.605 | -1.777 |
| 0.0018 | 0 | -66.417 | 1.135 |
| 0.0016 | 0 | -67.423 | -3.201 |
| 0.0015 | 0 | -62.95 | -0.38 |
| 0.0020 | 0 | -70.1958 | -6.8643 |
| 0.0022 | 0 | -73.668 | -7.461 |
| 0.0020 | 0 | -69.738 | -6.935 |
| 0.0018 | 0 | -67.395 | -3.313 |
| 0.0021 | 0 | -69.132 | -7.605 |
| 0.0017 | 0 | -68.618 | -3.237 |
| 0.0020 | 0 | -67.374 | -3.3 |
| 0.0018 | 0 | -67.137 | -3.298 |
| 0.0020 | 0 | -67.45 | -3.771 |
| 0.0018 | 0 | -65.333 | -2.411 |
| 0.0018 | 0 | -67.46 | -3.79 |
| 0.0021 | 0 | -69.132 | -7.605 |
| 0.0021 | 0 | -69.132 | -7.605 |
| 0.0020 | 0 | -71.36 | -8.83 |
| 0.0016 | 0 | -65.28 | 0.490 |
| 0.0015 | 0 | -67.423 | -3.201 |
| 0.0015 | 0 | -67.15 | -3.06 |
| 0.0019 | 0 | -68.618 | -3.237 |
| 0.0017 | 0 | -68.618 | -3.237 |
| 0.0018 | 0 | -69.2 | -1.69 |
| 0.0020 | 0 | -69.925 | -6.753 |
| 0.0020 | 0 | -70.196 | -6.864 |
| 0.0021 | 0 | -70.196 | -6.864 |
| 0.0020 | 0 | -69.925 | -6.753 |
| 0.0023 | 0 | -69.667 | -6.671 |
| 0.0021 | 0 | -69.667 | -6.671 |
| 0.0020 | 0 | -67.605 | -1.777 |
| 0.0019 | 0 | -67.605 | -1.777 |
| 0.0018 | 0 | -64.740 | -0.490 |
| 0.0018 | 0 | -64.650 | -0.490 |
| 0.0019 | 0 | -64.930 | -0.620 |
| 0.0019 | 0 | -64.910 | -0.580 |
| 0.0018 | 0 | -69.200 | -1.690 |
| 0.0018 | 0 | -69.340 | -1.660 |
| 0.0017 | 0 | -66.417 | 1.135 |
| NA | NA | NA | NA |
