## Supplementary Table S2: Sets of genes showing non-synonymous (NS) variation considered for ORA analyses. for "Whole genomes of the amazonian *Cacajao* reveal complex connectivity and fast differentiation driven by high environmental dynamism"

| Gene | Group | NS_mut | ENSEMBL | cds_L | NS_norm_cds_L |
|---|---|---|---|---|---|
| ACAD10 | Common_BALD | 1 | ENSP00000389813 | 3245 | 3,08E-04 |
| ACAD10 | Common_BLACK | 1 | ENSP00000389813 | 3245 | 3,08E-04 |
| ADAM29 | Common_BALD | 1 | ENSP00000384229 | 2399 | 4,17E-04 |
| ADAM29 | Common_BLACK | 1 | ENSP00000384229 | 2399 | 4,17E-04 |
| ADAMTSL3 | Common_BALD | 1 | ENSP00000286744 | 5105 | 1,96E-04 |
| ADAMTSL3 | Common_BLACK | 2 | ENSP00000286744 | 5105 | 3,92E-04 |
| ADGB | Common_BALD | 1 | ENSP00000381036 | 4953 | 2,02E-04 |
| ADGB | Common_BLACK | 2 | ENSP00000381036 | 4953 | 4,04E-04 |
| AKAP11 | Common_BALD | 1 | ENSP00000025301 | 5639 | 1,77E-04 |
| AKAP11 | Common_BLACK | 1 | ENSP00000025301 | 5639 | 1,77E-04 |
| AKAP6 | Common_BALD | 2 | ENSP00000280979 | 6634 | 3,01E-04 |
| AKAP6 | Common_BLACK | 1 | ENSP00000280979 | 6634 | 1,51E-04 |
| ALMS1 | Common_BALD | 1 | ENSP00000482968 | 11916 | 8,39E-05 |
| ALMS1 | Common_BLACK | 1 | ENSP00000482968 | 11916 | 8,39E-05 |
| ALPK1 | Common_BALD | 3 | ENSP00000177648 | 3725 | 8,05E-04 |
| ALPK1 | Common_BLACK | 2 | ENSP00000177648 | 3725 | 5,37E-04 |
| ALPK2 | Common_BALD | 1 | ENSP00000354991 | 6490 | 1,54E-04 |
| ALPK2 | Common_BLACK | 1 | ENSP00000354991 | 6490 | 1,54E-04 |
| ANKRD60 | Common_BALD | 1 | ENSP00000396747 | 1009 | 9,91E-04 |
| ANKRD60 | Common_BLACK | 2 | ENSP00000396747 | 1009 | 1,98E-03 |
| APOB | Common_BALD | 3 | ENSP00000233242 | 13654 | 2,20E-04 |
| APOB | Common_BLACK | 5 | ENSP00000233242 | 13654 | 3,66E-04 |
| ARHGEF5 | Common_BALD | 1 | ENSP00000056217 | 4763 | 2,10E-04 |
| ARHGEF5 | Common_BLACK | 1 | ENSP00000056217 | 4763 | 2,10E-04 |
| ARSK | Common_BALD | 1 | ENSP00000369346 | 1603 | 6,24E-04 |
| ARSK | Common_BLACK | 1 | ENSP00000369346 | 1603 | 6,24E-04 |
| ATM | Common_BALD | 1 | ENSP00000388058 | 9106 | 1,10E-04 |
| ATM | Common_BLACK | 1 | ENSP00000388058 | 9106 | 1,10E-04 |
| BRCA2 | Common_BALD | 5 | ENSP00000369497 | 10283 | 4,86E-04 |
| BRCA2 | Common_BLACK | 2 | ENSP00000369497 | 10283 | 1,94E-04 |
| BTBD8 | Common_BALD | 2 | ENSP00000490161 | 5354 | 3,74E-04 |
| BTBD8 | Common_BLACK | 2 | ENSP00000490161 | 5354 | 3,74E-04 |
| C1orf127 | Common_BALD | 5 | ENSP00000366203 | 2490 | 2,01E-03 |
| C1orf127 | Common_BLACK | 1 | ENSP00000366203 | 2490 | 4,02E-04 |
| C4orf19 | Common_BALD | 2 | ENSP00000371408 | 943 | 2,12E-03 |
| C4orf19 | Common_BLACK | 2 | ENSP00000371408 | 943 | 2,12E-03 |
| C8orf74 | Common_BALD | 1 | ENSP00000307129 | 891 | 1,12E-03 |
| C8orf74 | Common_BLACK | 1 | ENSP00000307129 | 891 | 1,12E-03 |
| CASP8AP2 | Common_BALD | 2 | ENSP00000478179 | 6050 | 3,31E-04 |
| CASP8AP2 | Common_BLACK | 1 | ENSP00000478179 | 6050 | 1,65E-04 |
| CASS4 | Common_BALD | 2 | ENSP00000353462 | 2314 | 8,64E-04 |
| CASS4 | Common_BLACK | 3 | ENSP00000353462 | 2314 | 1,30E-03 |
| CCDC39 | Common_BALD | 1 | ENSP00000417960 | 2809 | 3,56E-04 |
| CCDC39 | Common_BLACK | 1 | ENSP00000417960 | 2809 | 3,56E-04 |
| CCDC66 | Common_BALD | 1 | ENSP00000378167 | 1783 | 5,61E-04 |
| CCDC66 | Common_BLACK | 1 | ENSP00000378167 | 1783 | 5,61E-04 |
| CCDC73 | Common_BALD | 2 | ENSP00000335325 | 3198 | 6,25E-04 |

| | | | | | |
|---|---|---|---|---|---|
| CCDC73 | Common_BLACK | 1 | ENSP00000335325 | 3198 | 3,13E-04 |
| CD19 | Common_BALD | 1 | ENSP00000313419 | 1740 | 5,75E-04 |
| CD19 | Common_BLACK | 1 | ENSP00000313419 | 1740 | 5,75E-04 |
| CDK5RAP2 | Common_BALD | 2 | ENSP00000343818 | 5971 | 3,35E-04 |
| CDK5RAP2 | Common_BLACK | 1 | ENSP00000343818 | 5971 | 1,67E-04 |
| CEACAM20 | Common_BALD | 1 | ENSP00000481937 | 1745 | 5,73E-04 |
| CEACAM20 | Common_BLACK | 1 | ENSP00000481937 | 1745 | 5,73E-04 |
| CEACAM5 | Common_BALD | 1 | ENSP00000221992 | 3825 | 2,61E-04 |
| CEACAM5 | Common_BLACK | 3 | ENSP00000221992 | 3825 | 7,84E-04 |
| CENPJ | Common_BALD | 2 | ENSP00000371308 | 4007 | 4,99E-04 |
| CENPJ | Common_BLACK | 1 | ENSP00000371308 | 4007 | 2,50E-04 |
| CEP295 | Common_BALD | 1 | ENSP00000316681 | 7764 | 1,29E-04 |
| CEP295 | Common_BLACK | 3 | ENSP00000316681 | 7764 | 3,86E-04 |
| CFH | Common_BALD | 1 | ENSP00000356399 | 3615 | 2,77E-04 |
| CFH | Common_BLACK | 1 | ENSP00000356399 | 3615 | 2,77E-04 |
| CLMN | Common_BALD | 2 | ENSP00000298912 | 3005 | 6,66E-04 |
| CLMN | Common_BLACK | 1 | ENSP00000298912 | 3005 | 3,33E-04 |
| CLUL1 | Common_BALD | 1 | ENSP00000441726 | 1397 | 7,16E-04 |
| CLUL1 | Common_BLACK | 1 | ENSP00000441726 | 1397 | 7,16E-04 |
| CNTLN | Common_BALD | 2 | ENSP00000370021 | 4200 | 4,76E-04 |
| CNTLN | Common_BLACK | 1 | ENSP00000370021 | 4200 | 2,38E-04 |
| COBLL1 | Common_BALD | 1 | ENSP00000487041 | 3609 | 2,77E-04 |
| COBLL1 | Common_BLACK | 3 | ENSP00000487041 | 3609 | 8,31E-04 |
| CPXM1 | Common_BALD | 1 | ENSP00000369979 | 2177 | 4,59E-04 |
| CPXM1 | Common_BLACK | 1 | ENSP00000369979 | 2177 | 4,59E-04 |
| CR2 | Common_BALD | 1 | ENSP00000356024 | 3302 | 3,03E-04 |
| CR2 | Common_BLACK | 1 | ENSP00000356024 | 3302 | 3,03E-04 |
| CRNN | Common_BALD | 1 | ENSP00000271835 | 1641 | 6,09E-04 |
| CRNN | Common_BLACK | 1 | ENSP00000271835 | 1641 | 6,09E-04 |
| CTSG | Common_BALD | 2 | ENSP00000216336 | 763 | 2,62E-03 |
| CTSG | Common_BLACK | 2 | ENSP00000216336 | 763 | 2,62E-03 |
| CTSS | Common_BALD | 1 | ENSP00000357981 | 986 | 1,01E-03 |
| CTSS | Common_BLACK | 1 | ENSP00000357981 | 986 | 1,01E-03 |
| CUBN | Common_BALD | 2 | ENSP00000367064 | 10802 | 1,85E-04 |
| CUBN | Common_BLACK | 1 | ENSP00000367064 | 10802 | 9,26E-05 |
| CYP3A4 | Common_BALD | 2 | ENSP00000337915 | 1499 | 1,33E-03 |
| CYP3A4 | Common_BLACK | 1 | ENSP00000337915 | 1499 | 6,67E-04 |
| CYP4A11 | Common_BALD | 1 | ENSP00000360971 | 2930 | 3,41E-04 |
| CYP4A11 | Common_BLACK | 3 | ENSP00000360971 | 2930 | 1,02E-03 |
| DDX60 | Common_BALD | 1 | ENSP00000377344 | 5029 | 1,99E-04 |
| DDX60 | Common_BLACK | 2 | ENSP00000377344 | 5029 | 3,98E-04 |
| DNAH9 | Common_BALD | 1 | ENSP00000262442 | 13046 | 7,67E-05 |
| DNAH9 | Common_BLACK | 1 | ENSP00000262442 | 13046 | 7,67E-05 |
| EDDM3B | Common_BALD | 1 | ENSP00000314810 | 443 | 2,26E-03 |
| EDDM3B | Common_BLACK | 1 | ENSP00000314810 | 443 | 2,26E-03 |
| EFCAB6 | Common_BALD | 1 | ENSP00000262726 | 4475 | 2,23E-04 |
| EFCAB6 | Common_BLACK | 2 | ENSP00000262726 | 4475 | 4,47E-04 |
| EIF2B2 | Common_BALD | 1 | ENSP00000266126 | 1048 | 9,54E-04 |

| | | | | | |
|---|---|---|---|---|---|
| EIF2B2 | Common_BLACK | 1 | ENSP00000266126 | 1048 | 9,54E-04 |
| EPG5 | Common_BALD | 2 | ENSP00000282041 | 7631 | 2,62E-04 |
| EPG5 | Common_BLACK | 1 | ENSP00000282041 | 7631 | 1,31E-04 |
| EVC2 | Common_BALD | 1 | ENSP00000342144 | 3882 | 2,58E-04 |
| EVC2 | Common_BLACK | 1 | ENSP00000342144 | 3882 | 2,58E-04 |
| F5 | Common_BALD | 1 | ENSP00000356770 | 6567 | 1,52E-04 |
| F5 | Common_BLACK | 1 | ENSP00000356770 | 6567 | 1,52E-04 |
| FAM160A1 | Common_BALD | 1 | ENSP00000413196 | 3115 | 3,21E-04 |
| FAM160A1 | Common_BLACK | 1 | ENSP00000413196 | 3115 | 3,21E-04 |
| FANCA | Common_BALD | 1 | ENSP00000373952 | 4469 | 2,24E-04 |
| FANCA | Common_BLACK | 1 | ENSP00000373952 | 4469 | 2,24E-04 |
| FHAD1 | Common_BALD | 1 | ENSP00000365166 | 4195 | 2,38E-04 |
| FHAD1 | Common_BLACK | 2 | ENSP00000365166 | 4195 | 4,77E-04 |
| FKBP15 | Common_BALD | 1 | ENSP00000238256 | 3710 | 2,70E-04 |
| FKBP15 | Common_BLACK | 2 | ENSP00000238256 | 3710 | 5,39E-04 |
| FOLH1 | Common_BALD | 1 | ENSP00000256999 | 2234 | 4,48E-04 |
| FOLH1 | Common_BLACK | 1 | ENSP00000256999 | 2234 | 4,48E-04 |
| FREM1 | Common_BALD | 2 | ENSP00000370262 | 6554 | 3,05E-04 |
| FREM1 | Common_BLACK | 2 | ENSP00000370262 | 6554 | 3,05E-04 |
| GGT6 | Common_BALD | 2 | ENSP00000370962 | 1464 | 1,37E-03 |
| GGT6 | Common_BLACK | 1 | ENSP00000370962 | 1464 | 6,83E-04 |
| GLIPR1 | Common_BALD | 1 | ENSP00000266659 | 802 | 1,25E-03 |
| GLIPR1 | Common_BLACK | 1 | ENSP00000266659 | 802 | 1,25E-03 |
| GLIPR1L1 | Common_BALD | 1 | ENSP00000367967 | 697 | 1,43E-03 |
| GLIPR1L1 | Common_BLACK | 1 | ENSP00000367967 | 697 | 1,43E-03 |
| GZMK | Common_BALD | 1 | ENSP00000231009 | 790 | 1,27E-03 |
| GZMK | Common_BLACK | 1 | ENSP00000231009 | 790 | 1,27E-03 |
| HELB | Common_BALD | 1 | ENSP00000247815 | 3248 | 3,08E-04 |
| HELB | Common_BLACK | 1 | ENSP00000247815 | 3248 | 3,08E-04 |
| HIVEP1 | Common_BALD | 1 | ENSP00000368698 | 8126 | 1,23E-04 |
| HIVEP1 | Common_BLACK | 5 | ENSP00000368698 | 8126 | 6,15E-04 |
| HMOX1 | Common_BALD | 1 | ENSP00000216117 | 839 | 1,19E-03 |
| HMOX1 | Common_BLACK | 1 | ENSP00000216117 | 839 | 1,19E-03 |
| IFNK | Common_BALD | 2 | ENSP00000276943 | 623 | 3,21E-03 |
| IFNK | Common_BLACK | 1 | ENSP00000276943 | 623 | 1,61E-03 |
| IGSF10 | Common_BALD | 3 | ENSP00000282466 | 7866 | 3,81E-04 |
| IGSF10 | Common_BLACK | 1 | ENSP00000282466 | 7866 | 1,27E-04 |
| IQCH | Common_BALD | 1 | ENSP00000336861 | 3059 | 3,27E-04 |
| IQCH | Common_BLACK | 1 | ENSP00000336861 | 3059 | 3,27E-04 |
| ITIH4 | Common_BALD | 1 | ENSP00000417824 | 2782 | 3,59E-04 |
| ITIH4 | Common_BLACK | 1 | ENSP00000417824 | 2782 | 3,59E-04 |
| KIF20B | Common_BALD | 2 | ENSP00000360793 | 5431 | 3,68E-04 |
| KIF20B | Common_BLACK | 1 | ENSP00000360793 | 5431 | 1,84E-04 |
| LACC1 | Common_BALD | 1 | ENSP00000317619 | 1288 | 7,76E-04 |
| LACC1 | Common_BLACK | 1 | ENSP00000317619 | 1288 | 7,76E-04 |
| LCOR | Common_BALD | 1 | ENSP00000490116 | 4656 | 2,15E-04 |
| LCOR | Common_BLACK | 1 | ENSP00000490116 | 4656 | 2,15E-04 |
| LILRA4 | Common_BALD | 1 | ENSP00000291759 | 1415 | 7,07E-04 |

| | | | | | |
|---|---|---|---|---|---|
| LILRA4 | Common_BLACK | 1 | ENSP00000291759 | 1415 | 7,07E-04 |
| LNPEP | Common_BALD | 1 | ENSP00000231368 | 3040 | 3,29E-04 |
| LNPEP | Common_BLACK | 1 | ENSP00000231368 | 3040 | 3,29E-04 |
| LRP2 | Common_BALD | 1 | ENSP00000496870 | 13808 | 7,24E-05 |
| LRP2 | Common_BLACK | 2 | ENSP00000496870 | 13808 | 1,45E-04 |
| LRRC42 | Common_BALD | 1 | ENSP00000360421 | 1280 | 7,81E-04 |
| LRRC42 | Common_BLACK | 1 | ENSP00000360421 | 1280 | 7,81E-04 |
| LRRC53 | Common_BALD | 3 | ENSP00000294635 | 3719 | 8,07E-04 |
| LRRC53 | Common_BLACK | 1 | ENSP00000294635 | 3719 | 2,69E-04 |
| LYST | Common_BALD | 1 | ENSP00000374443 | 11357 | 8,81E-05 |
| LYST | Common_BLACK | 1 | ENSP00000374443 | 11357 | 8,81E-05 |
| MAP4 | Common_BALD | 2 | ENSP00000407602 | 6775 | 2,95E-04 |
| MAP4 | Common_BLACK | 2 | ENSP00000407602 | 6775 | 2,95E-04 |
| MDN1 | Common_BALD | 3 | ENSP00000358400 | 16677 | 1,80E-04 |
| MDN1 | Common_BLACK | 1 | ENSP00000358400 | 16677 | 6,00E-05 |
| MIA3 | Common_BALD | 1 | ENSP00000340900 | 5715 | 1,75E-04 |
| MIA3 | Common_BLACK | 1 | ENSP00000340900 | 5715 | 1,75E-04 |
| MMP3 | Common_BALD | 1 | ENSP00000299855 | 1421 | 7,04E-04 |
| MMP3 | Common_BLACK | 1 | ENSP00000299855 | 1421 | 7,04E-04 |
| MNDA | Common_BALD | 4 | ENSP00000357123 | 1183 | 3,38E-03 |
| MNDA | Common_BLACK | 3 | ENSP00000357123 | 1183 | 2,54E-03 |
| MOV10L1 | Common_BALD | 1 | ENSP00000262794 | 3606 | 2,77E-04 |
| MOV10L1 | Common_BLACK | 2 | ENSP00000262794 | 3606 | 5,55E-04 |
| MX1 | Common_BALD | 1 | ENSP00000381601 | 1976 | 5,06E-04 |
| MX1 | Common_BLACK | 1 | ENSP00000381601 | 1976 | 5,06E-04 |
| MYH15 | Common_BALD | 1 | ENSP00000273353 | 5759 | 1,74E-04 |
| MYH15 | Common_BLACK | 1 | ENSP00000273353 | 5759 | 1,74E-04 |
| MYRFL | Common_BALD | 1 | ENSP00000448753 | 2658 | 3,76E-04 |
| MYRFL | Common_BLACK | 1 | ENSP00000448753 | 2658 | 3,76E-04 |
| N4BP2 | Common_BALD | 1 | ENSP00000261435 | 5256 | 1,90E-04 |
| N4BP2 | Common_BLACK | 1 | ENSP00000261435 | 5256 | 1,90E-04 |
| NCOA3 | Common_BALD | 1 | ENSP00000361066 | 4246 | 2,36E-04 |
| NCOA3 | Common_BLACK | 1 | ENSP00000361066 | 4246 | 2,36E-04 |
| NCOA4 | Common_BALD | 1 | ENSP00000463027 | 1946 | 5,14E-04 |
| NCOA4 | Common_BLACK | 1 | ENSP00000463027 | 1946 | 5,14E-04 |
| NDUFAF1 | Common_BALD | 1 | ENSP00000260361 | 980 | 1,02E-03 |
| NDUFAF1 | Common_BLACK | 1 | ENSP00000260361 | 980 | 1,02E-03 |
| NECTIN2 | Common_BALD | 1 | ENSP00000252483 | 1506 | 6,64E-04 |
| NECTIN2 | Common_BLACK | 1 | ENSP00000252483 | 1506 | 6,64E-04 |
| NFKB1 | Common_BALD | 2 | ENSP00000226574 | 2849 | 7,02E-04 |
| NFKB1 | Common_BLACK | 1 | ENSP00000226574 | 2849 | 3,51E-04 |
| NHSL1 | Common_BALD | 1 | ENSP00000394546 | 4822 | 2,07E-04 |
| NHSL1 | Common_BLACK | 1 | ENSP00000394546 | 4822 | 2,07E-04 |
| NKTR | Common_BALD | 1 | ENSP00000232978 | 4411 | 2,27E-04 |
| NKTR | Common_BLACK | 1 | ENSP00000232978 | 4411 | 2,27E-04 |
| NLRP10 | Common_BALD | 1 | ENSP00000327763 | 1951 | 5,13E-04 |
| NLRP10 | Common_BLACK | 2 | ENSP00000327763 | 1951 | 1,03E-03 |
| NLRP4 | Common_BALD | 2 | ENSP00000301295 | 2948 | 6,78E-04 |

| | | | | | |
|---|---|---|---|---|---|
| NLRP4 | Common_BLACK | 2 | ENSP00000301295 | 2948 | 6,78E-04 |
| NRAP | Common_BALD | 1 | ENSP00000358365 | 5157 | 1,94E-04 |
| NRAP | Common_BLACK | 1 | ENSP00000358365 | 5157 | 1,94E-04 |
| NUTM1 | Common_BALD | 1 | ENSP00000444896 | 3482 | 2,87E-04 |
| NUTM1 | Common_BLACK | 1 | ENSP00000444896 | 3482 | 2,87E-04 |
| AADACL4 | BALD | 1 | ENSP00000365395 | 1220 | 8,20E-04 |
| AASS | BALD | 1 | ENSP00000377040 | 2758 | 3,63E-04 |
| ABHD10 | BALD | 1 | ENSP00000273359 | 912 | 1,10E-03 |
| ACAD8 | BALD | 1 | ENSP00000281182 | 1237 | 8,08E-04 |
| ACAD9 | BALD | 1 | ENSP00000312618 | 1887 | 5,30E-04 |
| ACKR2 | BALD | 1 | ENSP00000396150 | 836 | 1,20E-03 |
| ACOT7 | BALD | 1 | ENSP00000367086 | 1133 | 8,83E-04 |
| ACOX1 | BALD | 1 | ENSP00000293217 | 1972 | 5,07E-04 |
| ACSL6 | BALD | 1 | ENSP00000498260 | 2104 | 4,75E-04 |
| ADAMTS10 | BALD | 1 | ENSP00000471851 | 3185 | 3,14E-04 |
| ADAMTS12 | BALD | 1 | ENSP00000422554 | 4651 | 2,15E-04 |
| ADAT1 | BALD | 1 | ENSP00000310015 | 1500 | 6,67E-04 |
| ADPRM | BALD | 1 | ENSP00000369099 | 1035 | 9,66E-04 |
| AGBL2 | BALD | 1 | ENSP00000350228 | 2570 | 3,89E-04 |
| AHI1 | BALD | 1 | ENSP00000322478 | 3553 | 2,81E-04 |
| AKAP13 | BALD | 1 | ENSP00000354718 | 8411 | 1,19E-04 |
| AKNA | BALD | 1 | ENSP00000363201 | 4011 | 2,49E-04 |
| AKR7A3 | BALD | 1 | ENSP00000355377 | 1957 | 5,11E-04 |
| ALDH7A1 | BALD | 1 | ENSP00000490811 | 1552 | 6,44E-04 |
| ALG8 | BALD | 1 | ENSP00000299626 | 1565 | 6,39E-04 |
| ALKBH7 | BALD | 1 | ENSP00000245812 | 662 | 1,51E-03 |
| ANKRD2 | BALD | 1 | ENSP00000306163 | 1067 | 9,37E-04 |
| APOA4 | BALD | 1 | ENSP00000350425 | 1183 | 8,45E-04 |
| APPL2 | BALD | 1 | ENSP00000446917 | 1939 | 5,16E-04 |
| AQR | BALD | 2 | ENSP00000156471 | 4439 | 4,51E-04 |
| ARHGAP21 | BALD | 1 | ENSP00000379709 | 5758 | 1,74E-04 |
| ARHGAP40 | BALD | 2 | ENSP00000362442 | 1852 | 1,08E-03 |
| ARHGEF11 | BALD | 1 | ENSP00000357177 | 4638 | 2,16E-04 |
| ARHGEF37 | BALD | 1 | ENSP00000328083 | 2008 | 4,98E-04 |
| ARID3C | BALD | 1 | ENSP00000368189 | 1229 | 8,14E-04 |
| ASIC5 | BALD | 1 | ENSP00000442477 | 1464 | 6,83E-04 |
| ASPSCR1 | BALD | 1 | ENSP00000306625 | 1593 | 6,28E-04 |
| ASXL3 | BALD | 2 | ENSP00000269197 | 6735 | 2,97E-04 |
| ATE1 | BALD | 1 | ENSP00000358039 | 1548 | 6,46E-04 |
| ATF7IP2 | BALD | 2 | ENSP00000379808 | 2024 | 9,88E-04 |
| ATG12 | BALD | 1 | ENSP00000425107 | 381 | 2,62E-03 |
| ATP10D | BALD | 1 | ENSP00000273859 | 4250 | 2,35E-04 |
| ATP11B | BALD | 1 | ENSP00000321195 | 3502 | 2,86E-04 |
| ATP12A | BALD | 1 | ENSP00000218548 | 3122 | 3,20E-04 |
| ATP4B | BALD | 1 | ENSP00000334216 | 869 | 1,15E-03 |
| ATP6V0A1 | BALD | 1 | ENSP00000264649 | 2497 | 4,00E-04 |
| B3GALT5 | BALD | 1 | ENSP00000381699 | 935 | 1,07E-03 |
| BAALC | BALD | 1 | ENSP00000297574 | 539 | 1,86E-03 |

| | | | | | |
|---|---|---|---|---|---|
| BACH1 | BALD | 1 | ENSP00000286800 | 2207 | 4,53E-04 |
| BAG3 | BALD | 1 | ENSP00000358081 | 1717 | 5,82E-04 |
| BAIAP2L1 | BALD | 1 | ENSP00000005260 | 1363 | 7,34E-04 |
| BBS10 | BALD | 1 | ENSP00000497413 | 2170 | 4,61E-04 |
| BDNF | BALD | 1 | ENSP00000414303 | 991 | 1,01E-03 |
| BECN2 | BALD | 1 | ENSP00000488361 | 1295 | 7,72E-04 |
| BPIFB6 | BALD | 1 | ENSP00000344929 | 1884 | 5,31E-04 |
| BRCA1 | BALD | 2 | ENSP00000418960 | 5472 | 3,65E-04 |
| BRD8 | BALD | 1 | ENSP00000254900 | 3679 | 2,72E-04 |
| BTBD16 | BALD | 1 | ENSP00000260723 | 1531 | 6,53E-04 |
| BTNL8 | BALD | 1 | ENSP00000342197 | 1427 | 7,01E-04 |
| C10orf67 | BALD | 1 | ENSP00000490528 | 1493 | 6,70E-04 |
| C11orf24 | BALD | 1 | ENSP00000307264 | 1241 | 8,06E-04 |
| C11orf71 | BALD | 1 | ENSP00000325508 | 437 | 2,29E-03 |
| C11orf80 | BALD | 1 | ENSP00000354227 | 1508 | 6,63E-04 |
| C14orf39 | BALD | 1 | ENSP00000324920 | 1747 | 5,72E-04 |
| C17orf80 | BALD | 2 | ENSP00000440551 | 1832 | 1,09E-03 |
| C1orf105 | BALD | 2 | ENSP00000356700 | 524 | 3,82E-03 |
| C1QC | BALD | 1 | ENSP00000363771 | 723 | 1,38E-03 |
| C1QTNF1 | BALD | 1 | ENSP00000462990 | 808 | 1,24E-03 |
| C2CD3 | BALD | 1 | ENSP00000334379 | 7019 | 1,42E-04 |
| C2orf16 | BALD | 1 | ENSP00000403181 | 5279 | 1,89E-04 |
| C2orf69 | BALD | 1 | ENSP00000312770 | 1167 | 8,57E-04 |
| C4orf51 | BALD | 1 | ENSP00000391404 | 644 | 1,55E-03 |
| C5orf47 | BALD | 1 | ENSP00000340887 | 521 | 1,92E-03 |
| C6orf89 | BALD | 1 | ENSP00000347322 | 1055 | 9,48E-04 |
| C9orf43 | BALD | 1 | ENSP00000363280 | 1392 | 7,18E-04 |
| C9orf64 | BALD | 1 | ENSP00000365522 | 1018 | 9,82E-04 |
| CATSPER1 | BALD | 3 | ENSP00000309052 | 2280 | 1,32E-03 |
| CATSPERG | BALD | 1 | ENSP00000386962 | 3521 | 2,84E-04 |
| CC2D1B | BALD | 1 | ENSP00000360642 | 2538 | 3,94E-04 |
| CC2D2A | BALD | 1 | ENSP00000421809 | 4813 | 2,08E-04 |
| CCDC110 | BALD | 1 | ENSP00000427246 | 2458 | 4,07E-04 |
| CCDC121 | BALD | 1 | ENSP00000412150 | 1306 | 7,66E-04 |
| CCDC146 | BALD | 1 | ENSP00000285871 | 2850 | 3,51E-04 |
| CCDC170 | BALD | 1 | ENSP00000239374 | 2093 | 4,78E-04 |
| CCDC34 | BALD | 1 | ENSP00000330240 | 1089 | 9,18E-04 |
| CCDC68 | BALD | 1 | ENSP00000466690 | 945 | 1,06E-03 |
| CCDC71 | BALD | 1 | ENSP00000319006 | 1370 | 7,30E-04 |
| CCER2 | BALD | 1 | ENSP00000460665 | 753 | 1,33E-03 |
| CCPG1 | BALD | 1 | ENSP00000403400 | 2404 | 4,16E-04 |
| CD109 | BALD | 1 | ENSP00000287097 | 4244 | 2,36E-04 |
| CD1B | BALD | 1 | ENSP00000357150 | 1784 | 5,61E-04 |
| CD36 | BALD | 1 | ENSP00000399421 | 1407 | 7,11E-04 |
| CD37 | BALD | 1 | ENSP00000325708 | 830 | 1,20E-03 |
| CD44 | BALD | 1 | ENSP00000398632 | 2146 | 4,66E-04 |
| CD48 | BALD | 1 | ENSP00000484431 | 624 | 1,60E-03 |
| CD59 | BALD | 1 | ENSP00000498879 | 389 | 2,57E-03 |

| | | | | | |
|---|---|---|---|---|---|
| CDH19 | BALD | 1 | ENSP00000262150 | 2311 | 4,33E-04 |
| CENPT | BALD | 1 | ENSP00000400140 | 1691 | 5,91E-04 |
| CEP57L1 | BALD | 1 | ENSP00000357966 | 1486 | 6,73E-04 |
| CEP83 | BALD | 2 | ENSP00000380911 | 2091 | 9,56E-04 |
| CEP85 | BALD | 1 | ENSP00000252992 | 2276 | 4,39E-04 |
| CHCHD6 | BALD | 1 | ENSP00000422912 | 1037 | 9,64E-04 |
| CHD9 | BALD | 1 | ENSP00000381522 | 8658 | 1,16E-04 |
| CLDN17 | BALD | 1 | ENSP00000286808 | 674 | 1,48E-03 |
| CLEC16A | BALD | 1 | ENSP00000387122 | 3153 | 3,17E-04 |
| CMTM5 | BALD | 1 | ENSP00000344819 | 653 | 1,53E-03 |
| CNGA3 | BALD | 1 | ENSP00000386761 | 2078 | 4,81E-04 |
| CNST | BALD | 1 | ENSP00000355470 | 2129 | 4,70E-04 |
| COL11A1 | BALD | 1 | ENSP00000351163 | 5387 | 1,86E-04 |
| COL19A1 | BALD | 1 | ENSP00000480474 | 3370 | 2,97E-04 |
| COL6A5 | BALD | 2 | ENSP00000309762 | 7776 | 2,57E-04 |
| COMMD9 | BALD | 1 | ENSP00000263401 | 591 | 1,69E-03 |
| CPN1 | BALD | 1 | ENSP00000359446 | 1371 | 7,29E-04 |
| CPOX | BALD | 1 | ENSP00000497326 | 1344 | 7,44E-04 |
| CRP | BALD | 1 | ENSP00000255030 | 670 | 1,49E-03 |
| CSF1R | BALD | 2 | ENSP00000286301 | 2853 | 7,01E-04 |
| CSF2RB | BALD | 1 | ENSP00000385271 | 2656 | 3,77E-04 |
| CSF3 | BALD | 1 | ENSP00000225474 | 592 | 1,69E-03 |
| CSGALNACT | BALD | 1 | ENSP00000411816 | 1592 | 6,28E-04 |
| CSMD1 | BALD | 1 | ENSP00000430733 | 10531 | 9,50E-05 |
| CST9 | BALD | 1 | ENSP00000366170 | 483 | 2,07E-03 |
| CTTN | BALD | 1 | ENSP00000365745 | 1641 | 6,09E-04 |
| CXCL8 | BALD | 1 | ENSP00000306512 | 285 | 3,51E-03 |
| CYLD | BALD | 1 | ENSP00000308928 | 2846 | 3,51E-04 |
| CYP2S1 | BALD | 1 | ENSP00000308032 | 1489 | 6,72E-04 |
| CYP4B1 | BALD | 1 | ENSP00000360991 | 1557 | 6,42E-04 |
| DALRD3 | BALD | 1 | ENSP00000344989 | 1612 | 6,20E-04 |
| DAP3 | BALD | 1 | ENSP00000357320 | 1182 | 8,46E-04 |
| DBT | BALD | 1 | ENSP00000359151 | 1438 | 6,95E-04 |
| DCAF4 | BALD | 1 | ENSP00000351147 | 1484 | 6,74E-04 |
| DCT | BALD | 1 | ENSP00000392762 | 1688 | 5,92E-04 |
| DDX60L | BALD | 2 | ENSP00000422423 | 5031 | 3,98E-04 |
| DFFA | BALD | 1 | ENSP00000366237 | 990 | 1,01E-03 |
| DHX38 | BALD | 1 | ENSP00000268482 | 3651 | 2,74E-04 |
| DIP2A | BALD | 1 | ENSP00000498874 | 4614 | 2,17E-04 |
| DNAAF1 | BALD | 1 | ENSP00000367815 | 1975 | 5,06E-04 |
| DNAI2 | BALD | 1 | ENSP00000464197 | 1969 | 5,08E-04 |
| DNAJB6 | BALD | 1 | ENSP00000488263 | 993 | 1,01E-03 |
| DNAJB9 | BALD | 1 | ENSP00000249356 | 667 | 1,50E-03 |
| DNAJC1 | BALD | 1 | ENSP00000366179 | 1648 | 6,07E-04 |
| DNAJC4 | BALD | 1 | ENSP00000320548 | 734 | 1,36E-03 |
| DTNA | BALD | 1 | ENSP00000382064 | 2157 | 4,64E-04 |
| DTX3L | BALD | 2 | ENSP00000296161 | 2218 | 9,02E-04 |
| DUSP13 | BALD | 1 | ENSP00000475626 | 947 | 1,06E-03 |

| | | | | | |
|---|---|---|---|---|---|
| DYNC2H1 | BALD | 1 | ENSP00000497174 | 12834 | 7,79E-05 |
| ECM1 | BALD | 1 | ENSP00000358045 | 1696 | 5,90E-04 |
| ECSIT | BALD | 1 | ENSP00000270517 | 1276 | 7,84E-04 |
| EDRF1 | BALD | 1 | ENSP00000349244 | 3695 | 2,71E-04 |
| EFCAB8 | BALD | 1 | ENSP00000383366 | 3823 | 2,62E-04 |
| EGFLAM | BALD | 1 | ENSP00000346964 | 2922 | 3,42E-04 |
| EIF2B3 | BALD | 1 | ENSP00000353575 | 1304 | 7,67E-04 |
| EIF4G1 | BALD | 1 | ENSP00000371767 | 4764 | 2,10E-04 |
| EIF4G3 | BALD | 1 | ENSP00000383274 | 4996 | 2,00E-04 |
| ELF1 | BALD | 1 | ENSP00000239882 | 1998 | 5,01E-04 |
| EME1 | BALD | 1 | ENSP00000421700 | 1708 | 5,85E-04 |
| ENTPD3 | BALD | 1 | ENSP00000301825 | 1414 | 7,07E-04 |
| EPSTI1 | BALD | 1 | ENSP00000318982 | 1229 | 8,14E-04 |
| ERCC6 | BALD | 1 | ENSP00000348089 | 4462 | 2,24E-04 |
| ERICH5 | BALD | 1 | ENSP00000315614 | 1304 | 7,67E-04 |
| EXOSC2 | BALD | 1 | ENSP00000361433 | 873 | 1,15E-03 |
| FADD | BALD | 1 | ENSP00000301838 | 618 | 1,62E-03 |
| FAF1 | BALD | 1 | ENSP00000379457 | 1905 | 5,25E-04 |
| FAM124B | BALD | 1 | ENSP00000386895 | 1366 | 7,32E-04 |
| FAM151A | BALD | 2 | ENSP00000306888 | 1746 | 1,15E-03 |
| FAM160B1 | BALD | 1 | ENSP00000358251 | 2281 | 4,38E-04 |
| FAM189A1 | BALD | 1 | ENSP00000261275 | 1560 | 6,41E-04 |
| FAM192A | BALD | 1 | ENSP00000335808 | 756 | 1,32E-03 |
| FAM219B | BALD | 1 | ENSP00000350260 | 591 | 1,69E-03 |
| FAM71C | BALD | 1 | ENSP00000315247 | 721 | 1,39E-03 |
| FAM83B | BALD | 1 | ENSP00000304078 | 3032 | 3,30E-04 |
| FASTKD5 | BALD | 1 | ENSP00000369618 | 2294 | 4,36E-04 |
| FBXW10 | BALD | 1 | ENSP00000310382 | 3176 | 3,15E-04 |
| FEZ2 | BALD | 1 | ENSP00000368547 | 1168 | 8,56E-04 |
| FGD5 | BALD | 3 | ENSP00000285046 | 4315 | 6,95E-04 |
| FGFBP2 | BALD | 1 | ENSP00000259989 | 671 | 1,49E-03 |
| FGL1 | BALD | 1 | ENSP00000381133 | 927 | 1,08E-03 |
| FIP1L1 | BALD | 1 | ENSP00000336752 | 1557 | 6,42E-04 |
| FITM2 | BALD | 1 | ENSP00000380037 | 783 | 1,28E-03 |
| FLT1 | BALD | 1 | ENSP00000282397 | 3922 | 2,55E-04 |
| FNDC3B | BALD | 1 | ENSP00000389094 | 3586 | 2,79E-04 |
| FOXR1 | BALD | 1 | ENSP00000314806 | 811 | 1,23E-03 |
| FPGT-TNNI3 | BALD | 1 | ENSP00000450895 | 2823 | 3,54E-04 |
| FRRS1 | BALD | 1 | ENSP00000287474 | 1863 | 5,37E-04 |
| FTMT | BALD | 2 | ENSP00000313691 | 728 | 2,75E-03 |
| GALNT11 | BALD | 1 | ENSP00000395122 | 1816 | 5,51E-04 |
| GARNL3 | BALD | 1 | ENSP00000362485 | 3047 | 3,28E-04 |
| GART | BALD | 1 | ENSP00000371236 | 3012 | 3,32E-04 |
| GBA2 | BALD | 1 | ENSP00000367343 | 2763 | 3,62E-04 |
| GEMIN7 | BALD | 1 | ENSP00000466342 | 550 | 1,82E-03 |
| GEN1 | BALD | 1 | ENSP00000318977 | 2708 | 3,69E-04 |
| GLCE | BALD | 1 | ENSP00000261858 | 1854 | 5,39E-04 |
| GLIPR1L2 | BALD | 1 | ENSP00000448248 | 1006 | 9,94E-04 |

| | | | | | |
|---|---|---|---|---|---|
| GLMP | BALD | 1 | ENSP00000497576 | 1215 | 8,23E-04 |
| GOLGB1 | BALD | 2 | ENSP00000377275 | 9782 | 2,04E-04 |
| GOLPH3L | BALD | 1 | ENSP00000271732 | 854 | 1,17E-03 |
| GP2 | BALD | 1 | ENSP00000370767 | 1577 | 6,34E-04 |
| GPA33 | BALD | 1 | ENSP00000356842 | 941 | 1,06E-03 |
| GPATCH8 | BALD | 1 | ENSP00000467556 | 4394 | 2,28E-04 |
| GPR156 | BALD | 1 | ENSP00000417261 | 2412 | 4,15E-04 |
| GPR160 | BALD | 1 | ENSP00000348161 | 1016 | 9,84E-04 |
| GPR21 | BALD | 1 | ENSP00000362746 | 1049 | 9,53E-04 |
| GPRIN1 | BALD | 1 | ENSP00000305839 | 3015 | 3,32E-04 |
| GRK6 | BALD | 1 | ENSP00000433511 | 1635 | 6,12E-04 |
| GUF1 | BALD | 1 | ENSP00000281543 | 1993 | 5,02E-04 |
| GZMH | BALD | 1 | ENSP00000216338 | 736 | 1,36E-03 |
| HADH | BALD | 2 | ENSP00000425952 | 1227 | 1,63E-03 |
| HCLS1 | BALD | 1 | ENSP00000320176 | 1392 | 7,18E-04 |
| HEATR3 | BALD | 1 | ENSP00000299192 | 2028 | 4,93E-04 |
| HEG1 | BALD | 1 | ENSP00000311502 | 3847 | 2,60E-04 |
| HELLS | BALD | 1 | ENSP00000377601 | 2614 | 3,83E-04 |
| HLCS | BALD | 1 | ENSP00000338387 | 1705 | 5,87E-04 |
| HMCN1 | BALD | 1 | ENSP00000271588 | 16574 | 6,03E-05 |
| HMGCS1 | BALD | 1 | ENSP00000322706 | 1554 | 6,44E-04 |
| HOOK2 | BALD | 1 | ENSP00000380785 | 2132 | 4,69E-04 |
| HPX | BALD | 1 | ENSP00000265983 | 1376 | 7,27E-04 |
| HSDL2 | BALD | 1 | ENSP00000381785 | 1244 | 8,04E-04 |
| HSPA4L | BALD | 1 | ENSP00000422482 | 2504 | 3,99E-04 |
| ICAM1 | BALD | 1 | ENSP00000264832 | 1525 | 6,56E-04 |
| ICAM4 | BALD | 1 | ENSP00000342114 | 806 | 1,24E-03 |
| IFNB1 | BALD | 1 | ENSP00000369581 | 563 | 1,78E-03 |
| IGHE | BALD | 2 | ENSP00000492979 | 1428 | 1,40E-03 |
| IL10RA | BALD | 1 | ENSP00000227752 | 1657 | 6,04E-04 |
| IL12RB2 | BALD | 2 | ENSP00000262345 | 2568 | 7,79E-04 |
| IL15RA | BALD | 1 | ENSP00000380421 | 867 | 1,15E-03 |
| IL17RC | BALD | 1 | ENSP00000295981 | 2339 | 4,28E-04 |
| IL18 | BALD | 1 | ENSP00000280357 | 501 | 2,00E-03 |
| IL1RL2 | BALD | 1 | ENSP00000264257 | 1715 | 5,83E-04 |
| IL24 | BALD | 1 | ENSP00000375795 | 571 | 1,75E-03 |
| IL31 | BALD | 1 | ENSP00000366234 | 487 | 2,05E-03 |
| IL9 | BALD | 1 | ENSP00000274520 | 430 | 2,33E-03 |
| INHBE | BALD | 1 | ENSP00000266646 | 1051 | 9,51E-04 |
| INO80D | BALD | 1 | ENSP00000384198 | 3075 | 3,25E-04 |
| INTU | BALD | 1 | ENSP00000334003 | 2670 | 3,75E-04 |
| IQCA1 | BALD | 1 | ENSP00000407213 | 2463 | 4,06E-04 |
| IQCE | BALD | 1 | ENSP00000480715 | 2042 | 4,90E-04 |
| ITGA2 | BALD | 1 | ENSP00000296585 | 3451 | 2,90E-04 |
| ITGAL | BALD | 1 | ENSP00000349252 | 3503 | 2,85E-04 |
| ITGB3 | BALD | 1 | ENSP00000452786 | 2185 | 4,58E-04 |
| ITIH1 | BALD | 1 | ENSP00000273283 | 2687 | 3,72E-04 |
| IZUMO1 | BALD | 1 | ENSP00000327786 | 1090 | 9,17E-04 |

| | | | | | |
|---|---|---|---|---|---|
| JRK | BALD | 1 | ENSP00000485390 | 1661 | 6,02E-04 |
| KANSL1 | BALD | 1 | ENSP00000387393 | 3300 | 3,03E-04 |
| KAT2B | BALD | 2 | ENSP00000263754 | 2411 | 8,30E-04 |
| KDM3B | BALD | 1 | ENSP00000326563 | 5015 | 1,99E-04 |
| KHDC3L | BALD | 1 | ENSP00000359392 | 648 | 1,54E-03 |
| KIF18A | BALD | 1 | ENSP00000263181 | 2678 | 3,73E-04 |
| KIF24 | BALD | 2 | ENSP00000384433 | 4119 | 4,86E-04 |
| KLF15 | BALD | 1 | ENSP00000296233 | 1287 | 7,77E-04 |
| KLHL31 | BALD | 1 | ENSP00000384644 | 1903 | 5,25E-04 |
| KLRC2 | BALD | 1 | ENSP00000371327 | 1377 | 7,26E-04 |
| KNOP1 | BALD | 1 | ENSP00000219837 | 1379 | 7,25E-04 |
| KRT12 | BALD | 1 | ENSP00000251643 | 1480 | 6,76E-04 |
| KRT39 | BALD | 1 | ENSP00000347823 | 1469 | 6,81E-04 |
| KRT73 | BALD | 1 | ENSP00000307014 | 1614 | 6,20E-04 |
| KRT8 | BALD | 1 | ENSP00000449404 | 2980 | 3,36E-04 |
| LAG3 | BALD | 1 | ENSP00000203629 | 1570 | 6,37E-04 |
| LAMA2 | BALD | 1 | ENSP00000400365 | 9163 | 1,09E-04 |
| LAMB3 | BALD | 1 | ENSP00000348384 | 3496 | 2,86E-04 |
| LCN2 | BALD | 1 | ENSP00000362089 | 589 | 1,70E-03 |
| LETM1 | BALD | 1 | ENSP00000305653 | 2201 | 4,54E-04 |
| LIG4 | BALD | 1 | ENSP00000402030 | 2735 | 3,66E-04 |
| LIMCH1 | BALD | 2 | ENSP00000425631 | 4365 | 4,58E-04 |
| LIN28B | BALD | 1 | ENSP00000490468 | 792 | 1,26E-03 |
| LIN54 | BALD | 1 | ENSP00000341947 | 2235 | 4,47E-04 |
| LIPI | BALD | 1 | ENSP00000343331 | 1329 | 7,52E-04 |
| LPAR3 | BALD | 1 | ENSP00000395389 | 1060 | 9,43E-04 |
| LPCAT1 | BALD | 1 | ENSP00000423472 | 1587 | 6,30E-04 |
| LPCAT4 | BALD | 1 | ENSP00000317300 | 1557 | 6,42E-04 |
| LRIG2 | BALD | 1 | ENSP00000355396 | 3180 | 3,14E-04 |
| LRRC43 | BALD | 1 | ENSP00000344233 | 1861 | 5,37E-04 |
| LRRC49 | BALD | 1 | ENSP00000453273 | 2060 | 4,85E-04 |
| LRRC52 | BALD | 1 | ENSP00000294818 | 940 | 1,06E-03 |
| LRRC66 | BALD | 1 | ENSP00000341944 | 2629 | 3,80E-04 |
| LRRC8D | BALD | 1 | ENSP00000338887 | 2576 | 3,88E-04 |
| LRTM2 | BALD | 1 | ENSP00000299194 | 1238 | 8,08E-04 |
| LY75-CD302 | BALD | 3 | ENSP00000423463 | 5483 | 5,47E-04 |
| MAP2 | BALD | 2 | ENSP00000353508 | 5481 | 3,65E-04 |
| MAP3K19 | BALD | 1 | ENSP00000376647 | 3904 | 2,56E-04 |
| MCC | BALD | 1 | ENSP00000386227 | 2426 | 4,12E-04 |
| MCM2 | BALD | 1 | ENSP00000265056 | 2700 | 3,70E-04 |
| METTL18 | BALD | 1 | ENSP00000307077 | 1121 | 8,92E-04 |
| MFAP2 | BALD | 1 | ENSP00000364685 | 549 | 1,82E-03 |
| MFN1 | BALD | 2 | ENSP00000420617 | 2215 | 9,03E-04 |
| MFSD6 | BALD | 1 | ENSP00000376141 | 2360 | 4,24E-04 |
| MGAT4C | BALD | 1 | ENSP00000481096 | 1538 | 6,50E-04 |
| MIOX | BALD | 1 | ENSP00000216075 | 846 | 1,18E-03 |
| MIPOL1 | BALD | 1 | ENSP00000333539 | 1254 | 7,97E-04 |
| MKX | BALD | 1 | ENSP00000364946 | 1045 | 9,57E-04 |

| | | | | | |
|---|---|---|---|---|---|
| MLIP | BALD | 1 | ENSP00000426290 | 2952 | 3,39E-04 |
| MLN | BALD | 1 | ENSP00000388825 | 342 | 2,92E-03 |
| MMP27 | BALD | 1 | ENSP00000260229 | 1523 | 6,57E-04 |
| MRGPRD | BALD | 1 | ENSP00000310631 | 962 | 1,04E-03 |
| MRM3 | BALD | 1 | ENSP00000306080 | 1208 | 8,28E-04 |
| MROH2B | BALD | 1 | ENSP00000382476 | 4759 | 2,10E-04 |
| MRPL16 | BALD | 1 | ENSP00000300151 | 752 | 1,33E-03 |
| MRPL32 | BALD | 1 | ENSP00000223324 | 564 | 1,77E-03 |
| MRPS10 | BALD | 1 | ENSP00000053468 | 599 | 1,67E-03 |
| MSH4 | BALD | 1 | ENSP00000263187 | 2791 | 3,58E-04 |
| MSRB3 | BALD | 1 | ENSP00000347324 | 265 | 3,77E-03 |
| MTERF4 | BALD | 1 | ENSP00000241527 | 1092 | 9,16E-04 |
| MTFMT | BALD | 1 | ENSP00000220058 | 1153 | 8,67E-04 |
| MTG1 | BALD | 1 | ENSP00000323047 | 1008 | 9,92E-04 |
| MTRF1L | BALD | 1 | ENSP00000356202 | 1142 | 8,76E-04 |
| MTUS1 | BALD | 2 | ENSP00000262102 | 3505 | 5,71E-04 |
| MUC12 | BALD | 1 | ENSP00000368755 | 3006 | 3,33E-04 |
| MYBL2 | BALD | 1 | ENSP00000217026 | 2083 | 4,80E-04 |
| MYO1A | BALD | 1 | ENSP00000300119 | 3159 | 3,17E-04 |
| MYO3A | BALD | 1 | ENSP00000495965 | 4836 | 2,07E-04 |
| MYO5B | BALD | 1 | ENSP00000285039 | 5508 | 1,82E-04 |
| MYO9A | BALD | 1 | ENSP00000348349 | 7606 | 1,31E-04 |
| MYOM1 | BALD | 3 | ENSP00000348821 | 5024 | 5,97E-04 |
| NAA15 | BALD | 1 | ENSP00000296543 | 2525 | 3,96E-04 |
| NAT10 | BALD | 1 | ENSP00000257829 | 3056 | 3,27E-04 |
| NAV3 | BALD | 1 | ENSP00000381007 | 6856 | 1,46E-04 |
| NCKAP5 | BALD | 1 | ENSP00000387128 | 5658 | 1,77E-04 |
| NCOA7 | BALD | 1 | ENSP00000357341 | 2773 | 3,61E-04 |
| NDUFA2 | BALD | 1 | ENSP00000252102 | 412 | 2,43E-03 |
| NDUFAF8 | BALD | 1 | ENSP00000400184 | 195 | 5,13E-03 |
| NEK1 | BALD | 1 | ENSP00000424757 | 3852 | 2,60E-04 |
| NFATC3 | BALD | 1 | ENSP00000300659 | 3233 | 3,09E-04 |
| NIN | BALD | 2 | ENSP00000436092 | 6376 | 3,14E-04 |
| NIPBL | BALD | 1 | ENSP00000282516 | 8365 | 1,20E-04 |
| NKAPL | BALD | 2 | ENSP00000345716 | 1211 | 1,65E-03 |
| NLRP11 | BALD | 1 | ENSP00000466285 | 3057 | 3,27E-04 |
| NMRAL1 | BALD | 1 | ENSP00000283429 | 963 | 1,04E-03 |
| NOL6 | BALD | 1 | ENSP00000297990 | 3427 | 2,92E-04 |
| NOL8 | BALD | 1 | ENSP00000401177 | 3520 | 2,84E-04 |
| NOP16 | BALD | 1 | ENSP00000484969 | 532 | 1,88E-03 |
| NPVF | BALD | 1 | ENSP00000222674 | 588 | 1,70E-03 |
| NRIP1 | BALD | 1 | ENSP00000383060 | 3476 | 2,88E-04 |
| NTAN1 | BALD | 1 | ENSP00000287706 | 920 | 1,09E-03 |
| NTRK3 | BALD | 1 | ENSP00000486784 | 2559 | 3,91E-04 |
| NUP153 | BALD | 1 | ENSP00000444029 | 4391 | 2,28E-04 |
| OAS3 | BALD | 1 | ENSP00000228928 | 3239 | 3,09E-04 |
| OLR1 | BALD | 1 | ENSP00000309124 | 730 | 1,37E-03 |
| OR2S2 | BALD | 1 | ENSP00000344040 | 1081 | 9,25E-04 |

| | | | | | |
|---|---|---|---|---|---|
| OR6Q1 | BALD | 1 | ENSP00000307734 | 1105 | 9,05E-04 |
| OR8S1 | BALD | 1 | ENSP00000310632 | 1924 | 5,20E-04 |
| OVCA2 | BALD | 1 | ENSP00000461388 | 709 | 1,41E-03 |
| PALMD | BALD | 1 | ENSP00000263174 | 1646 | 6,08E-04 |
| PAPPA2 | BALD | 1 | ENSP00000356634 | 5354 | 1,87E-04 |
| AACS | BLACK | 1 | ENSP00000324842 | 1997 | 5,01E-04 |
| AASDH | BLACK | 1 | ENSP00000205214 | 3283 | 3,05E-04 |
| AATF | BLACK | 1 | ENSP00000477848 | 1896 | 5,27E-04 |
| ABCA1 | BLACK | 1 | ENSP00000363868 | 6762 | 1,48E-04 |
| ABCA13 | BLACK | 2 | ENSP00000411096 | 15112 | 1,32E-04 |
| ABLIM1 | BLACK | 1 | ENSP00000497150 | 2348 | 4,26E-04 |
| ACAT1 | BLACK | 1 | ENSP00000265838 | 1207 | 8,29E-04 |
| ACCS | BLACK | 1 | ENSP00000263776 | 1507 | 6,64E-04 |
| ACKR1 | BLACK | 1 | ENSP00000357103 | 1019 | 9,81E-04 |
| ACP5 | BLACK | 1 | ENSP00000496973 | 966 | 1,04E-03 |
| ACRBP | BLACK | 1 | ENSP00000229243 | 1619 | 6,18E-04 |
| ACSF2 | BLACK | 1 | ENSP00000401831 | 1918 | 5,21E-04 |
| ACSL5 | BLACK | 1 | ENSP00000348429 | 2173 | 4,60E-04 |
| ADAMTS3 | BLACK | 1 | ENSP00000286657 | 3603 | 2,78E-04 |
| ADCY10 | BLACK | 1 | ENSP00000356825 | 4901 | 2,04E-04 |
| ADCY3 | BLACK | 1 | ENSP00000384484 | 3403 | 2,94E-04 |
| ADGRG6 | BLACK | 1 | ENSP00000356581 | 3729 | 2,68E-04 |
| ADGRV1 | BLACK | 1 | ENSP00000384582 | 18059 | 5,54E-05 |
| ADH1B | BLACK | 1 | ENSP00000306606 | 1076 | 9,29E-04 |
| AFDN | BLACK | 1 | ENSP00000383623 | 5466 | 1,83E-04 |
| AGBL1 | BLACK | 1 | ENSP00000413001 | 3285 | 3,04E-04 |
| AHCTF1 | BLACK | 2 | ENSP00000355464 | 6738 | 2,97E-04 |
| AHR | BLACK | 1 | ENSP00000242057 | 2501 | 4,00E-04 |
| AIPL1 | BLACK | 1 | ENSP00000370521 | 1077 | 9,29E-04 |
| AK2 | BLACK | 1 | ENSP00000346921 | 710 | 1,41E-03 |
| AK8 | BLACK | 1 | ENSP00000298545 | 1427 | 7,01E-04 |
| AKAP9 | BLACK | 1 | ENSP00000351922 | 11676 | 8,56E-05 |
| ALCAM | BLACK | 1 | ENSP00000305988 | 1663 | 6,01E-04 |
| ALG1 | BLACK | 1 | ENSP00000262374 | 1377 | 7,26E-04 |
| ALG2 | BLACK | 1 | ENSP00000417764 | 1245 | 8,03E-04 |
| ANKRD12 | BLACK | 1 | ENSP00000262126 | 6168 | 1,62E-04 |
| ANKRD34C | BLACK | 1 | ENSP00000401089 | 1607 | 6,22E-04 |
| ANKZF1 | BLACK | 1 | ENSP00000321617 | 2162 | 4,63E-04 |
| ANO3 | BLACK | 2 | ENSP00000256737 | 2914 | 6,86E-04 |
| ANO5 | BLACK | 1 | ENSP00000315371 | 3215 | 3,11E-04 |
| AP4E1 | BLACK | 1 | ENSP00000261842 | 3399 | 2,94E-04 |
| APC | BLACK | 1 | ENSP00000257430 | 8517 | 1,17E-04 |
| ARFGAP1 | BLACK | 1 | ENSP00000314615 | 1202 | 8,32E-04 |
| ARFGAP2 | BLACK | 1 | ENSP00000434442 | 1553 | 6,44E-04 |
| ARG1 | BLACK | 1 | ENSP00000349446 | 961 | 1,04E-03 |
| ARHGAP24 | BLACK | 1 | ENSP00000378611 | 1982 | 5,05E-04 |
| ARHGAP29 | BLACK | 1 | ENSP00000260526 | 3797 | 2,63E-04 |
| ARHGAP31 | BLACK | 1 | ENSP00000264245 | 4328 | 2,31E-04 |

| | | | | | |
|---|---|---|---|---|---|
| ARHGEF28 | BLACK | 1 | ENSP00000411459 | 5175 | 1,93E-04 |
| ARID4A | BLACK | 1 | ENSP00000347602 | 3749 | 2,67E-04 |
| ARL6IP1 | BLACK | 1 | ENSP00000306788 | 301 | 3,32E-03 |
| ARMC2 | BLACK | 2 | ENSP00000376417 | 2587 | 7,73E-04 |
| ARMC7 | BLACK | 1 | ENSP00000245543 | 586 | 1,71E-03 |
| ASH1L | BLACK | 1 | ENSP00000357330 | 8871 | 1,13E-04 |
| ASTE1 | BLACK | 2 | ENSP00000426421 | 2107 | 9,49E-04 |
| ATP13A4 | BLACK | 1 | ENSP00000339182 | 3564 | 2,81E-04 |
| ATP6V0A2 | BLACK | 1 | ENSP00000332247 | 2551 | 3,92E-04 |
| ATP6V1E2 | BLACK | 1 | ENSP00000304891 | 680 | 1,47E-03 |
| ATP9B | BLACK | 1 | ENSP00000398076 | 3336 | 3,00E-04 |
| BAZ2B | BLACK | 1 | ENSP00000376534 | 6326 | 1,58E-04 |
| BBX | BLACK | 1 | ENSP00000319974 | 2817 | 3,55E-04 |
| BCAS3 | BLACK | 1 | ENSP00000468592 | 1970 | 5,08E-04 |
| BCL6B | BLACK | 1 | ENSP00000293805 | 1427 | 7,01E-04 |
| BDP1 | BLACK | 1 | ENSP00000351575 | 7502 | 1,33E-04 |
| BID | BLACK | 1 | ENSP00000318822 | 625 | 1,60E-03 |
| BOD1L1 | BLACK | 1 | ENSP00000040738 | 8946 | 1,12E-04 |
| BRD7 | BLACK | 1 | ENSP00000378181 | 1942 | 5,15E-04 |
| BRICD5 | BLACK | 1 | ENSP00000455052 | 753 | 1,33E-03 |
| BRWD1 | BLACK | 1 | ENSP00000330753 | 6877 | 1,45E-04 |
| BST1 | BLACK | 1 | ENSP00000371783 | 772 | 1,30E-03 |
| BTBD10 | BLACK | 1 | ENSP00000431186 | 1444 | 6,93E-04 |
| BTG2 | BLACK | 1 | ENSP00000433553 | 475 | 2,11E-03 |
| BTNL9 | BLACK | 1 | ENSP00000330200 | 1613 | 6,20E-04 |
| C12orf45 | BLACK | 1 | ENSP00000490540 | 817 | 1,22E-03 |
| C19orf44 | BLACK | 2 | ENSP00000221671 | 1967 | 1,02E-03 |
| C1QTNF6 | BLACK | 1 | ENSP00000380299 | 834 | 1,20E-03 |
| C3orf33 | BLACK | 1 | ENSP00000342512 | 880 | 1,14E-03 |
| C3orf62 | BLACK | 1 | ENSP00000341139 | 801 | 1,25E-03 |
| CA2 | BLACK | 1 | ENSP00000285379 | 786 | 1,27E-03 |
| CA3 | BLACK | 1 | ENSP00000285381 | 776 | 1,29E-03 |
| CAMKMT | BLACK | 1 | ENSP00000367755 | 801 | 1,25E-03 |
| CARD11 | BLACK | 1 | ENSP00000380150 | 3428 | 2,92E-04 |
| CARD14 | BLACK | 1 | ENSP00000499145 | 3241 | 3,09E-04 |
| CARS2 | BLACK | 1 | ENSP00000257347 | 1661 | 6,02E-04 |
| CASP2 | BLACK | 1 | ENSP00000312664 | 1274 | 7,85E-04 |
| CASR | BLACK | 1 | ENSP00000420194 | 3261 | 3,07E-04 |
| CBLN4 | BLACK | 1 | ENSP00000064571 | 599 | 1,67E-03 |
| CCDC148 | BLACK | 1 | ENSP00000386674 | 1800 | 5,56E-04 |
| CCDC153 | BLACK | 1 | ENSP00000423567 | 618 | 1,62E-03 |
| CCDC168 | BLACK | 1 | ENSP00000320232 | 20045 | 4,99E-05 |
| CCDC181 | BLACK | 1 | ENSP00000356780 | 1516 | 6,60E-04 |
| CCDC190 | BLACK | 1 | ENSP00000356886 | 906 | 1,10E-03 |
| CCDC51 | BLACK | 1 | ENSP00000379047 | 1233 | 8,11E-04 |
| CCDC63 | BLACK | 1 | ENSP00000312399 | 1678 | 5,96E-04 |
| CCL2 | BLACK | 1 | ENSP00000225831 | 297 | 3,37E-03 |
| CCNA1 | BLACK | 1 | ENSP00000255465 | 1332 | 7,51E-04 |

| | | | | | |
|---|---|---|---|---|---|
| CCNDBP1 | BLACK | 1 | ENSP00000300213 | 1082 | 9,24E-04 |
| CCR6 | BLACK | 2 | ENSP00000493637 | 959 | 2,09E-03 |
| CD101 | BLACK | 1 | ENSP00000358482 | 3096 | 3,23E-04 |
| CD163L1 | BLACK | 1 | ENSP00000393474 | 4374 | 2,29E-04 |
| CD22 | BLACK | 1 | ENSP00000085219 | 2526 | 3,96E-04 |
| CD38 | BLACK | 1 | ENSP00000226279 | 857 | 1,17E-03 |
| CD79B | BLACK | 1 | ENSP00000376544 | 693 | 1,44E-03 |
| CD8A | BLACK | 1 | ENSP00000283635 | 702 | 1,42E-03 |
| CD96 | BLACK | 2 | ENSP00000283285 | 1696 | 1,18E-03 |
| CDC16 | BLACK | 1 | ENSP00000348554 | 1845 | 5,42E-04 |
| CDC25C | BLACK | 1 | ENSP00000321656 | 1513 | 6,61E-04 |
| CDCA7 | BLACK | 1 | ENSP00000306968 | 1064 | 9,40E-04 |
| CDCP1 | BLACK | 2 | ENSP00000296129 | 2422 | 8,26E-04 |
| CDHR2 | BLACK | 1 | ENSP00000424565 | 3500 | 2,86E-04 |
| CDON | BLACK | 2 | ENSP00000376458 | 3848 | 5,20E-04 |
| CDV3 | BLACK | 1 | ENSP00000264993 | 769 | 1,30E-03 |
| CEACAM6 | BLACK | 1 | ENSP00000199764 | 956 | 1,05E-03 |
| CELA2A | BLACK | 1 | ENSP00000352639 | 789 | 1,27E-03 |
| CENPE | BLACK | 1 | ENSP00000265148 | 7999 | 1,25E-04 |
| CENPF | BLACK | 1 | ENSP00000355922 | 9329 | 1,07E-04 |
| CENPN | BLACK | 1 | ENSP00000377007 | 1049 | 9,53E-04 |
| CENPW | BLACK | 1 | ENSP00000357308 | 331 | 3,02E-03 |
| CEP120 | BLACK | 1 | ENSP00000303058 | 2990 | 3,34E-04 |
| CEP126 | BLACK | 1 | ENSP00000263468 | 3316 | 3,02E-04 |
| CEP152 | BLACK | 1 | ENSP00000370337 | 5107 | 1,96E-04 |
| CEP164 | BLACK | 2 | ENSP00000278935 | 4263 | 4,69E-04 |
| CEP290 | BLACK | 1 | ENSP00000308021 | 7376 | 1,36E-04 |
| CEP350 | BLACK | 1 | ENSP00000356579 | 9317 | 1,07E-04 |
| CEP89 | BLACK | 1 | ENSP00000306105 | 2338 | 4,28E-04 |
| CFAP43 | BLACK | 1 | ENSP00000349568 | 5029 | 1,99E-04 |
| CFAP54 | BLACK | 1 | ENSP00000431759 | 9205 | 1,09E-04 |
| CGNL1 | BLACK | 1 | ENSP00000281282 | 3851 | 2,60E-04 |
| CHGB | BLACK | 1 | ENSP00000368244 | 1973 | 5,07E-04 |
| CHRNA3 | BLACK | 1 | ENSP00000452896 | 1437 | 6,96E-04 |
| CIITA | BLACK | 1 | ENSP00000485010 | 3335 | 3,00E-04 |
| CILP | BLACK | 1 | ENSP00000261883 | 3547 | 2,82E-04 |
| CLEC17A | BLACK | 1 | ENSP00000393719 | 1116 | 8,96E-04 |
| CLEC6A | BLACK | 1 | ENSP00000371505 | 595 | 1,68E-03 |
| CLSPN | BLACK | 1 | ENSP00000312995 | 3954 | 2,53E-04 |
| CMA1 | BLACK | 1 | ENSP00000250378 | 739 | 1,35E-03 |
| CNGB3 | BLACK | 1 | ENSP00000316605 | 2379 | 4,20E-04 |
| CNN2 | BLACK | 1 | ENSP00000456436 | 874 | 1,14E-03 |
| CNTN1 | BLACK | 1 | ENSP00000447006 | 2875 | 3,48E-04 |
| CNTRL | BLACK | 2 | ENSP00000362962 | 6964 | 2,87E-04 |
| COL15A1 | BLACK | 2 | ENSP00000364140 | 4094 | 4,89E-04 |
| COL16A1 | BLACK | 1 | ENSP00000362776 | 4672 | 2,14E-04 |
| COL6A3 | BLACK | 1 | ENSP00000295550 | 9630 | 1,04E-04 |
| COL8A1 | BLACK | 1 | ENSP00000261037 | 2236 | 4,47E-04 |

| | | | | | |
|---|---|---|---|---|---|
| COL9A1 | BLACK | 1 | ENSP00000349790 | 2711 | 3,69E-04 |
| COMT | BLACK | 1 | ENSP00000354511 | 809 | 1,24E-03 |
| COQ7 | BLACK | 1 | ENSP00000322316 | 648 | 1,54E-03 |
| CPA1 | BLACK | 1 | ENSP00000011292 | 1250 | 8,00E-04 |
| CPAMD8 | BLACK | 1 | ENSP00000498697 | 5869 | 1,70E-04 |
| CRACR2A | BLACK | 1 | ENSP00000409382 | 2182 | 4,58E-04 |
| CRISP1 | BLACK | 1 | ENSP00000338276 | 678 | 1,47E-03 |
| CRYBG3 | BLACK | 1 | ENSP00000374273 | 8821 | 1,13E-04 |
| CSF1 | BLACK | 1 | ENSP00000327513 | 1599 | 6,25E-04 |
| CX3CL1 | BLACK | 1 | ENSP00000456830 | 1139 | 8,78E-04 |
| CYP2B6 | BLACK | 1 | ENSP00000324648 | 1297 | 7,71E-04 |
| CYP4V2 | BLACK | 1 | ENSP00000368079 | 1567 | 6,38E-04 |
| CYP7A1 | BLACK | 1 | ENSP00000301645 | 1498 | 6,68E-04 |
| DAPK1 | BLACK | 1 | ENSP00000350785 | 4204 | 2,38E-04 |
| DBX1 | BLACK | 1 | ENSP00000436881 | 1033 | 9,68E-04 |
| DCAF4L2 | BLACK | 2 | ENSP00000316496 | 1187 | 1,68E-03 |
| DCDC2 | BLACK | 1 | ENSP00000367715 | 1420 | 7,04E-04 |
| DCDC2B | BLACK | 1 | ENSP00000386870 | 1038 | 9,63E-04 |
| DCTN3 | BLACK | 1 | ENSP00000259632 | 554 | 1,81E-03 |
| DDIAS | BLACK | 1 | ENSP00000435421 | 3002 | 3,33E-04 |
| DDIT3 | BLACK | 1 | ENSP00000448665 | 577 | 1,73E-03 |
| DDX20 | BLACK | 1 | ENSP00000358716 | 2469 | 4,05E-04 |
| DDX21 | BLACK | 2 | ENSP00000346120 | 2337 | 8,56E-04 |
| DDX31 | BLACK | 1 | ENSP00000361232 | 2523 | 3,96E-04 |
| DHODH | BLACK | 1 | ENSP00000219240 | 1141 | 8,76E-04 |
| DHRS2 | BLACK | 1 | ENSP00000344674 | 917 | 1,09E-03 |
| DISC1 | BLACK | 1 | ENSP00000403888 | 2539 | 3,94E-04 |
| DLGAP5 | BLACK | 1 | ENSP00000247191 | 2520 | 3,97E-04 |
| DNAH10 | BLACK | 2 | ENSP00000386770 | 13529 | 1,48E-04 |
| DNAH3 | BLACK | 1 | ENSP00000261383 | 12283 | 8,14E-05 |
| DNAH7 | BLACK | 1 | ENSP00000311273 | 11936 | 8,38E-05 |
| DNAJC6 | BLACK | 1 | ENSP00000360108 | 2894 | 3,46E-04 |
| DNMBP | BLACK | 1 | ENSP00000315659 | 4692 | 2,13E-04 |
| DOCK10 | BLACK | 1 | ENSP00000493664 | 6383 | 1,57E-04 |
| DOK5 | BLACK | 1 | ENSP00000262593 | 848 | 1,18E-03 |
| DONSON | BLACK | 1 | ENSP00000307143 | 1674 | 5,97E-04 |
| DPEP3 | BLACK | 1 | ENSP00000268793 | 1519 | 6,58E-04 |
| DSC2 | BLACK | 1 | ENSP00000280904 | 2616 | 3,82E-04 |
| DSC3 | BLACK | 1 | ENSP00000353608 | 2675 | 3,74E-04 |
| DYTN | BLACK | 1 | ENSP00000396593 | 1720 | 5,81E-04 |
| E2F8 | BLACK | 1 | ENSP00000434199 | 2581 | 3,87E-04 |
| ECI2 | BLACK | 1 | ENSP00000369461 | 1176 | 8,50E-04 |
| ECT2L | BLACK | 1 | ENSP00000387388 | 2695 | 3,71E-04 |
| EEF2K | BLACK | 1 | ENSP00000263026 | 2161 | 4,63E-04 |
| EFCAB12 | BLACK | 1 | ENSP00000420854 | 1721 | 5,81E-04 |
| EIF2B4 | BLACK | 2 | ENSP00000429323 | 1628 | 1,23E-03 |
| EIF3F | BLACK | 1 | ENSP00000431800 | 1076 | 9,29E-04 |
| EME2 | BLACK | 1 | ENSP00000457353 | 1110 | 9,01E-04 |

| | | | | | |
|---|---|---|---|---|---|
| ENAM | BLACK | 1 | ENSP00000379383 | 3436 | 2,91E-04 |
| ENO4 | BLACK | 1 | ENSP00000482973 | 1902 | 5,26E-04 |
| EP300 | BLACK | 1 | ENSP00000263253 | 7145 | 1,40E-04 |
| EPX | BLACK | 1 | ENSP00000225371 | 2129 | 4,70E-04 |
| ERAP2 | BLACK | 1 | ENSP00000400376 | 2846 | 3,51E-04 |
| ERCC6L2 | BLACK | 1 | ENSP00000499221 | 4569 | 2,19E-04 |
| ERFE | BLACK | 2 | ENSP00000442304 | 1073 | 1,86E-03 |
| ESPL1 | BLACK | 1 | ENSP00000449831 | 6323 | 1,58E-04 |
| ETNPPL | BLACK | 1 | ENSP00000296486 | 1428 | 7,00E-04 |
| ETV1 | BLACK | 1 | ENSP00000384085 | 1491 | 6,71E-04 |
| EVC | BLACK | 1 | ENSP00000264956 | 2680 | 3,73E-04 |
| EXOC3 | BLACK | 1 | ENSP00000425587 | 2210 | 4,52E-04 |
| EXTL1 | BLACK | 1 | ENSP00000363398 | 2009 | 4,98E-04 |
| EYA4 | BLACK | 1 | ENSP00000432770 | 1843 | 5,43E-04 |
| F2RL3 | BLACK | 1 | ENSP00000248076 | 1148 | 8,71E-04 |
| FAM104A | BLACK | 1 | ENSP00000384832 | 614 | 1,63E-03 |
| FAM160A2 | BLACK | 1 | ENSP00000265978 | 2942 | 3,40E-04 |
| FAM161B | BLACK | 1 | ENSP00000499021 | 1890 | 5,29E-04 |
| FAM171A1 | BLACK | 1 | ENSP00000367356 | 2563 | 3,90E-04 |
| FAM172A | BLACK | 1 | ENSP00000379294 | 1125 | 8,89E-04 |
| FAM184A | BLACK | 1 | ENSP00000342604 | 3387 | 2,95E-04 |
| FAM217B | BLACK | 1 | ENSP00000351040 | 1163 | 8,60E-04 |
| FANCC | BLACK | 1 | ENSP00000289081 | 1663 | 6,01E-04 |
| FANCD2 | BLACK | 1 | ENSP00000287647 | 4367 | 2,29E-04 |
| FANCF | BLACK | 3 | ENSP00000330875 | 1124 | 2,67E-03 |
| FARP2 | BLACK | 1 | ENSP00000264042 | 3027 | 3,30E-04 |
| FASTK | BLACK | 1 | ENSP00000297532 | 1626 | 6,15E-04 |
| FAT1 | BLACK | 3 | ENSP00000479573 | 13741 | 2,18E-04 |
| FBXO6 | BLACK | 1 | ENSP00000365944 | 866 | 1,15E-03 |
| FBXW9 | BLACK | 1 | ENSP00000465387 | 1249 | 8,01E-04 |
| FCN3 | BLACK | 1 | ENSP00000270879 | 899 | 1,11E-03 |
| FCRL5 | BLACK | 5 | ENSP00000354691 | 2863 | 1,75E-03 |
| FMO1 | BLACK | 1 | ENSP00000481732 | 1715 | 5,83E-04 |
| FN1 | BLACK | 1 | ENSP00000346839 | 7387 | 1,35E-04 |
| FNDC3A | BLACK | 1 | ENSP00000417257 | 3572 | 2,80E-04 |
| FOLR1 | BLACK | 1 | ENSP00000308137 | 764 | 1,31E-03 |
| FOXI1 | BLACK | 1 | ENSP00000304286 | 1127 | 8,87E-04 |
| FREM3 | BLACK | 1 | ENSP00000332886 | 6415 | 1,56E-04 |
| FSIP2 | BLACK | 4 | ENSP00000401306 | 20633 | 1,94E-04 |
| GAL3ST1 | BLACK | 1 | ENSP00000385825 | 1266 | 7,90E-04 |
| GALR2 | BLACK | 1 | ENSP00000329684 | 1154 | 8,67E-04 |
| GATB | BLACK | 1 | ENSP00000263985 | 1661 | 6,02E-04 |
| GBP6 | BLACK | 2 | ENSP00000359485 | 3775 | 5,30E-04 |
| GCNT1 | BLACK | 1 | ENSP00000365920 | 1286 | 7,78E-04 |
| GDF2 | BLACK | 1 | ENSP00000463051 | 1288 | 7,76E-04 |
| GEMIN5 | BLACK | 1 | ENSP00000285873 | 4496 | 2,22E-04 |
| GHR | BLACK | 2 | ENSP00000483403 | 1926 | 1,04E-03 |
| GIN1 | BLACK | 1 | ENSP00000381970 | 1562 | 6,40E-04 |

| | | | | | |
|---|---|---|---|---|---|
| GIPC2 | BLACK | 1 | ENSP00000359795 | 942 | 1,06E-03 |
| GLB1 | BLACK | 1 | ENSP00000306920 | 1934 | 5,17E-04 |
| GLB1L3 | BLACK | 1 | ENSP00000396615 | 1934 | 5,17E-04 |
| GLCCI1 | BLACK | 1 | ENSP00000223145 | 1622 | 6,17E-04 |
| GLE1 | BLACK | 1 | ENSP00000308622 | 2077 | 4,81E-04 |
| GNAS | BLACK | 1 | ENSP00000360141 | 3583 | 2,79E-04 |
| GOLGA1 | BLACK | 1 | ENSP00000362656 | 2278 | 4,39E-04 |
| GPC2 | BLACK | 1 | ENSP00000292377 | 1746 | 5,73E-04 |
| GPR152 | BLACK | 1 | ENSP00000310255 | 1427 | 7,01E-04 |
| GPR157 | BLACK | 1 | ENSP00000366628 | 994 | 1,01E-03 |
| GPR158 | BLACK | 1 | ENSP00000365529 | 2625 | 3,81E-04 |
| GRHPR | BLACK | 1 | ENSP00000475569 | 970 | 1,03E-03 |
| GSAP | BLACK | 1 | ENSP00000257626 | 2427 | 4,12E-04 |
| GSDMD | BLACK | 1 | ENSP00000433958 | 1460 | 6,85E-04 |
| GTPBP4 | BLACK | 1 | ENSP00000354040 | 1847 | 5,41E-04 |
| GUCY2D | BLACK | 1 | ENSP00000254854 | 3290 | 3,04E-04 |
| HACD4 | BLACK | 1 | ENSP00000419503 | 678 | 1,47E-03 |
| HAO1 | BLACK | 1 | ENSP00000368066 | 1105 | 9,05E-04 |
| HBQ1 | BLACK | 1 | ENSP00000199708 | 416 | 2,40E-03 |
| HEATR5A | BLACK | 2 | ENSP00000437968 | 6106 | 3,28E-04 |
| HELQ | BLACK | 1 | ENSP00000295488 | 3288 | 3,04E-04 |
| HERC1 | BLACK | 1 | ENSP00000390158 | 12848 | 7,78E-05 |
| HEYL | BLACK | 1 | ENSP00000361943 | 978 | 1,02E-03 |
| HFE | BLACK | 1 | ENSP00000417404 | 1009 | 9,91E-04 |
| HHIPL2 | BLACK | 1 | ENSP00000342118 | 2149 | 4,65E-04 |
| HINFP | BLACK | 1 | ENSP00000318085 | 1542 | 6,49E-04 |
| HJURP | BLACK | 1 | ENSP00000414109 | 2179 | 4,59E-04 |
| HKDC1 | BLACK | 1 | ENSP00000346643 | 2666 | 3,75E-04 |
| HMGCR | BLACK | 1 | ENSP00000426745 | 2598 | 3,85E-04 |
| HOOK1 | BLACK | 1 | ENSP00000360252 | 2100 | 4,76E-04 |
| HPS1 | BLACK | 1 | ENSP00000355310 | 2129 | 4,70E-04 |
| HRH2 | BLACK | 1 | ENSP00000489742 | 1267 | 7,89E-04 |
| HTRA4 | BLACK | 1 | ENSP00000305919 | 1445 | 6,92E-04 |
| HYDIN | BLACK | 2 | ENSP00000377197 | 15379 | 1,30E-04 |
| IARS2 | BLACK | 1 | ENSP00000355889 | 3008 | 3,32E-04 |
| IBTK | BLACK | 1 | ENSP00000305721 | 4034 | 2,48E-04 |
| ICA1L | BLACK | 1 | ENSP00000376070 | 1434 | 6,97E-04 |
| IFT172 | BLACK | 1 | ENSP00000260570 | 5230 | 1,91E-04 |
| IGKV6D-41 | BLACK | 1 | ENSP00000374806 | 343 | 2,92E-03 |
| IGSF6 | BLACK | 1 | ENSP00000268389 | 652 | 1,53E-03 |
| IL16 | BLACK | 2 | ENSP00000302935 | 3962 | 5,05E-04 |
| IL19 | BLACK | 1 | ENSP00000499459 | 529 | 1,89E-03 |
| IL1RN | BLACK | 1 | ENSP00000259206 | 413 | 2,42E-03 |
| IL20RB | BLACK | 1 | ENSP00000328133 | 873 | 1,15E-03 |
| IL27RA | BLACK | 1 | ENSP00000263379 | 1883 | 5,31E-04 |
| IL2RA | BLACK | 1 | ENSP00000369293 | 729 | 1,37E-03 |
| IL6 | BLACK | 1 | ENSP00000385675 | 638 | 1,57E-03 |
| IL7R | BLACK | 1 | ENSP00000306157 | 1372 | 7,29E-04 |

| | | | | | |
|---|---|---|---|---|---|
| ILDR1 | BLACK | 1 | ENSP00000345667 | 1623 | 6,16E-04 |
| INPP5F | BLACK | 1 | ENSP00000497527 | 3382 | 2,96E-04 |
| INVS | BLACK | 1 | ENSP00000262457 | 3176 | 3,15E-04 |
| IPP | BLACK | 1 | ENSP00000379739 | 1747 | 5,72E-04 |
| IRAK1BP1 | BLACK | 1 | ENSP00000358956 | 779 | 1,28E-03 |
| IRAK3 | BLACK | 1 | ENSP00000261233 | 1773 | 5,64E-04 |
| ISG20L2 | BLACK | 1 | ENSP00000323424 | 1059 | 9,44E-04 |
| ITGA10 | BLACK | 1 | ENSP00000358310 | 3474 | 2,88E-04 |
| ITGAM | BLACK | 1 | ENSP00000496959 | 3573 | 2,80E-04 |
| ITGAV | BLACK | 1 | ENSP00000261023 | 3103 | 3,22E-04 |
| ITGB7 | BLACK | 1 | ENSP00000267082 | 2383 | 4,20E-04 |
| ITK | BLACK | 1 | ENSP00000398655 | 1846 | 5,42E-04 |
| IYD | BLACK | 1 | ENSP00000229447 | 865 | 1,16E-03 |
| KAT6A | BLACK | 1 | ENSP00000385888 | 5995 | 1,67E-04 |
| KCNN4 | BLACK | 1 | ENSP00000496939 | 1299 | 7,70E-04 |
| KHDRBS3 | BLACK | 1 | ENSP00000348108 | 1032 | 9,69E-04 |
| KIAA0319L | BLACK | 1 | ENSP00000318406 | 3156 | 3,17E-04 |
| KIAA0408 | BLACK | 1 | ENSP00000435150 | 2071 | 4,83E-04 |
| KIAA0586 | BLACK | 1 | ENSP00000346359 | 4872 | 2,05E-04 |
| KIAA1324 | BLACK | 1 | ENSP00000358955 | 3020 | 3,31E-04 |
| KIAA2012 | BLACK | 2 | ENSP00000419834 | 3527 | 5,67E-04 |
| KIF14 | BLACK | 2 | ENSP00000356319 | 4963 | 4,03E-04 |
| KLF12 | BLACK | 1 | ENSP00000366897 | 902 | 1,11E-03 |
| KLF17 | BLACK | 1 | ENSP00000361373 | 1081 | 9,25E-04 |
| KLHL38 | BLACK | 1 | ENSP00000321475 | 1743 | 5,74E-04 |
| KMO | BLACK | 1 | ENSP00000355517 | 1442 | 6,93E-04 |
| KRT36 | BLACK | 1 | ENSP00000329165 | 1675 | 5,97E-04 |
| KRTAP13-2 | BLACK | 1 | ENSP00000382777 | 527 | 1,90E-03 |
| KTN1 | BLACK | 1 | ENSP00000378725 | 4009 | 2,49E-04 |
| L3MBTL3 | BLACK | 1 | ENSP00000431962 | 2322 | 4,31E-04 |
| L3MBTL4 | BLACK | 1 | ENSP00000382976 | 1305 | 7,66E-04 |
| LAMC2 | BLACK | 1 | ENSP00000264144 | 3479 | 2,87E-04 |
| LARP4 | BLACK | 1 | ENSP00000415464 | 2157 | 4,64E-04 |
| LARP7 | BLACK | 1 | ENSP00000422626 | 1675 | 5,97E-04 |
| LCORL | BLACK | 3 | ENSP00000490600 | 5611 | 5,35E-04 |
| LGALS2 | BLACK | 1 | ENSP00000215886 | 426 | 2,35E-03 |
| LIFR | BLACK | 2 | ENSP00000263409 | 3275 | 6,11E-04 |
| LIMA1 | BLACK | 1 | ENSP00000378400 | 2276 | 4,39E-04 |
| LMAN1L | BLACK | 1 | ENSP00000310431 | 1580 | 6,33E-04 |
| LMBR1 | BLACK | 1 | ENSP00000326604 | 1457 | 6,86E-04 |
| LMO7 | BLACK | 1 | ENSP00000349571 | 4832 | 2,07E-04 |
| LMTK2 | BLACK | 1 | ENSP00000297293 | 4373 | 2,29E-04 |
| LPIN3 | BLACK | 1 | ENSP00000487971 | 2526 | 3,96E-04 |
| LRIG3 | BLACK | 1 | ENSP00000326759 | 3340 | 2,99E-04 |
| LRMP | BLACK | 1 | ENSP00000489956 | 4317 | 2,32E-04 |
| LRPPRC | BLACK | 1 | ENSP00000260665 | 4203 | 2,38E-04 |
| LRSAM1 | BLACK | 1 | ENSP00000322937 | 2153 | 4,64E-04 |
| LSG1 | BLACK | 1 | ENSP00000265245 | 2008 | 4,98E-04 |

| | | | | | |
|---|---|---|---|---|---|
| LYSMD3 | BLACK | 1 | ENSP00000314518 | 919 | 1,09E-03 |
| MALRD1 | BLACK | 3 | ENSP00000412763 | 6446 | 4,65E-04 |
| MAML3 | BLACK | 1 | ENSP00000421180 | 3219 | 3,11E-04 |
| MAN1A2 | BLACK | 1 | ENSP00000348959 | 1905 | 5,25E-04 |
| MANSC1 | BLACK | 1 | ENSP00000438205 | 1296 | 7,72E-04 |
| MASP2 | BLACK | 1 | ENSP00000383690 | 2034 | 4,92E-04 |
| MCM3AP | BLACK | 3 | ENSP00000380820 | 5915 | 5,07E-04 |
| MECOM | BLACK | 1 | ENSP00000498411 | 3648 | 2,74E-04 |
| MEIOB | BLACK | 1 | ENSP00000314484 | 1288 | 7,76E-04 |
| METTL4 | BLACK | 1 | ENSP00000458290 | 1411 | 7,09E-04 |
| METTL5 | BLACK | 1 | ENSP00000387056 | 725 | 1,38E-03 |
| MFAP5 | BLACK | 1 | ENSP00000352455 | 276 | 3,62E-03 |
| MFN2 | BLACK | 1 | ENSP00000235329 | 2260 | 4,42E-04 |
| MIS18BP1 | BLACK | 2 | ENSP00000309790 | 3380 | 5,92E-04 |
| MLH1 | BLACK | 1 | ENSP00000231790 | 2259 | 4,43E-04 |
| MLH3 | BLACK | 1 | ENSP00000452316 | 4387 | 2,28E-04 |
| MLPH | BLACK | 3 | ENSP00000264605 | 1756 | 1,71E-03 |
| MLXIP | BLACK | 1 | ENSP00000312834 | 2735 | 3,66E-04 |
| MMS19 | BLACK | 1 | ENSP00000412698 | 3039 | 3,29E-04 |
| MOAP1 | BLACK | 1 | ENSP00000298894 | 1058 | 9,45E-04 |
| MPHOSPH9 | BLACK | 1 | ENSP00000475489 | 3425 | 2,92E-04 |
| MPV17L | BLACK | 1 | ENSP00000379669 | 574 | 1,74E-03 |
| MPZL1 | BLACK | 1 | ENSP00000352513 | 804 | 1,24E-03 |
| MPZL3 | BLACK | 1 | ENSP00000278949 | 629 | 1,59E-03 |
| MRGPRX1 | BLACK | 2 | ENSP00000305766 | 1420 | 1,41E-03 |
| MRPL55 | BLACK | 1 | ENSP00000355699 | 382 | 2,62E-03 |
| MRPS27 | BLACK | 1 | ENSP00000426941 | 1235 | 8,10E-04 |
| MS4A18 | BLACK | 1 | ENSP00000431607 | 1191 | 8,40E-04 |
| MTIF2 | BLACK | 1 | ENSP00000384481 | 2171 | 4,61E-04 |
| MTMR11 | BLACK | 1 | ENSP00000391668 | 2127 | 4,70E-04 |
| MTSS1 | BLACK | 1 | ENSP00000322804 | 2057 | 4,86E-04 |
| MTUS2 | BLACK | 1 | ENSP00000498251 | 2938 | 3,40E-04 |
| MUC13 | BLACK | 1 | ENSP00000485028 | 1472 | 6,79E-04 |
| MUSK | BLACK | 1 | ENSP00000363571 | 2507 | 3,99E-04 |
| MYBPC2 | BLACK | 1 | ENSP00000350332 | 3299 | 3,03E-04 |
| MYLK | BLACK | 1 | ENSP00000418335 | 5753 | 1,74E-04 |
| MYO19 | BLACK | 1 | ENSP00000479518 | 2882 | 3,47E-04 |
| MYOC | BLACK | 1 | ENSP00000037502 | 1505 | 6,64E-04 |
| MYOD1 | BLACK | 1 | ENSP00000250003 | 953 | 1,05E-03 |
| MYOM2 | BLACK | 1 | ENSP00000262113 | 4358 | 2,29E-04 |
| MZB1 | BLACK | 1 | ENSP00000303920 | 569 | 1,76E-03 |
| N4BP2L1 | BLACK | 1 | ENSP00000369476 | 717 | 1,39E-03 |
| NAA11 | BLACK | 1 | ENSP00000286794 | 689 | 1,45E-03 |
| NAAA | BLACK | 1 | ENSP00000286733 | 1064 | 9,40E-04 |
| NAF1 | BLACK | 1 | ENSP00000274054 | 1479 | 6,76E-04 |
| NAGS | BLACK | 1 | ENSP00000293404 | 1590 | 6,29E-04 |
| NBN | BLACK | 2 | ENSP00000265433 | 2188 | 9,14E-04 |
| NCAPH2 | BLACK | 1 | ENSP00000299821 | 1918 | 5,21E-04 |

| | | | | | |
|---|---|---|---|---|---|
| NCL | BLACK | 1 | ENSP00000318195 | 1772 | 5,64E-04 |
| NDUFA10 | BLACK | 1 | ENSP00000302321 | 1300 | 7,69E-04 |
| NDUFA6 | BLACK | 2 | ENSP00000482543 | 459 | 4,36E-03 |
| NELL2 | BLACK | 1 | ENSP00000416341 | 2553 | 3,92E-04 |
| NID2 | BLACK | 1 | ENSP00000216286 | 4203 | 2,38E-04 |
| NIF3L1 | BLACK | 1 | ENSP00000498853 | 1128 | 8,87E-04 |
| NKG7 | BLACK | 1 | ENSP00000221978 | 494 | 2,02E-03 |
| NOB1 | BLACK | 1 | ENSP00000268802 | 1233 | 8,11E-04 |
| NOL9 | BLACK | 1 | ENSP00000366934 | 2085 | 4,80E-04 |
| NOSTRIN | BLACK | 1 | ENSP00000402140 | 1692 | 5,91E-04 |
| NPHP4 | BLACK | 2 | ENSP00000367398 | 4244 | 4,71E-04 |
| NPHS1 | BLACK | 1 | ENSP00000368190 | 2539 | 3,94E-04 |
| NPY | BLACK | 1 | ENSP00000242152 | 289 | 3,46E-03 |
| NRDE2 | BLACK | 1 | ENSP00000346335 | 3469 | 2,88E-04 |
| NRP1 | BLACK | 1 | ENSP00000265371 | 2693 | 3,71E-04 |
| NUDT7 | BLACK | 1 | ENSP00000268533 | 678 | 1,47E-03 |
| ODF2L | BLACK | 1 | ENSP00000359600 | 1746 | 5,73E-04 |
| OMA1 | BLACK | 1 | ENSP00000360270 | 1567 | 6,38E-04 |
| OR4C16 | BLACK | 1 | ENSP00000485295 | 428 | 2,34E-03 |
| OR9I1 | BLACK | 1 | ENSP00000493370 | 1003 | 9,97E-04 |
| ORC1 | BLACK | 1 | ENSP00000360623 | 2561 | 3,90E-04 |
| OSBPL6 | BLACK | 1 | ENSP00000376293 | 2755 | 3,63E-04 |
| P4HA2 | BLACK | 1 | ENSP00000384999 | 1594 | 6,27E-04 |
| P4HA3 | BLACK | 1 | ENSP00000401749 | 1855 | 5,39E-04 |
| PABPC3 | BLACK | 1 | ENSP00000281589 | 1904 | 5,25E-04 |
| PALB2 | BLACK | 1 | ENSP00000261584 | 3545 | 2,82E-04 |
| PANX3 | BLACK | 1 | ENSP00000284288 | 1175 | 8,51E-04 |
